# Temperature-Dependent Ion Migration Underlies Sequence-Specific Collapse of Unstructured RNA

**DOI:** 10.1101/2025.10.20.683600

**Authors:** Heyang Zhang, Hiranmay Maity, Hung T. Nguyen

## Abstract

Ions and temperature jointly regulate RNA structure, dynamics and phase behavior, yet their coupled effects remain poorly understood at the molecular level. Single-stranded RNA (ssRNA), a ubiquitous and functionally versatile class of RNA, presents a particularly challenging target due to its intrinsic flexibility and pronounced sensitivity to ionic and thermal perturbations. Here, we extend our previously validated coarse-grained RNA model by introducing temperature-dependent divalent ion-phosphate potentials along with revised stacking interactions to elucidate how electrostatics, stacking, and hydration collectively determine ssRNA behavior. Our simulations quantitatively reproduce experimental SAXS profiles across a broad range of ionic conditions and reveal a non-monotonic temperature dependence of RNA compaction: ssRNAs expand upon heating, reach a sequence-specific maximum size, and then collapse as enhanced counterion condensation dominates. Rising temperature strengthens ion-RNA interactions, leading to a reorganization from diffusive to inner-sphere coordination, directly linking RNA collapse to ion dehydration. Our results establish that the ion atmosphere is a dynamic, sequence-encoded extension of RNA structure. This framework provides molecular insight into how temperature and ions govern RNA conformational transitions, offering a microscopic basis for RNA thermoadaptation, cold-induced misfolding, and RNA phase transitions.

**Statement of Significance:** Ions and temperature strongly influence RNA structure and dynamics, yet the molecular mechanisms by which these factors jointly regulate RNA behavior remain poorly understood. Using coarse-grained simulations with temperature-dependent Mg^2+^-phosphate interactions, we report how ion binding reorganizes around single-stranded RNAs as temperature increases. We found that unstructured RNAs undergo a non-monotonic structural transition: thermal disruption of base stacking first expands the chain, followed by the collapse driven by enhanced Mg^2+^ binding. This collapse arises from a temperature-induced migration of Mg^2+^ from diffusive ion atmosphere to direct inner-sphere binding, linking RNA compaction to ion dehydration and entropy-driven binding. These results reveal that the RNA ion atmosphere is a dynamic, structure-coupled component of RNA organization and provide a mechanistic basis for thermoresponsive RNA condensation.

**Graphical Abstract:** 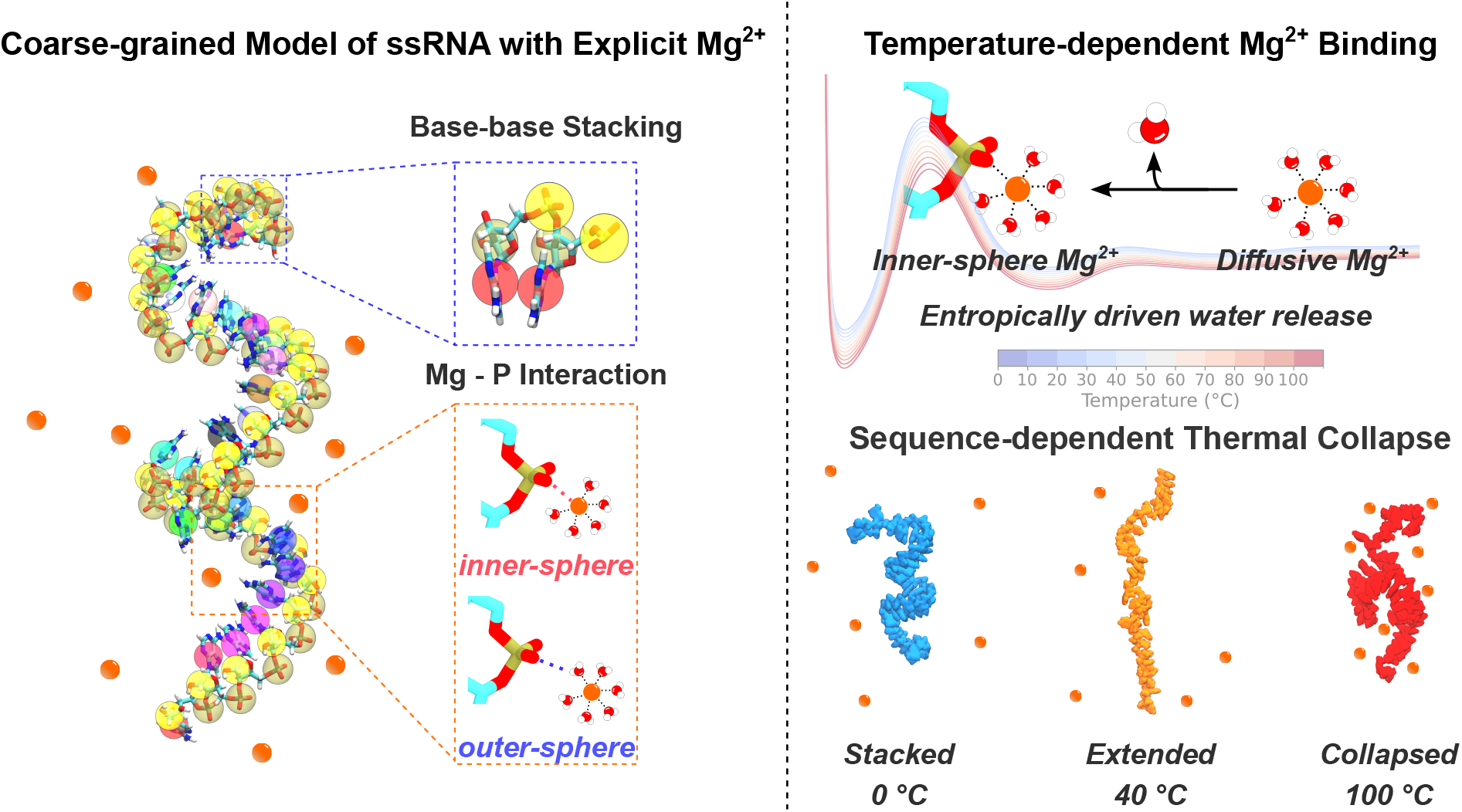

## Introduction

RNA plays a central role in virtually all biological processes, with its function intricately linked to its structure.^1–7^ The structural landscape of RNA is highly dynamic and shaped by a wide range of intrinsic and extrinsic factors,^8^ including the primary sequence,^9^ physiological conditions,^7^ ion concentrations, ^10–14^ molecular interactions with other biomolecules, ^15,16^ as well as covalent RNA modifications.^17–20^ Among its diverse structural motifs, single-stranded RNA (ssRNA) is particularly versatile. Beyond serving as linkers between structured regions, ssRNA contributes to higher-order RNA architecture and functions as regulatory elements throughout the mRNA lifecycle. ^21–26^ A wide array of cellular proteins, including those that are highly expressed, exhibit selective affinity for ssRNA, mediating essential steps in gene expression at both the transcriptional and post-transcriptional levels.^27–31^

The interactions between ssRNAs and regulatory proteins are expected to be dependent on sequence-specific structures. However, the molecular mechanisms underlying many processes involving ssRNAs remain poorly understood, primarily due to the limited understanding of their structural features. This challenge has persisted since the pioneering studies of the late 1950s.^32^ Early experimental techniques provided the fundamental insights for both intrinsic and extrinsic factors influencing the single-stranded helices of poly(rA), poly(rU), and poly(rC).^33–44^ In contrast, poly(rG) readily self-associates into higher-order structures, most notably the G-quadruplex.^45^ These studies identified base stacking as a dominant energetic determinant of helix formation, with temperature and ion concentration significantly impacting the helix formation kinetics. ^46^ Subsequent experimental advances have highlighted the substantial effects of Mg^2+^ on the structures of ssRNA.^47–51^ More recently, with growing interest in understanding biomolecular condensates, ssRNA has been shown to interact with arginine/glycine-rich intrinsically disordered regions, affecting the viscoelastic and dynamic properties of these condensates.^52–56^ Short poly(rA) sequences form Mg^2+^-dependent condensates, yet the structural organization of ssRNA within these assemblies remains largely unknown.^57^

Atomistic molecular dynamics (MD) simulations provide a detailed framework to probe ssRNA conformational ensembles. Recent improvements in ion-phosphate parameterization have led to more reliable descriptions of ion effects on RNA structure and dynamics.^58,59^ Coarse-grained (CG) models offer a complementary approach by reducing the number of degrees of freedom, enabling the study of larger systems and longer timescales. However, this simplification can limit the ability to capture fine-scale structural details. Existing CG models have primarily focused on well-folded RNAs. Applications to ssRNAs reveal that some approaches overestimate ssRNA size, potentially due to simplified representations of base stacking and *ad hoc* ion-RNA interactions. ^60–62^ Despite these limitations, such models provide valuable insights into the microscopic behavior of RNA.^63–66^ In our previous work, we extended the Single-Interaction-Site (SIS)^67^ model by incorporating explicit Mg^2+^-RNA interactions, revealing features of the ssRNA ion atmosphere.^68^ However, the SIS model does not explicitly account for base stacking, limiting its ability to describe sequence-dependent structural ensembles of these homopolymeric RNAs.

To overcome these limitations, we resorted to the Three-Interaction-Sites (TIS) model of RNA, where each nucleotide is represented by three CG beads corresponding to the ribose, phosphate, and nucleobase groups (Fig. 1A).^64^ In this work, we further develop temperature-dependent potentials of mean force (PMFs) for divalent ion-phosphate interactions, extending our previously reported model^69^ to incorporate thermal effects via liquid-state integral equation theory.^70–73^ The resulting PMFs naturally capture both inner- and outer-sphere coordination of Mg^2+^ with RNA, while interactions involving monovalent ions are treated implicitly through a mean-field Debye–Hückel (DH) term. Compared to our earlier model,^69^ the present implementation introduces revised functional forms and an optimized parameter set (see Methods section). These refinements improve numerical stability and yield a more accurate balance among base stacking, backbone flexibility, and electrostatic interactions.

**Figure 1:**
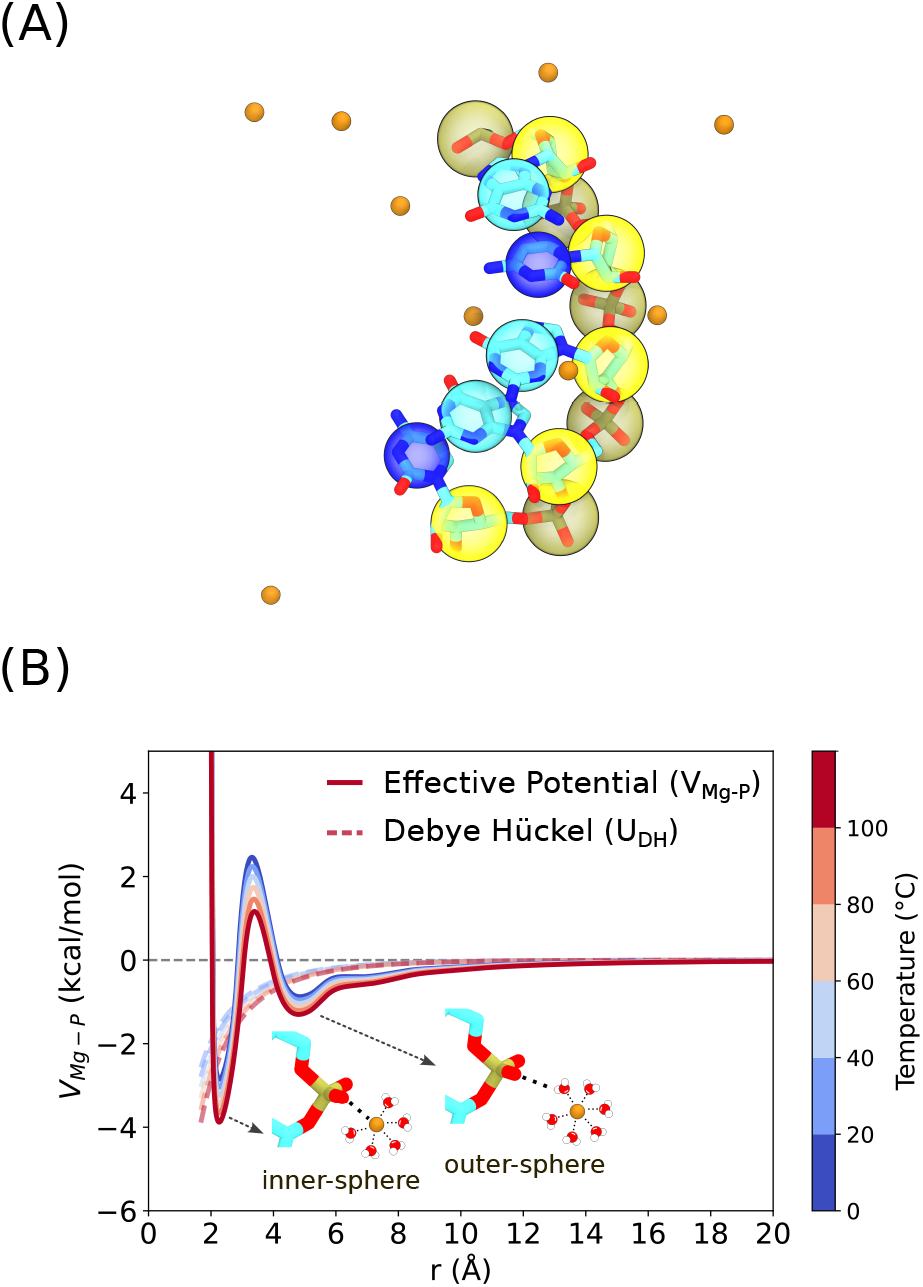
**(A)** Three-interaction-site (TIS) representation of RNA. Beads represent the ribose, phosphate, and base, with explicit Mg^2+^ ions. **(B)** Temperature-dependent effective potential between Mg^2+^ and phosphate. At short distances, the potential captures both inner-sphere and outer-sphere coordination. At longer distances, it transitions smoothly to the Debye-Hückel form, reflecting electrostatic screening, and plateaus at large separations.

Using unbiased CG simulations, we find excellent agreement between the calculated ssRNA conformational ensembles and experimental small-angle X-ray scattering (SAXS) data,^50^ capturing transient helical structures populated by rA_30_. More importantly, we reveal a pronounced temperature dependence of ssRNA conformations and ion distributions. As temperature increases, RNA undergoes a non-monotonic transition, first expanding and subsequently collapsing, due to the interplay between base unstacking and temperature-enhanced ion-mediated attractions. Strikingly, we observed a reorganization of the ion atmosphere at elevated temperatures in which diffuse ions condense inward to form direct contacts with the phosphate backbone, leading to the partial collapse of the extended ion cloud. Together, these findings establish a robust computational framework for investigating the structural and electrostatic features of RNA under varying ionic and thermal conditions.

## Methods

### Three-interaction-site (TIS) RNA model with explicit Mg^2+^

The TIS model represents each nucleotide with three spherical beads (interaction sites), corresponding to the phosphate, ribose and nucleobase. ^74^ The beads are positioned at the center-of-mass (COM) of the chemical groups (Fig. 1A). The details of the model are presented elsewhere.^69,74^ Briefly, the energy function describing the interactions between RNA beads is expressed as a sum of contributions from the bond length (*U*_*BL*_), bond angle (*U*_*BA*_), single-stranded stacking (*U*_*ST*_), hydrogen bonding (*U*_*HB*_), excluded volume (*U*_*EV*_), and electrostatic (*U*_*EL*_) interactions:

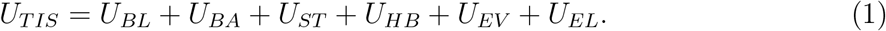

We use the harmonic approximation for *U*_*BL*_ and *U*_*BA*_, where the equilibrium values (*k*_*b*_, *k*_*θ*_, *r*_0_, and *θ*_0_) are listed in Tables S1 and S2. Excluded volume interactions are modeled using the Weeks–Chandler–Andersen (WCA) potential:^75^

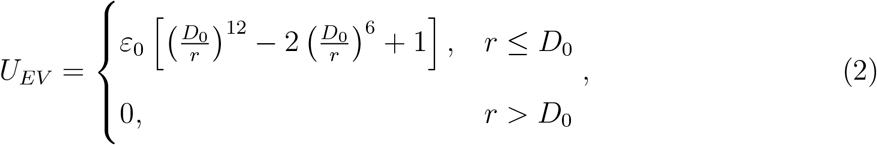

where *ε*_0_ = 1 kcal/mol and *D*_0_ = *R*_*i*_ +*R*_*j*_. *R*_*i*_ for ions and RNA beads are listed in Table S3. The WCA term vanishes if the interacting sites are separated by a distance greater than *D*_0_. For interactions between two bases, we set *D*_0_ = 3.2 Åto favor stacking.

#### Stacking potential

Stacking interactions *U*_*ST*_ are applied to any two consecutive nucleotides along the chain:

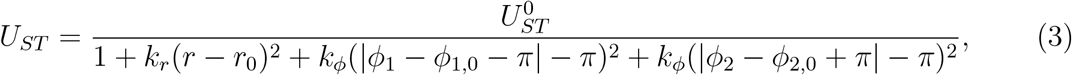

where *r* is the distance between B_*i*_ and B_*i*+1_, *ϕ*_1_(*P*_*i*_, *S*_*i*_, *P*_*i*+1_, *S*_*i*+1_) and *ϕ*_2_(*P*_*i*+2_, *S*_*i*+1_, *P*_*i*+1_, *S*_*i*_) are dihedral angles around the sugar-phosphate bonds (Fig. S1A). The absolute values ensure the dihedral angles are continuously wrapped around −*π* and *π* such that positive and negative dihedrals are treated equivalently. To avoid introducing two minima, we shift the functional form so that a single minimum occurs between −*π* and *π*, while preserving the shape of the potential elsewhere (Fig. S1B). With this functional form, the simulations remain stable, with no observable energy drift (Fig S40, S41). The structural reference parameters *r*_0_, *ϕ*_1,0_, and *ϕ*_2,0_ were obtained by coarse-graining an A-form RNA helix,^76^ where the dihedral angles were averaged to yield *ϕ*_1,0_ = −2.587 rad and *ϕ*_2,0_ = 3.071 rad. Previous simulations suggested *k*_*r*_ = 1.4 and *k*_*ϕ*_ = 4 rad^−2^ to reproduce the fluctuations of (*r* −*r*_0_)^2^, (*ϕ*_1_ −*ϕ*_1,0_)^2^ and (*ϕ*_2_ −*ϕ*_2,0_)^2^ observed in NMR structure.^74^ Here, the A-form structure is used solely to define local geometric reference values within the coarse-grained representation. This choice does not constrain the ssRNA to adopt an A-form conformation, and the simulated ssRNA remain flexible.

To recalibrate the model, we simulate base stacking of all 16 RNA dinucleotides. We use the stacking potential *U*_*ST*_ described in Eq. 3 with 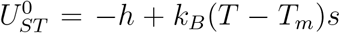 where *T* is the temperature (in Kelvin), and *h* and *s* are adjustable parameters representing enthalpic and entropic contributions, respectively. We classify configurations as stacked or unstacked based on *U*_*ST*_ following the original procedure.^74^ Specifically, configurations with *U*_*ST*_ *<* −*k*_*B*_*T* are classified as stacked, while those with higher energies are considered unstacked. This threshold corresponds to the thermal energy scale, and therefore provides a physically grounded criterion to identifying configurations that are energetically stabilized relative to thermal fluctuations. The stacking free energy for each dimer is then calculated using 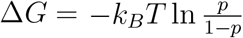 where *p* is the fraction of stacked configurations satisfying *U*_*ST*_ *<* −*k*_*B*_*T* (Figs. S21 - S36). The optimized *h* and *s* parameters used in simulations are listed in Table S4. The simulated thermodynamics parameters have excellent agreement with experimentally derived values (Fig. S1C).^74,77^

### Temperature-Dependent Ion-Phosphate Interaction

Following our previous procedure,^69^ we resort to the well-known Reference Interaction Site Model (RISM) theory to calculate the PMFs between Mg^2+^ and phosphate. We use 1D-RISM^70,71,73^ to calculate the radial distribution function *g*_*Mg*−*P*_ (*r*), and then convert it to PMF: *W* (*r*) = −*k*_*B*_*T* ln *g*_*Mg*−*P*_ (*r*). The theory starts with the Ornstein–Zernike (OZ) equation:

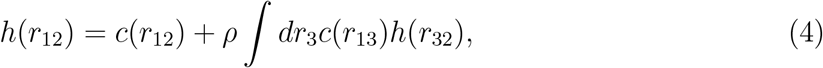

where *r*_*ij*_ is the distance between particles *i* and *j, c* is the direct correlation function, and *h* (the total correlation function) is related to *g* as *h*(*r*_*ij*_) = *g*(*r*_*ij*_) −1. To solve the OZ equation, another equation (known as the closure relation) connects *h* and *c*:

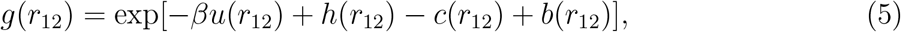

where *u*_*ij*_ is the pairwise potential energy function and *b*_*ij*_ is an unknown “bridge function”. In our work, we choose to use the partial series expansion of order-n (PSE-n) closure^78^ to improve the results for highly charged systems (compared to Kovalenko–Hirata closure^79^), while circumventing the convergence issues associated with the hypernetted-chain closure:^73,80,81^

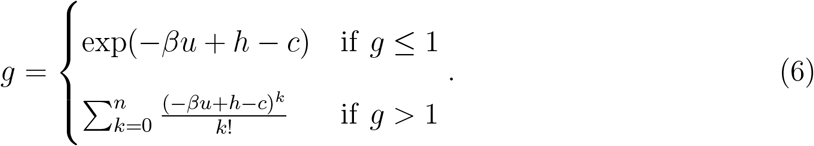

To ensure the effective potential has the correct long-ranged behavior in different conditions (ion, temperature, etc), we smoothly merge the short-ranged PMF *W* (*r*) with the mean-field DH potential *U*_*DH*_(*r*) as described in previous work:^69^

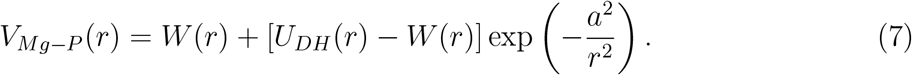

The parameter *a* is set to 5.0 Åto preserve the direct-contact binding energy of Mg^2+^. The resulting effective potential *V*_*Mg*−*P*_ (*r*) asymptotically reaches the DH approximation at large distances, while displaying water-mediated short-ranged interactions according to RISM theory (Fig. 1B).

Electrostatic interactions between other charged sites (phosphate-phosphate, and Mg^2+^-Mg^2+^ repulsions) are treated using DH theory, which approximates the screening behavior of implicit monovalent salt buffer:

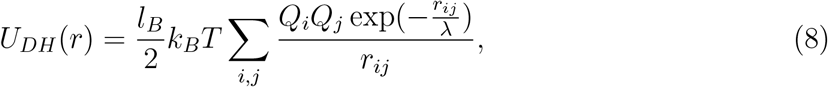

where *λ* = (8*πl*_*B*_*ρ*_1_)^−1*/*2^ is the DH screening length determined by the number densities of monovalent ions, *ρ*. The Bjerrum length 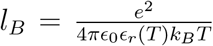 sets the strength of the interactions, where *e* is the elementary charge, *ϵ*_0_ is the vacuum permittivity, and *ϵ*_*r*_(*T*) = 87.74 −0.4008*T* +9.398 ×10^−4^*T*^2^ −1.410 ×10^−6^*T* ^3^ is the temperature-dependent dielectric constant of water (where T is in °C).^82^ The effective charge of phosphate, *Q*_*P*_ (*T, C*_1_, *C*_2_), depends on concentrations of both the monovalent (*C*_1_) and divalent ions (*C*_2_) using counterion condensation theory^83–85^ as described in our previous work.^69^ This approach adjusts the bare charge of phosphate according to ion concentrations and temperature, providing effective charges that reflect ion-mediated screening and condensation in a physically consistent manner.

By incorporating explicit Mg^2+^ while keep monovalent ions implicit, our framework provides a balance between physical realism and computational efficiency. It has been successfully applied to capture RNA folding and ion-mediated interactions of related systems.^68,69,86,87^ Monovalent ions (e.g. Na^+^, K^+^, Cl^−^) are not treated explicitly, and their effects are only included at the mean-field level. As a result, properties that depend on explicit counterion configurations, such as ion-ion correlations or spatially resolved monovalent ion distributions are not directly accessible. In addition, the present model focuses on Mg^2+^-phosphate interactions and does not explicitly include Mg^2+^-base interactions, which may become important for other RNAs.

### Langevin dynamics simulations

All simulations are performed using OpenMM 8.0.0^88^ on NVIDIA A100 Graphics Processing Unit (GPU). We use LangevinMiddleIntegrator in OpenMM to integrate the equations of motion in NVT ensemble with a timestep of 2 fs to ensure numerical stability due to the small size of Mg^2+^ ions. Mg^2+^ ions were initially placed at random positions within a cubic box containing the ssRNA of interest. The box size ranges from 420 Åto 720 Å, depending on the bulk Mg^2+^ concentrations, and is chosen such that at least 200 Mg^2+^ ions are present. We apply periodic boundary conditions in all three dimensions to minimize finite-size effects. We use low-friction Langevin dynamics to enhance conformational sampling, with a friction coefficient of 0.01 ps^−1^. Snapshots are saved every 10,000 integration steps, and all trajectories are examined for equilibration using the timeseries analysis implemented in PyMBAR.^89^ The equilibration time *t*_0_ was determined by maximizing the number of effectively uncorrelated samples, thereby identifying the optimal production window. Data prior to *t*_0_ were discarded from all analyses. Statistical convergence was further evaluated by verifying the stability of key observables (e.g. *R*_*g*_, ion counts) over time, and by dividing each trajectory into two equal halves and confirming consistent ensemble averages between them. The simulations were run for 2 *µ*s in the absence of Mg^2+^ and 8–10 *µ*s for simulations with Mg^2+^.

### Analyses

#### Calculations of Mg^2+^ preferential interaction coefficient

First, all frames are aligned to the ssRNA COM. The local concentration of Mg^2+^ is subsequently computed as:

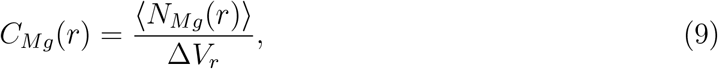

where ⟨*N*_*Mg*_(*r*)⟩is the average number of Mg^2+^ ions within the spherical shell of volume Δ*V*_*r*_ between *r* and (*r* +Δ*r*). The resulting number density is converted to molar concentration using the appropriate unit conversion, 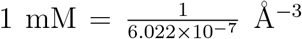. Because ssRNA is highly negatively charged, Mg^2+^ are preferentially accumulated near phosphate groups, leading to a depletion of Mg^2+^ in the bulk region. As a result, the effective bulk concentration, 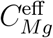 is lower than the initial value. We determine 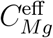 as the plateau value in the concentration profile *C*_*Mg*_(*r*) at large separation (Figs. S37, S38, and S39).

The preferential interaction coefficient (Γ_Mg_) reports the excess number of ions associated with RNA relative to bulk solution.^90,91^ We calculate Γ_Mg_ using the Kirkwood–Buff integral:

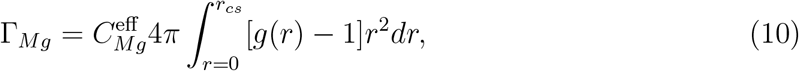

where 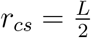 is half the simulation box length, and *r* is the distance from Mg^2+^ to the RNA COM.

#### Partitioning of inner-sphere, outer-sphere, and diffusive ions

A Mg^2+^ ion is classified as inner-sphere if the Mg-phosphate distance falls within the first peak of *g*_*Mg*−*P*_ (*r*) (*r <* 3.2 Å). Mg^2+^ within the second peak (3.2 Å*<g*_*Mg*−*P*_ (*r*) *<* 6.1 Å) is assigned to the outer-sphere population. The number of diffuse ions is then computed as *N*_diff_ = Γ_Mg_ −*N*_inner_ −*N*_outer_, where *N*_inner_ and *N*_outer_ correspond to bound ions, and *N*_diff_ represent the remaining excess ions that are not directly bound but are still part of the RNA ion atmosphere.

#### Calculations of SAXS Profiles from CG Trajectories

We first backmap 10,000 randomly selected conformations to all-atom representations using Arena.^92^ Because Arena only recognizes standard RNA atom names, we map the phosphate, sugar, and base beads in the CG model to P, C2’, and C4 atoms, respectively. To remove steric clashes after this procedure, each structure is subjected to an energy minimization using Amber bsc0 force field with *χOL*3 corrections (Fig. S2A).^93,94^ We then calculate SAXS profiles under the same conditions in experiments using CRYSOL with default parameters.^95^

#### Ion atmosphere 2D projection

Direct superposition of instantaneous ion distributions from successive trajectory snapshots provides a qualitative view of the ion atmosphere surrounding structured RNAs. However, for flexible ssRNA, such averaging leads to severe blurring that obscures spatial features of ion localization.^96^ To generate an interpretable representation of the Mg^2+^ ion atmosphere, we first remove the global translational and rotational motions of ssRNA by aligning each frame to the ssRNA COM and principle axes.

This alignment defines a consistent global reference frame, with the first principal axis corresponding to the 5’to 3’direction of the RNA. For each phosphate group, we then calculate the position of its closest Mg^2+^ ion relative to a local coordinate system centered at the phosphate. In this frame, the displacement vector between Mg^2+^ and phosphate is decomposed into an axial component (*r*_*z*_), defined along the RNA principal axis, and a lateral component 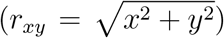, defined in the xy-plane. To preserve directional information in the 2D projection, the sign of the lateral displacement is assigned according to the sign of *θ*, where 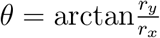, resulting in the directional in-plane distance: 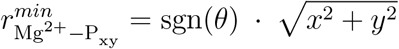. Together, (*r*_*xy*_, *r*_*z*_) provides a two-dimensional representation of the ion distribution that preserves both axial and directional information relative to the RNA. Because this representation is defined relative to each phosphate group after global alignment, it is insensitive to the overall curvature or conformational variability of the ssRNA chain (Figs. 3C, S19).

#### Calculations of solvent-accessible surface area (SASA)

SASA is calculated using the FreeSASA Python package.^97^ In this method, a spherical probe representing the solvent molecule rolls over the solute molecule. We use the classical Lee–Richards algorithm, ^98^ in which the surface area is approximated by the contour of successive slices. We set the probe radius to 2.0 Åto approximate the size of a hydrated Mg^2+^ ion. The total SASA can be decomposed into contributions from the phosphate, sugar, and nucleobase groups. Lacking atomic details, SASA estimated from CG models may be biased. To access this, we bench-marked CG-derived SASA against atomistic models and found a systematic underestimation of ≈10%, likely due to surface smoothing in the CG representation (Fig. S10). Since we are interested in the relative values, comparative analysis still remains meaningful.

#### Calculations of the orientational correlation function

The orientational correlation function (OCF) is defined as (Eq. 11):

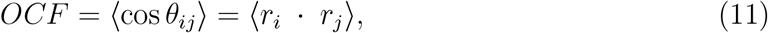

where *r*_*i*_ is the normalized bond vector connecting the *i*^*th*^ and (*i*+1)^*th*^ phosphate groups. The OCF is evaluated as a function of sequence separation |*i* −*j*|, which measures the correlation between local backbone orientations over increasing contour distances. The resulting OCF is averaged over all conformations in the structural ensemble to characterize the mean chain stiffness and directional persistence of ssRNA.

#### Statistical analysis and uncertainty estimation

Because MD trajectories are temporally correlated, individual frames were not treated as independent samples. Instead, each time series (e.g. ion counts) was divided into contiguous, non-overlapping blocks of 5,000 frames. To justify this block size, we analyze the normalized autocorrelation functions of representative observables (Fig. S44). The autocorrelation decays rapidly and approaches zero within ≈400 ps, indicating that configurations separated by this timescale are effectively uncorrelated. This correlation time is more than two orders of magnitude shorter than the block length (100 ns), ensuring that block-averaged samples are statistically independent. The mean value within each block was calculated, and these block-averaged values were treated as approximately independent samples, and the effective sample size was therefore given by the number of blocks. To estimate uncertainty of the mean values and the differences between sequences, nonparametric bootstrap resampling was applied to the block-averaged data. For each comparison, 40,000 bootstrap resamples were drawn, and the resulting distribution of mean differences was used to calculate 95%confidence intervals. These intervals are reported as the primary measure of statistical significance.

## Results and Discussion

### Distinct Structural Ensembles of rA_30_ and rU_30_

To access the model’s accuracy in capturing ssRNA structural ensembles, we compare the simulated radii of gyration (*R*_*g*_) and SAXS profiles^50^ for rA_30_ and rU_30_ at 20 °C (results for rC_30_ are in SI, Fig. S17). Across a wide range of concentrations of both Mg^2+^ and monovalent ions, *R*_*g*_ values agree well with SAXS-derived measurements, capturing ion-induced compaction (Figs. 2A, S7 and Table S5).^50,99^ Overall, our model predicts that rA_30_ consistently adopts more compact conformations than rU_30_ due to stronger adenine stacking interactions, which is in agreement with experimental data.^50^ Despite small *R*_*g*_ discrepancies, the computed SAXS profiles are in quantitative agreement with experiments (Fig. S4). Kratky plots further highlight sequence-dependent differences in structural ensemble and collapse upon addition of Mg^2+^ (Fig. 2B). Both RNAs remain largely unstructured across the tested ion conditions;however, at 5 mM Mg^2+^, the Kratky curve of rA_30_ bends downward, suggesting the formation of partially folded, compact conformations (Figs. S5 and S6). Such a difference between two constructs is further supported by the simulated end-to-end distances (*R*_*ee*_), where rA_30_ exhibits shorter *R*_*ee*_ than rU_30_ (Fig. S8, Table S7). The small discrepancies between simulation and experiment are primarily observed in the high-*q* region of the Kratky plots (Figs. S5 and S6). These deviations likely reflect the limited resolution of the coarse-grained model in capturing fine-scale structural features, as well as increased experimental uncertainty at high *q*, where scattering intensities are inherently weaker and more sensitive to noise. Nonetheless, the excellent agreement in low-*q* and intermediate-*q* regimes indicates that the model reliably captures global dimensions and overall conformational behavior of the RNAs.

**Figure 2:**
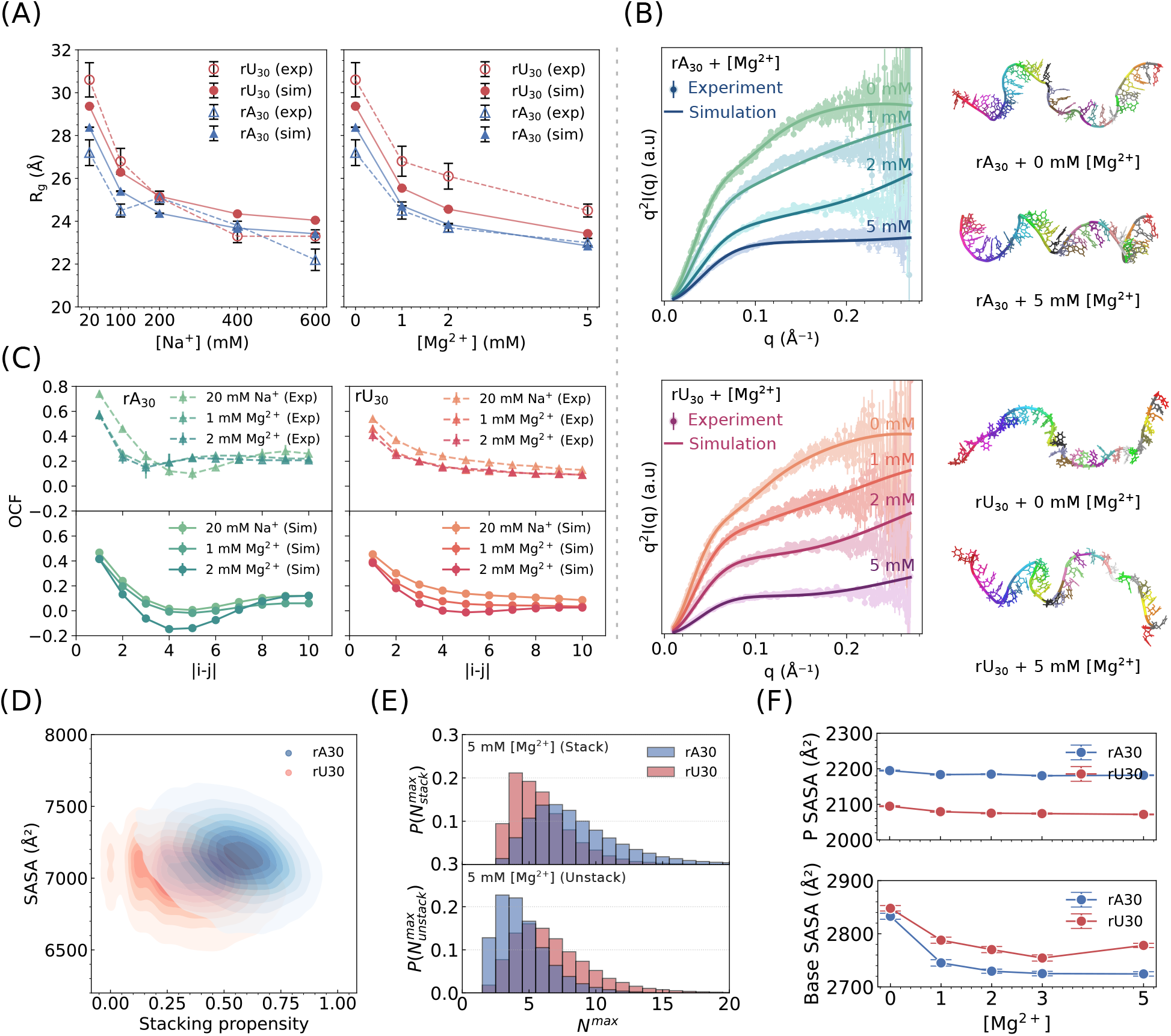
rA_30_ and rU_30_ populate distinct structural ensembles. **(A)** Radii of gyration from simulation plotted against SAXS-derived values for rA_30_ (blue) and rU_30_ (red) under varying Mg^2+^ and Na^+^ concentrations, showing sequence- and ion-dependent compaction. Error bars are standard error of the mean. **(B)** Comparison between simulated SAXS profiles with experiments for rA_30_ (top) and rU_30_ (bottom). On the right shown representative conformations obtained by all-atom backmapping from CG coordinates. Error bars are standard error of the mean. **(C)** Orientation correlation functions for rA_30_ (left) and rU_30_ (right) at various ion concentrations. **(D)** Solvent-accessible surface area *vs*. stacking propensity distribution of rA_30_ and rU_30_ in 5 mM Mg^2+^. **(E)** Distribution of maximum consecutive stacking (top) and unstacking (bottom) in 20 mM Na^+^ +5 mM Mg^2+^. **(F)** Decomposing solvent-accessible surface area to phosphate (top) and base (bottom) contributions. Error bars represent 95%confidence intervals.

**Figure 3:**
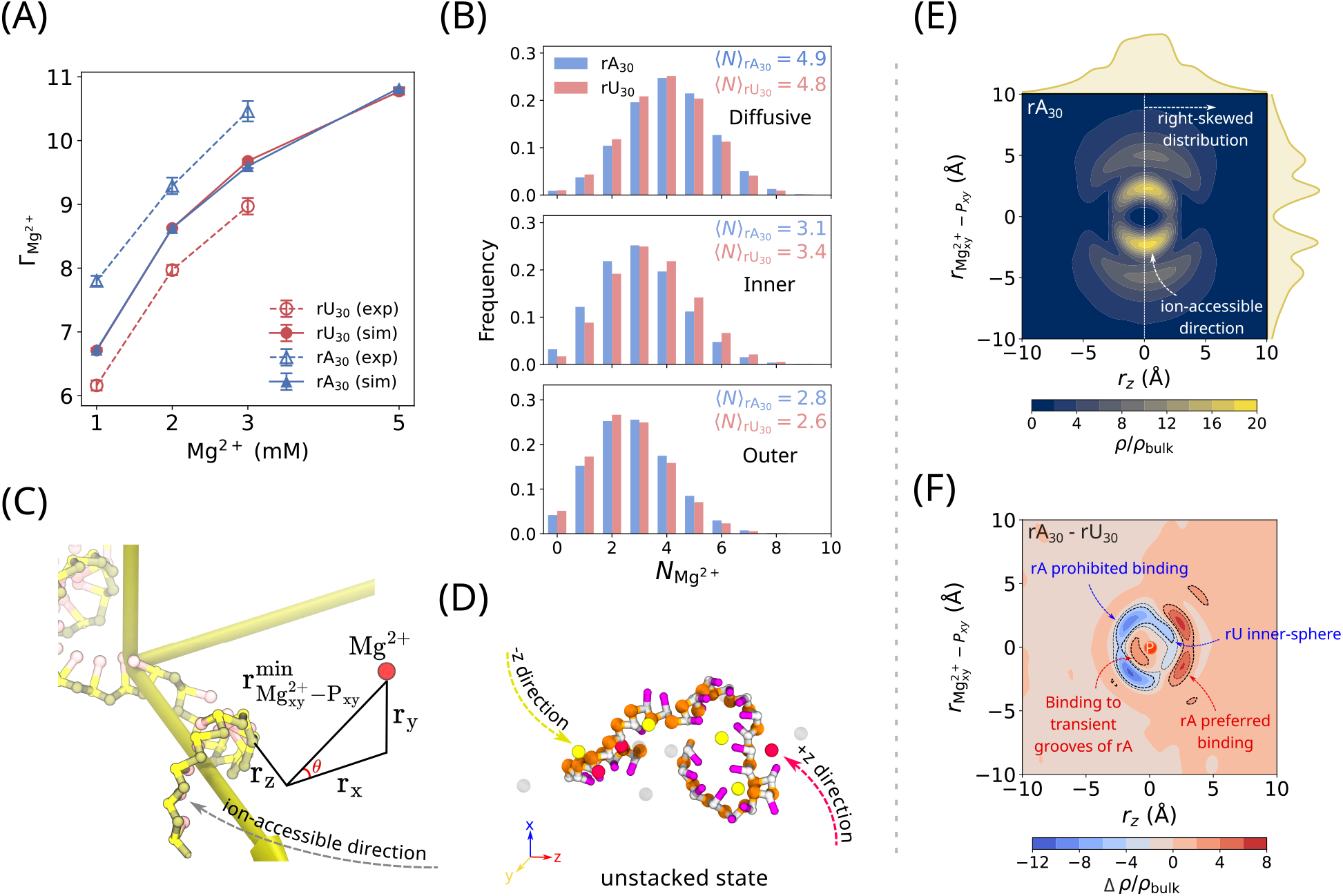
Sequence- and structure-dependent ion atmosphere. **(A)** Excess Mg^2+^ per chain for rA_30_ (blue) and rU_30_ (red) from simulations and buffer exchange-atomic emission spectroscopy.^50^ Error bars are standard error of the mean. **(B)** Partitioning of Mg^2+^ into diffusive (top), inner-sphere (middle), and outer-sphere (bottom) binding modes. **(C)** Schematic of the alignment for ion projection, where the RNA major axis is along the z-axis. Distances 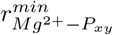 and *r*_*z*_ are Mg^2+^–phosphate separation in the xy- and z-directions, respectively;*θ* is the azimuthal angle. In the right-handed helical state, Mg^2+^ accessibility is restricted from a certain direction due to the presence of the sugar and base. **(D)** Schematic of the unstacked state, where Mg^2+^ can approach from both directions along the z-axis. **(E)** 2D density map of Mg^2+^ around rA_30_, showing a right-skewed distribution consistent with the transient helical structures. The ion-accessible direction corresponds to the most populated region. **(F)** Difference density map between rA_30_ and rU_30_. Red regions highlight preferred Mg^2+^ binding for rA_30_ in the positive z-direction and transient groove binding near the phosphate, whereas blue regions are sites more accessible in rU_30_.

To directly probe the ssRNA structures, we compute the OCF between phosphate groups (Fig. 2C). Changes in backbone direction reduce the OCF, which becomes negative when the vectors approach an antiparallel orientation. For rA_30_, OCFs exhibit oscillatory behavior across ionic conditions. With added Mg^2+^, the correlation decays rapidly, changing from positive to negative values, strongly suggesting a signature of helical structures. In contrast, rU_30_ shows monotonic decay in most conditions, suggesting a largely featureless chain. The overall shapes of our simulated OCFs are comparable to recently reported results from other CG RNA models.^61^ They are also qualitatively consistent with previous reconstruction of structural ensembles from SAXS for those sequences (Fig. S9).^50^ We note that multiple distinct conformational ensembles can yield similar SAXS profiles, underscoring the need for local structural probing to fully resolve such degeneracy.

### Stacking Interactions Dictate Helical Formation

To elucidate the difference in helical propensity between the two sequences, we calculate the 2D distributions of SASA and stacking propensity, defined as the fraction of nucleobases in stacked conformations (results for 20 mM Na^+^ +5 mM Mg^2+^ shown in Fig. 2D). The two distributions of rA_30_ and rU_30_ are well separated, suggesting distinct conformational ensembles. rA_30_ transiently adopts a right-handed single-stranded helical conformation (Fig. 2B), characterized by a maximum contiguous stacking length 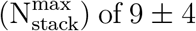 of 9 ±4 nucleotides, whereas rU_30_ exhibits shorter stacked segments (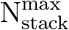 of 6 ±2) (Fig. 2E, top). As a result, the coil (unstacked) regions are shorter for rA_30_ (5 ±2 nucleotides) than for rU_30_ (7 ±3 nucleotides), consistent with stronger base stacking in the former (Fig. 2E, bottom). Because stacking interactions occur primarily between nearest neighbors, such helicity is expected to be only weakly cooperative.^100,101^ Our results agree with single-molecule force experiments that revealed alternating helical and coil regions for poly(rA), but not for poly(rU).^102^ It is also supported by crystallographic analyses of the ApApA trimer demonstrating intrinsic helical order in poly(rA).^103^ Together, our simulations support a block copolymer picture of ssRNA, in which stacked and unstacked domains coexist.

These differences in stacking behavior between rA_30_ and rU_30_ directly affect solvent exposure. In particular, the helical organization of rA_30_ orients phosphate groups outward, resulting in higher phosphate SASA for rA_30_ compared to rU_30_ (Fig. 2F, top). On the other hand, nucleobase SASA is lower in rA_30_ due to stronger stacking and thus reducing base exposure (Fig. 2F, bottom). Such modulation of solvent exposure may in turn influence local ion interactions and potentially affect the recognition of single-stranded regions by regulatory proteins. ^104,105^

### Structure-Dependent Ion Atmosphere

The degree of ion binding to RNA can be quantified using the preferential interaction coefficient (Γ_Mg_), which reports the excess number of ions associated with RNA relative to bulk solution.^90,91,106^ Consistent with ion counting experiments,^50^ our simulations reproduce the overall magnitude of Γ_Mg_ (Fig. 3A, Table S6). Experimentally, rA_30_ attracts approximately 1.5 more Mg^2+^ ions per chain than rU_30_ over the range of 0 - 3 mM Mg^2+^ and 20 mM Na^+^, corresponding to a lower competition coefficient (*M*_1*/*2_ = 0.9 ±0.1 mM for rA_30_ vs. 1.3 ±0.3 mM for rU_30_). Although these values overlap within experimental uncertainty, their mean difference implies that nearly 40%higher bulk Mg^2+^ concentration is required to displace half of the monovalent ions surrounding rU_30_.^50^ This subtle sequence dependence is not captured by our simulations, which likely reflects the level of coarse-graining in the model, including the implicit treatment of monovalent ions and the absence of explicit base-ion interactions. However, prior work on ssDNA also found identical Γ_Mg_ for dA_30_ and dT_30_ despite pronounced differences in stacking propensity.^49,87^

To further probe the ion atmosphere in greater detail, we partition the Mg^2+^ binding into inner-sphere (direct phosphate coordination), outer-sphere (water-mediated interaction), and diffusive regions (Fig. 3B).^68,107^ We note that while ion-counting experiments measures the overall extent of Mg^2+^ association, detailed features such as coordination modes, and spatial ion distributions are currently not directly resolved experimentally. The majority of Mg^2+^ remain in the diffusive regime, consistent with our previous findings that unstructured RNAs favor diffuse, weakly localized ionic environments.^68^ Interestingly, we find that rU_30_ has slightly more inner-sphere Mg^2+^ binding than rA_30_, with mean counts of 3.40 ±0.03 versus 3.09 ±0.03, respectively. This result suggests that the overall ion atmosphere of rU_30_ is slightly more compact than rA_30_ despite both sharing the same Γ_Mg_. Such a small difference could be amplified in longer sequences, multi-component assemblies or condensates.

To visualize sequence-dependent ion binding patterns, we align the RNA COM and principle axes, thus removing the RNA translational and rotational movement. Subsequently, we project the distance between every Mg^2+^ and its closest phosphate group onto a 2D surface (see Methods). Since rA_30_ has a small preference for helical structures, stacked phosphates align along the positive z-axis from 5’to 3’, establishing a preferred direction of ion approach toward the backbone (Fig. 3C). In contrast, in the unstacked state, phosphates are disordered, allowing ions to approach from both the +z and −z directions (Fig. 3D). The resulting 2D density map of rA_30_ shows a slightly right-skewed distribution along the z-axis, consistent with the anisotropic ion localization expected for a transient helical structure (Figs. 3E, S20). Additional insights emerge by subtracting the rU_30_’s distribution from rA_30_’s (Fig. 3F): a distinct region of enhanced Mg^2+^ density appears along the positive z-axis, indicating a site where rA_30_ preferentially binds ions. Conversely, the region in the corresponding negative z-axis is depleted for rA_30_, suggesting that this binding site is more accessible in rU_30_. Furthermore, localized hotspots of density indicate Mg^2+^ association with the transient helical grooves of rA_30_, reflecting its stacked, groove-forming structure that is absent in the largely unstacked rU_30_ ensemble. In other words, the chirality of the RNA imposes a chiral organization on its counterions. This phenomenon is analogous to how water molecules form a chiral hydration structure around duplex DNA, aligning in helical patterns along the major and minor grooves of the double helix.^108–111^ Together, these results demonstrate that Mg^2+^ ions neutralize ssRNA by organizing to form a complex ion atmosphere shaped by the RNA’s 3D organization. The ion atmosphere is therefore an extension of the RNA’s structure, inheriting its helical grooves, directional geometry, and chirality rather than forming a uniform shell.^112^ Such a coupling between the RNA shape and ion distribution underscores that ions “sense”the helix’s handedness and stabilize the structure in an asymmetric fashion, which has important implications for RNA folding and molecular recognition.

### Thermal Compaction of Homopolymeric ssRNA

The newly developed PMF allows us to study how ions modulate RNA conformations at various temperatures. Remarkably, we found that *R*_*g*_ values of ssRNAs display non-monotonic behaviors when the temperature increases from 0 °C to 100 °C (Fig. 4B). Even in the absence of Mg^2+^, rA_30_ exhibits a relatively long expansion phase, reaching a maximum at *T*_max_ = 76 °C before undergoing mild collapse, whereas rU_30_ shows limited expansion and collapses earlier at *T*_max_ = 48 °C. Introduction of 1 mM Mg^2+^ sharply lowers *T*_max_ (to 59 °C for rA_30_ and 15 °C for rU_30_), with little additional shift up to 5 mM.

**Figure 4:**
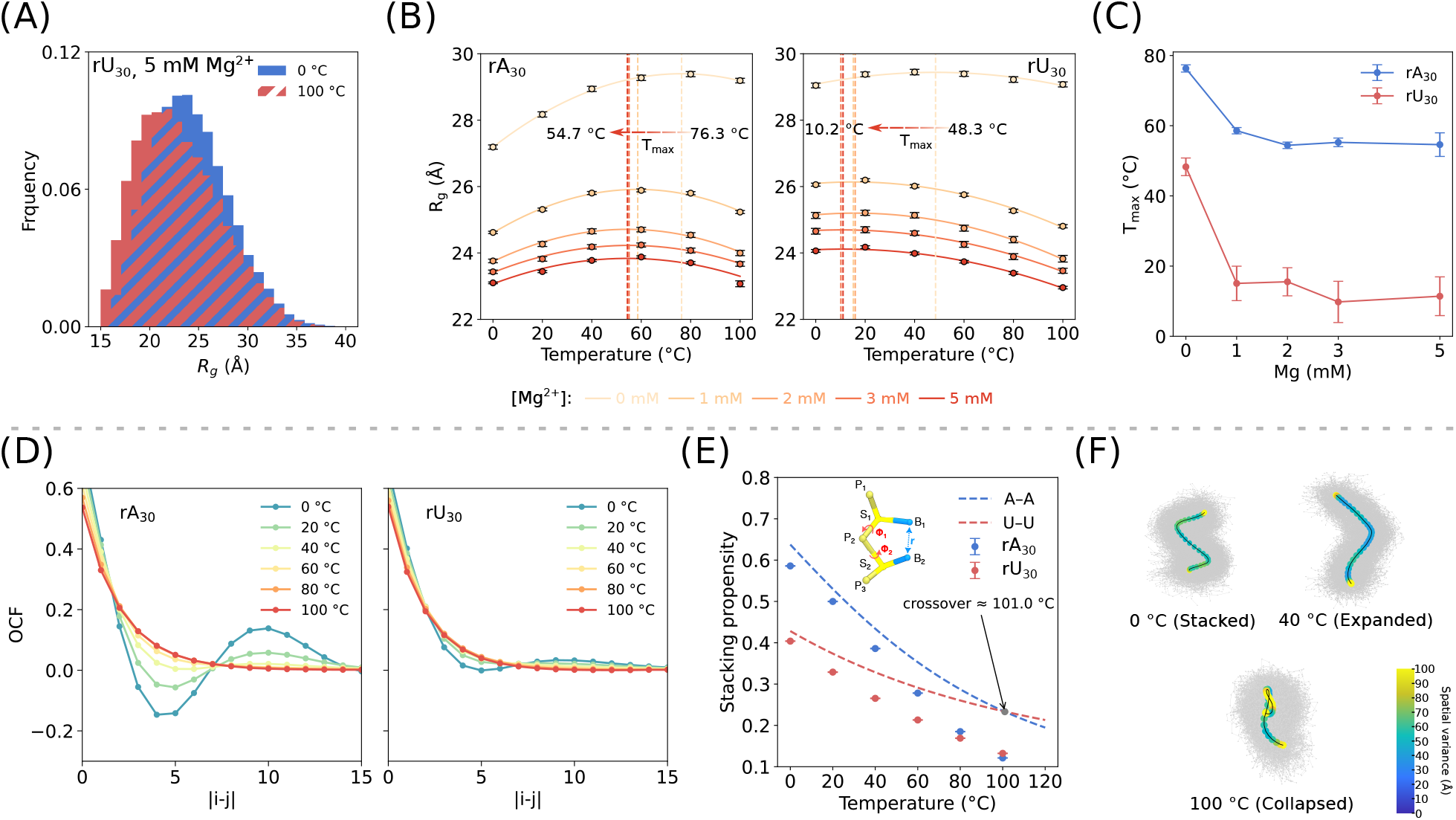
ssRNA collapses at high temperature. **(A)** Distribution of *R*_*g*_ at 0 °C (blue) and 100 °C (red) of rU_30_ in 5 mM Mg^2+^. **(B)** *R*_*g*_ of rA_30_ (left) and rU_30_ (right) as a function of temperature in 0-5 mM Mg^2+^. Vertical dashed lines denote *T*_max_, the temperature of maximal expansion, which marks the transition from chain expansion to collapse. Error bars represent 95%confidence intervals. Numerical values are reported in Table S8 and S9. **(C)** Dependence of *T*_*max*_ on Mg^2+^ concentration for rA_30_ (blue) and rU_30_ (red). Error bars are standard error of the mean. **(D)** Orientational correlation functions from 0 °C to 100 °C for rA_30_ (left) and rU_30_ (right). **(E)** Stacking propensity decreases with temperature for rA_30_ and rU_30_. Dotted lines show prediction from the dinucleotide model, which underestimates the accelerated decay observed in long sequences. Error bars are standard error of the mean. **(F)** Representative conformational ensemble of rA_30_ at 0 °C, 40 °C, and 100 °C.

To further dissect the conformational ensembles of ssRNAs across temperatures, we examined the 2D distribution of SASA and stacking propensity at various ionic conditions. As temperature increases, all RNAs exhibit a shift towards lower stacking propensity and higher SASA. This shift occurs more rapidly for rA_30_ than for rU_30_ (Figs. S12). Consistently, projection of the higher-dimensional space comprising of *R*_*g*_, *R*_*ee*_, SASA, stacking potential and Mg–P interaction energies onto two dimensions using t-SNE reveals the same trend (Figs. S11, S13). For rA_30_, pronounced helical signatures are evident at 0 °C and 20 °C, characterized by oscillatory OCFs that alternate between negative and positive values. These helical correlations vanish at high temperatures, indicating loss of ordered stacking. In contrast, rU_30_ displays only weak helicity even at 0 °C, and its OCFs decay monotonically at all temperatures, suggesting that uracil-rich structures respond more weakly to thermal perturbation (Fig. 4D). Compared with their corresponding dinucleotides, both rA_30_ and rU_30_ unstack more rapidly upon heating (Fig. 4E). Structural ensembles^113^ of rA_30_ highlight drastic changes across temperatures: stacked and compact conformations dominate at low temperatures, expanded intermediates emerge at moderate temperatures, whereas collapsed structures prevail at high temperatures (Fig. 4F).

Similar temperature-induced collapse has been observed in all-atom simulations of poly phosphate.^52^ Light scattering experiments on poly(rA) also found it collapsing when temperature increases, where stacking disruption was proposed as the main mechanism.^114^ However, disruption of base stacking is expected to render rA_30_ indistinguishable from rU_30_, leading to a larger *R*_*g*_. Consistent with this expectation, at high temperatures where the stacking fractions of both sequences converge to similar values (Fig. 4E), we observe that their *R*_*g*_ become identical, and the sequence dependence vanishes (Fig. 4B).

To understand why such collapses occur, we turn our attention to ion-RNA interactions as hinted by the PMF. It has been known that the binding of divalent ions to phosphate groups is driven by entropy, thus increasing temperature leads to a more stable binding. ^115,116^ In our framework, temperature affects the ion binding through two coupled terms: (i) the inverse thermal factor *β* = 1*/k*_*B*_*T* in the RISM theory and when calculating *W* (*r*) from *g*_*Mg*−*P*_ (*r*), and (ii) the dielectric constant of water, *ϵ*_*r*_(*T*), which decreases at elevated temperature and thereby effectively strengthens electrostatic interactions. Together, those effects stabilize both inner- and outer-sphere minima of the Mg^2+^–phosphate PMF at high temperatures (Fig. 1B). We note that this enhanced affinity has an entropic origin, arising from the release of ordered water molecules and competing counterions into the bulk solution. In particular, the temperature dependence of the dielectric constant of water gives rise to a substantial entropic contribution to the electrostatic interaction used in CG modeling, as illustrated in a recent work.^117^

We rationalize the non-monotonic behavior of *R*_*g*_ by considering the coupled effects of electrostatics and base stacking. The Bjerrum length 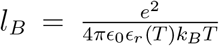 (where *ϵ*_*r*_ is the temperature-dependent dielectric constant of water) sets the strength of electrostatic interactions. With increasing temperature, *ϵ*_*r*_*T* decreases, resulting in an overall lengthening of *l*_*B*_ (Fig. S14A). Consequently, electrostatic interactions become stronger at higher temperatures, promoting enhanced counterion condensation on the phosphate backbone (Fig. S14B). The resulting effective P–P repulsion thus exhibits a non-monotonic temperature dependence: it initially increases as the electrostatic interactions strengthen but subsequently weakens once extensive counterion adsorption more effectively neutralizes the backbone charges (Fig. S14D). In the presence of Mg^2+^, such effect is much more pronounced because of its stronger affinity towards phosphate groups at elevated temperatures (Fig. 1B), which has been demonstrated in all-atom simulations.^52,61,118^

Additionally, base-base stacking further modulates this behavior in a sequence-dependent manner. Stacking propensity decreases monotonically with temperature (Fig. 4E), so the thermal disruption of stacking interactions drives an initial expansion of RNA. The overall temperature profile of *R*_*g*_ therefore reflects a balance between stacking and electrostatic effects: *R*_*g*_ first increases as stacking weakens, then subsequently decreases as counterion condensation dominates at higher temperatures. Adenines stack more favorably than uracils, explaining why rA_30_ expands substantially upon heating, whereas rU_30_ with weak stacking shows minimal expansion. Interestingly, Mg^2+^ affects stacking interactions minimally (Fig. S15). Therefore, the pronounced dependence of *T*_max_ on Mg^2+^ arises mainly because divalent cations are far more effective at neutralizing RNA charge than monovalent ions, altering the electrostatic balance that governs the transition. This mechanism also explains the observed saturation at high Mg^2+^ concentrations.

To directly assess the role of the temperature-dependent ion interactions, we also perform control simulations in which the Mg^2+^-phosphate PMF is held fixed with respect to temperature. Under these conditions, the ssRNA does not undergo collapse upon heating in the presence of Mg^2+^ (Fig. S42A). These results demonstrate that temperature-dependent ion interactions are essential for driving the observed collapse behavior.

We note that structured or partially folded RNAs generally display monotonic expansion with increasing temperature.^119^ One key difference between the sequences studied here and structured RNAs is the lack of a stable tertiary fold in unstructured RNAs. As a result, those RNAs do not undergo unfolding at high temperature, which often correlates to a large expansion in size, as seen in folded domains. The collapse at high temperatures observed here and elsewhere^52,114^ does not correspond to a formation of folded or native-like structure, but it reflects polymer-level reorganization driven by enhanced ion condensation. In the case of folded structures, this ion-driven collapse is masked by a much larger expansion due to RNA unfolding. It is therefore desirable to separate the contribution of ion-facilitated interaction from RNA unfolding using the simple systems studied here.

### Reorganization of the Ion Atmosphere at Elevated Temperature

To quantify how ions induce such collapses, we calculate Γ_Mg_ at various temperatures. Across all conditions, Γ_Mg_ increases slightly with temperature (Fig. 5A, Tables S10, S11), indicating enhanced Mg^2+^ absorption into the RNA ion atmosphere. Our result is consistent with the strengthening of Mg^2+^-P and thus Mg^2+^-RNA interactions at high temperatures.^52,61,115,116,118^ Strikingly, our simulations unveil a clear redistribution of Mg^2+^ from the diffusive to the inner-sphere populations as temperature rises, while the outer-sphere contribution remains largely unchanged (Fig. 5B). Quantitatively, the number of inner-sphere ions increases by nearly 1.5 per RNA, accompanied by a corresponding decrease of about 1.0 ion in the diffusive population. At elevated temperatures, the inner-sphere ions outnumber the diffusive counterparts, signaling a temperature-driven collapse of the Mg^2+^ atmosphere surrounding the RNA (Fig. S18). 2D ion density maps comparing between 0 °C and 100 °C vividly illustrate this rearrangement (Fig. 5D). For rA_30_, the diffusive layer becomes markedly depleted as ions migrate inward to form direct, inner-sphere contacts with phosphate groups. The concurrent loss of base stacking further facilitates this redistribution. rU_30_ also exhibits a similar shift (Fig. 5C), but the magnitude is much less pronounced, consistent with its weaker stacking propensity (Fig. 5D). In control simulations where the Mg^2+^-phosphate interactions are held fixed with respect to temperature, we observe no shift from diffusive to inner-sphere coordination (Fig. S42B, C). In such cases, the diffusive contribution keeps increasing with temperature.

**Figure 5:**
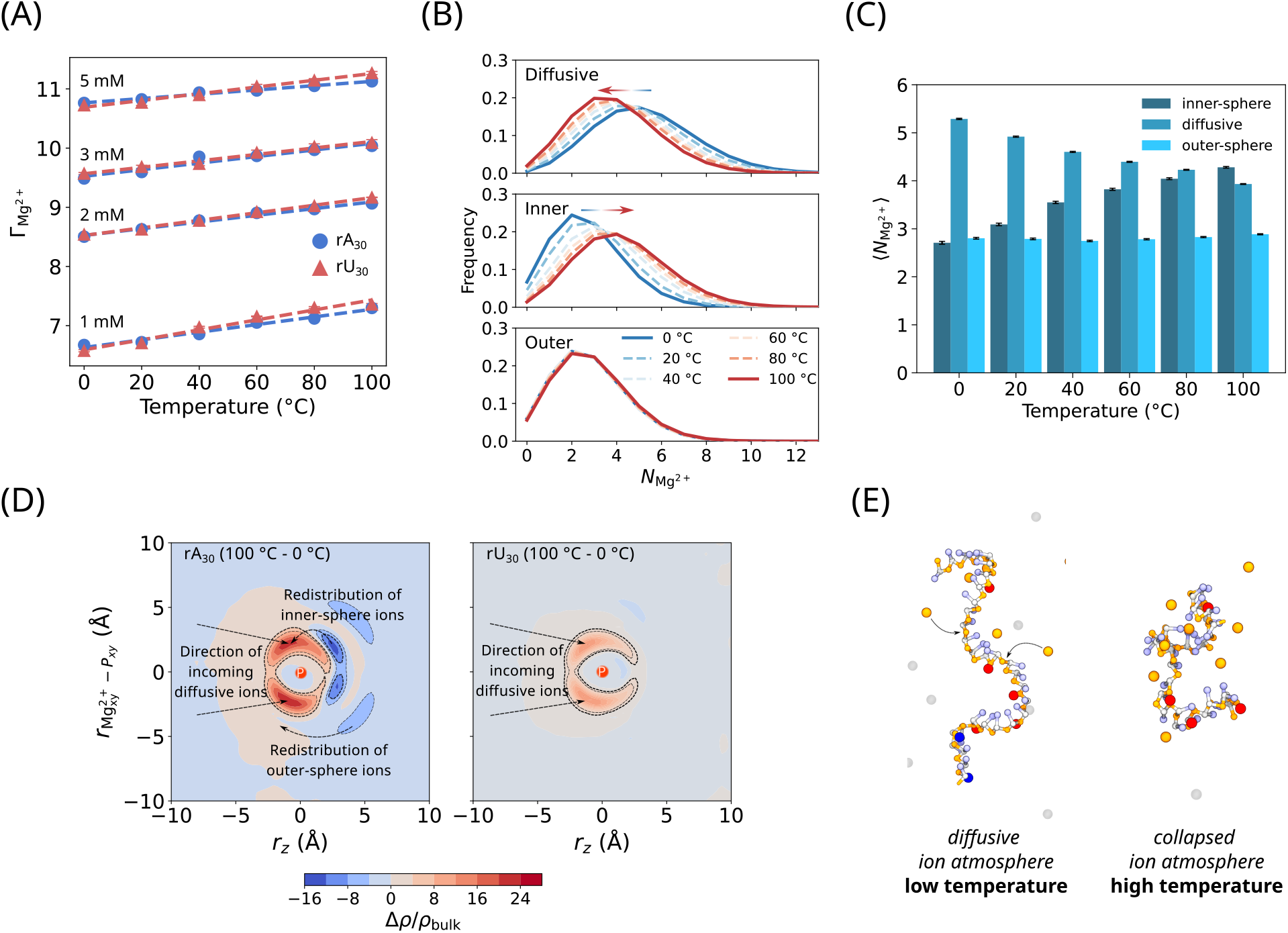
Ion reorganization at high temperature. **(A)** Preferential interaction coefficient for Mg^2+^ increases linearly with temperature for both rA_30_ (blue) and rU_30_ (red) at all Mg^2+^ concentrations. Error bars are standard error of the mean. **(B)** Partitioning of rA_30_ ion atmosphere into diffusive (top), inner-sphere (middle), and outer-sphere (bottom) bindings. Increasing temperature drives ions from the diffusive to the inner-sphere regions. **(C)** Ion atmosphere composition from 0 to 100 °C of rU_30_ in 5 mM Mg^2+^. Error bars represent 95%confidence intervals. **(D)** Difference in 2D ion density map (100 °C - 0 °C). **Left:** rA_30_ shows heating reorganizes diffusive ions towards inner-sphere binding, while outer-sphere and inner-sphere ions shift with the loss of stacking. **Right:** Reorganization is much weaker for rU_30_ than in rA_30_ due to smaller changes of stacking during heating. **(E)** Schematic illustration of diffusive-to-inner-sphere ion migration underlying temperature-induced collapse of the ion atmosphere.

This temperature-driven transition therefore reflects an entropically favored reorganization of the ion atmosphere. Formation of inner-sphere contacts requires partial dehydration of Mg(H_2_O)_6_^2+^, an energetically costly process that releases tightly bound water molecules into bulk solvent. The resulting gain in translational and rotational entropy of released water grows with temperature, progressively offsetting the dehydration enthalpy and thus stabilizing direct Mg^2+^–phosphate coordination. Such entropy-driven binding and assembly processes have also been found in the ion-induced assembly of polyphosphates,^52^ RNA,^52,61,118^ and polyelectrolyte coacervation.^117,120,121^ Our findings have implications for thermoresponsive behavior of RNA condensates. The expansion of ssRNA at relatively low temperature followed by collapse at high temperature (Fig. 5E), as observed here, is a hallmark of lower critical solution temperature (LCST)-type phase separation.^52,122–124^ The strong coupling between coil-globule transitions and Mg^2+^ binding uncovered in this study may provide a microscopic basis for LCST-driven RNA condensation, and help explain how RNAs maintain structural integrity and function across a wide thermal range, including in thermophilic environments,^125–128^ or cold-induced RNA misfolding transitions.^129^

## Conclusion

Our work establishes a molecular framework for understanding how temperature and ions jointly shape the structural ensemble of ssRNAs. Using coarse-grained simulations, we quantitatively reproduce experimental SAXS profiles (and *R*_*g*_) across a wide range of ion concentrations. Although ssRNAs are highly flexible and lack stable secondary structure, we show that base-stacking interactions govern the degree of right-handed helix formation, with rA_30_ forming the most persistent helical structure. Remarkably, while all sequences attract comparable numbers of ions, rU_30_ shows a slight preference for inner-sphere coordination, indicating sequence-dependent ion binding propensities. In contrast, the ion atmosphere around rA_30_ exhibits a small but detectable degree of chirality. These modest variations, though minor at the single-chain level, could enable binding partners to “sense”sequence identity through electrostatic or chiral recognition and are likely to be amplified in multichain contexts such as RNA condensate formation.

By incorporating a novel temperature-dependent Mg^2+^-phosphate potential into the simulations, we are able to illustrate the interplay between RNA structural ensemble and its ion atmosphere across temperatures. Our simulations reveal a sequence-specific collapse of RNAs at high temperature due to differences in base-stacking propensity. Each sequence exhibits a non-monotonic compaction, with distinct expansion temperature maxima that shift downward in the presence of Mg^2+^, underscoring the electrostatic origin of the transition. Such observations could be tested experimentally using recent advancements in variable-temperature SAXS measurements^130,131^ or other techniques. At high temperatures, transient helical structures are lost, rendering all sequences being equal. We show that Mg^2+^ binding increases slightly with temperature, consistent with the entropic nature of ion binding.^117,132^ As temperature rises, ions migrate from diffuse to inner-sphere regions, directly linking RNA compaction to dehydration and tighter phosphate coordination. Together, these findings highlight that the RNA ion atmosphere is not a passive screening layer but a dynamic, sequence-encoded extension of the RNA structure itself. By integrating counterion condensation, base stacking, and temperature-driven collapse within a unified framework, our study provides a molecular basis for understanding how simple RNA sequences undergo ion-mediated structural transitions and phase behaviors.

## Data Availability

All analysis and simulation code in this study are publicly available on GitHub (https://github.com/peter-zhang-chem/biophysical-temperature-dependent-ssRNA). The molecular dynamics trajectory data are not publicly archived, owing to their large size, but are available from the corresponding author upon reasonable request.

## Author Contributions Statement

H.Z. implemented the model and the computational framework, analyzed the data, and took the lead in writing the manuscript. H.M. assisted with data analysis and model validation. H.T.N. supervised the project, guided data interpretation, and revised the manuscript.

## Acknowledgments

We gratefully acknowledge financial support from the University at Buffalo to H.T.N.. Computational resources were provided by the Center for Computational Research at the University at Buffalo. We thank Peter Eastman for valuable discussions regarding the implementation of simulations in OpenMM.

