## Supplementary Information for "Temperature-Dependent Ion Migration Underlies Sequence-Specific Collapse of Unstructured RNA"

Heyang Zhang,[†] Hiranmay Maity,[†,‡] and Hung T. Nguyen[*,†,¶]

†*Department of Chemistry, State University of New York at Buffalo, Buffalo, NY, USA*

‡*Department of General Science, Birla Institute of Technology and Science, Pilani, Dubai Campus, Dubai International Academic City, Dubai 345055, UAE*

¶*Institute for Artificial Intelligence & Data Science, State University of New York at Buffalo, Buffalo, NY, USA*

### Analyses

All analysis scripts were written in Python 3.9.6 or Tcl 8.4.1 utilizing scientific libraries including Numpy (v1.26.4),[1] Pandas (v2.2.3),[2] SciPy (v1.13.1),[3] Seaborn (v0.13.2),[4] Matplotlib (v3.9.4),[5] and MDAnalysis (v2.7.0).[6,7] All scripts used in this study are available on GitHub: `https://github.com/peter-zhang-chem/biophysical-temperature-dependent-ssRNA`.

#### Autocorrelation Analysis

Normalized autocorrelation functions were computed to quantify temporal correlations in simulation observables. For a given time series $A(t)$, the correlation function was defined as

$$C(\tau) = \frac{\langle (A(t) - \langle A \rangle)(A(t+\tau) - \langle A \rangle) \rangle}{\langle (A(t) - \langle A \rangle)^2 \rangle}, \quad (1)$$

where $\tau$ is the lag time and angular brackets denote time averaging over the production trajectory. The autocorrelation functions were evaluated using a fast Fourier transform (FFT)-based approach and normalized such that $C(0) = 1$. Lag times were converted to physical units using a frame spacing of 20 ps. Correlation times were estimated from the decay of $C(\tau)$ to near zero, and were found to be on the order of a few hundred picoseconds for representative observables.

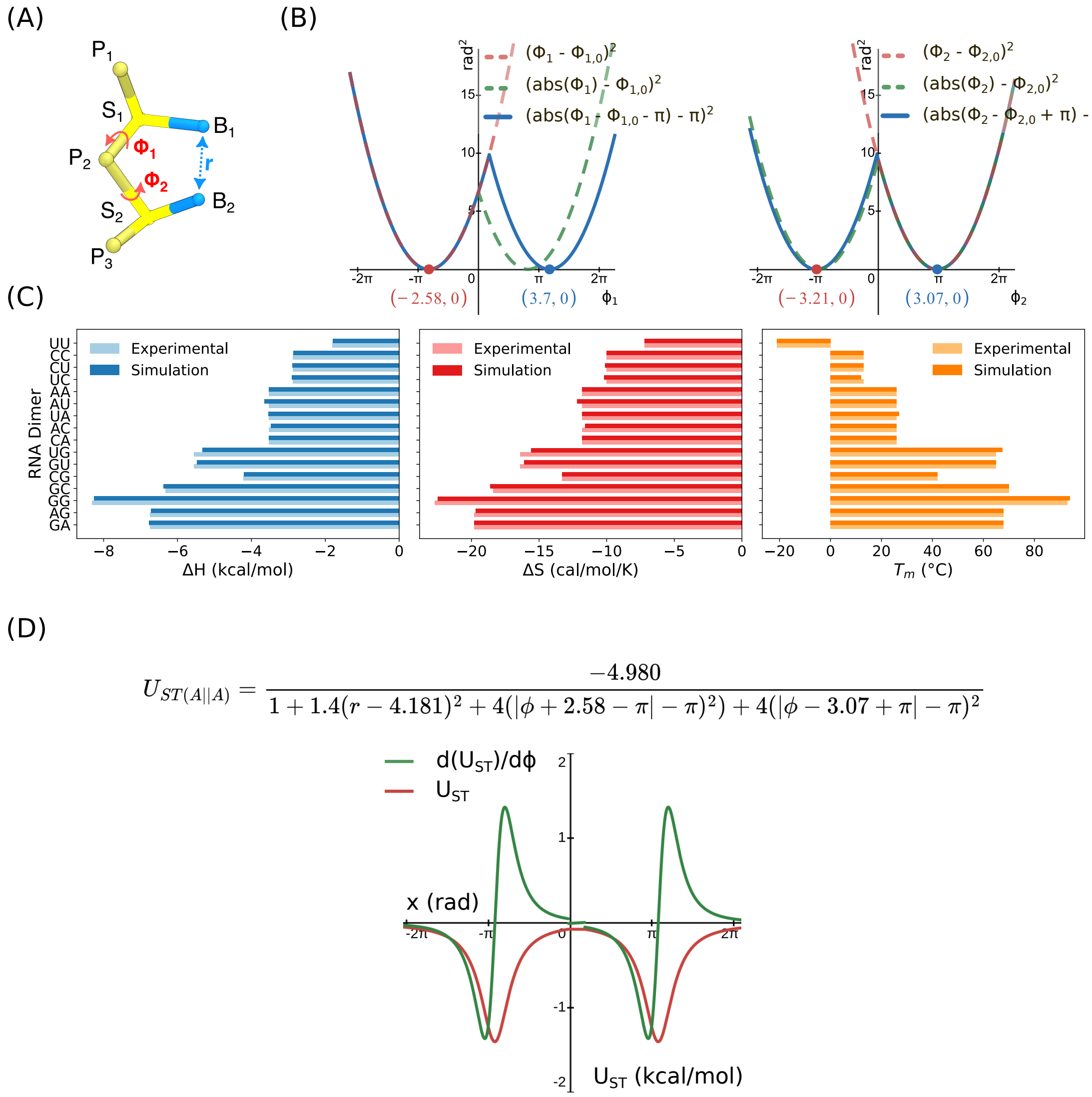

Figure S1: **Calibration of RNA stacking parameters.** **(A)** Illustration of the structural parameters defined in Eq. 3. **(B)** *Left:* Comparison of numerical evaluations of $(\phi_1 - \phi_{1,0})^2$ with $\phi_{1,0} = -2.587$ rad, which does not wrap continuously across 0 rad (red dashed line), leading to numerical instabilities during simulation. Replacing this term with $(|\phi_1| - \phi_{1,0})^2$ restores periodicity and resolves the instability. Further shifting the function to $(\phi_1 - \phi_{1,0} - \pi| - \pi)^2$ enforces a single minimum within $[-\pi, \pi]$ while preserving the overall functional shape. *Right:* Same analysis as the left panel, but for $\phi_2$ with $\phi_{2,0} = 3.071$ rad. **(C)** Calculated thermodynamic parameters reproduce values derived from the nearest-neighbor model following a previously reported procedure.[8,9] In this approach, duplex nearest-neighbor free energies are decomposed into contributions from base stacking along individual strands and hydrogen bonding between complementary bases, enabling extraction of effective stacking parameters for dinucleotides. **(D)** Final functional form of an example AA stacking interaction. The resulting potential has a single minimum within $[-\pi, \pi)$, consistent with the OpenMM dihedral domain. The resulting energy shows no numerical instability or energy drift during simulations despite discontinuities in the force near 0 rad.

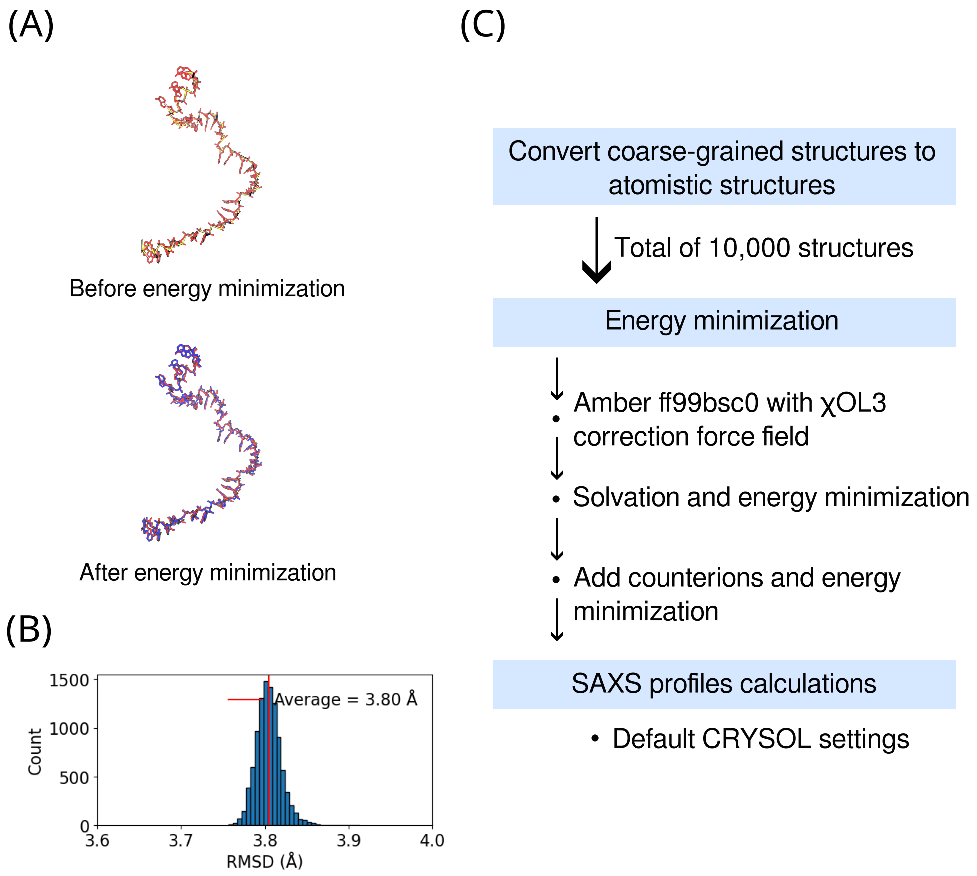

Figure S2: **(A)** Example of backmapping from coarse-grained coordinates to atomistic structure. The RMSD between structures before and after energy minimization is 3.8 Å, reflecting local relaxation required to remove steric clashes introduced during backmapping, while preserving the overall RNA geometry. **(B)** Distribution of RMSD values for 10,000 backmapped structures after minimization, showing an average of 3.8 Å, indicating that this value is representative of the ensemble. **(C)** Procedural outline from backmapping to SAXS profile calculations.

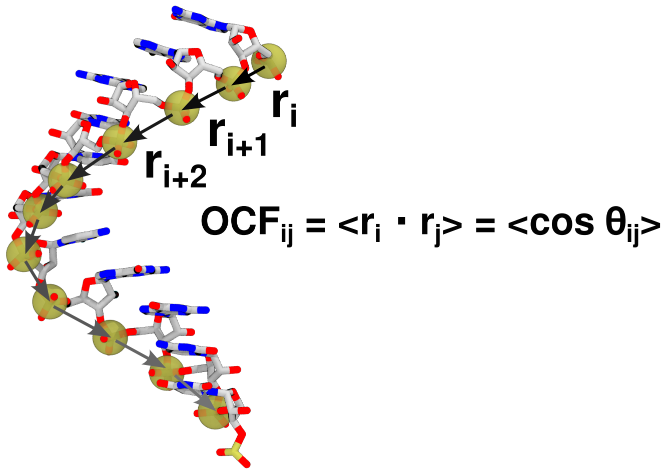

Figure S3: **Calculation of OCF using bond vectors between neighboring phosphates.** The OCF, defined as $\langle \cos\theta_{ij} \rangle$ quantifies the directional correlation between phosphate-phosphate bond vectors separated by $|i-j|$ along the ssRNA backbone.

Table S1: **Parameters for bond length in TIS RNA model**. This table presents the values of the spring constant ($k_b$) and the equilibrium bond distance ($r_0$). In the first column, B is nucleobase, P is phosphate, S is sugar, and P3 is 3' phosphate. The direction from 5' to 3' is indicated by an arrow. In the third column, A, G, C and U are the four nucleobases.

| $U_{BL} = k_b(r - r_0)^2$ | | | |
|---|---|---|---|
| 5' → 3' | $k_b$, kcal/mol/Å$^2$ | Pair | $r_0$, Å |
| B → S | 10 | A → S | 4.8515 |
| | | C → S | 4.2738 |
| | | G → S | 4.9659 |
| | | U → S | 4.2733 |
| P → S | 23 | | 3.8157 |
| S → P | 64 | | 4.601 |
| S → P3 | 64 | | 3.8157 |

Table S2: **Parameters for bond angle in TIS RNA model.** This table presents the values of the spring constant ($k_\theta$) and the equilibrium bond angle ($\theta_0$). In the first column, P is phosphate, S is sugar, (P+1) and (S+1) denote the phosphate and sugar of the next nucleotide, respectively, A, G, C and U are the four nucleobases.

| $U_{BA} = k_\theta(\theta - \theta_0)^2$ | | |
|---|---|---|
| Angle | $k_\theta$, kcal/mol/rad$^2$ | $\theta$, radian |
| A—S—(P+1) | 5 | 1.9259 |
| C—S—(P+1) | | 1.9655 |
| G—S—(P+1) | | 1.9150 |
| U—S—(P+1) | | 1.9663 |
| A—S—P | | 1.7029 |
| C—S—P | | 1.5803 |
| G—S—P | | 1.7690 |
| U—S—P | | 1.5735 |
| P—S—(P+1) | 20 | 1.4440 |
| S—(P+1)—(S+1) | | 1.5256 |

Table S3: **Parameters for excluded volume and electrostatic interactions.** Exception: if both interacting sites are nucleobase, we take $R_i + R_j = 3.2$ Å. The phosphate charge depends on [$Na^+$], [$Mg^{2+}$] and temperature. In the first column, P is phosphate, S is sugar, P3 is the 3' phosphate, A, G, C and U are the four nucleobases.

| | $\varepsilon$, kcal/mol | $R_i$, Å | $Q_i$ | Mass, Da |
|---|---|---|---|---|
| P | 1 | 1.89 | varies | 78.9596 |
| S | | 2.61 | 0 | 117.0557 |
| A | | 2.52 | 0 | 134.0472 |
| G | | 2.7 | 0 | 150.132 |
| C | | 2.43 | 0 | 110.036 |
| U | | 2.43 | 0 | 111.02 |
| $Mg^{2+}$ | | 2.00 | +2 | 24.305 |
| P3 | | 1.89 | 0 | 80.9742 |

Table S4: **Parameters $U_{ST}^0$ and $r_0$ for base stacking.** $T_m$ is the melting temperature for each nucleotide dimer, $k_B$ is the Boltzmann constant, h and s are adjustable parameters. In the first column, the 5' to 3' direction is shown by an arrow.

| $U_{ST}^0 = -h + k_B(T - T_m)s$ | | | | |
|---|---|---|---|---|
| 5' → 3' | h, kcal/mol | s | $T_m$, °C | $r_0$ (Å) |
| UU | 4.100 | -3.563 | -21 | 4.245 |
| CC | 4.700 | -1.567 | 13 | 4.250 |
| CU | 4.700 | -1.567 | 13 | 4.227 |
| UC | 4.690 | -1.567 | 13 | 4.268 |
| AA | 4.980 | -0.309 | 26 | 4.181 |
| AC | 4.970 | -0.700 | 26 | 3.826 |
| AU | 4.970 | -0.289 | 26 | 3.826 |
| CA | 4.940 | -0.309 | 26 | 4.701 |
| UA | 4.963 | -0.319 | 26 | 4.701 |
| CG | 5.450 | 0.300 | 42 | 4.979 |
| GU | 5.700 | 2.200 | 65 | 3.661 |
| UG | 5.700 | 2.200 | 65 | 4.998 |
| AG | 5.735 | 5.280 | 68 | 4.426 |
| GA | 5.732 | 5.240 | 68 | 4.013 |
| GC | 5.783 | 4.000 | 70 | 3.678 |
| GG | 6.198 | 7.346 | 93 | 4.243 |

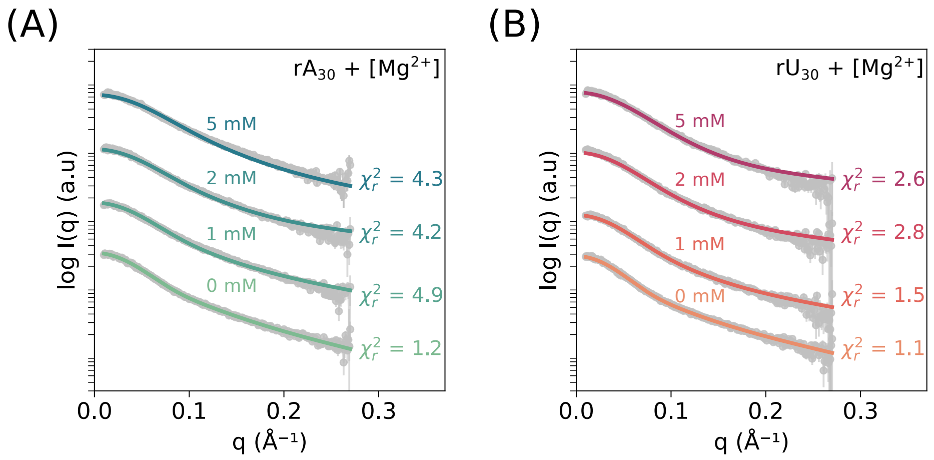

Figure S4: Experimental SAXS profiles (gray) compared with simulated scattering curves for **(A)** $rA_{30}$ and **(B)** $rU_{30}$ at 20 mM $Na^+$ with varying $[Mg^{2+}]$. Corresponding $\chi^2$ values between experiment and simulation are shown. The experimental SAXS data are available from the Small Angle Scattering Biological Data Bank (SASBDB) codes: SASDFA9, SASDFF9, SASDFG9, SASDFH9, SASDFJ9, SASDFP9, SASDFQ9, SASDFR9.

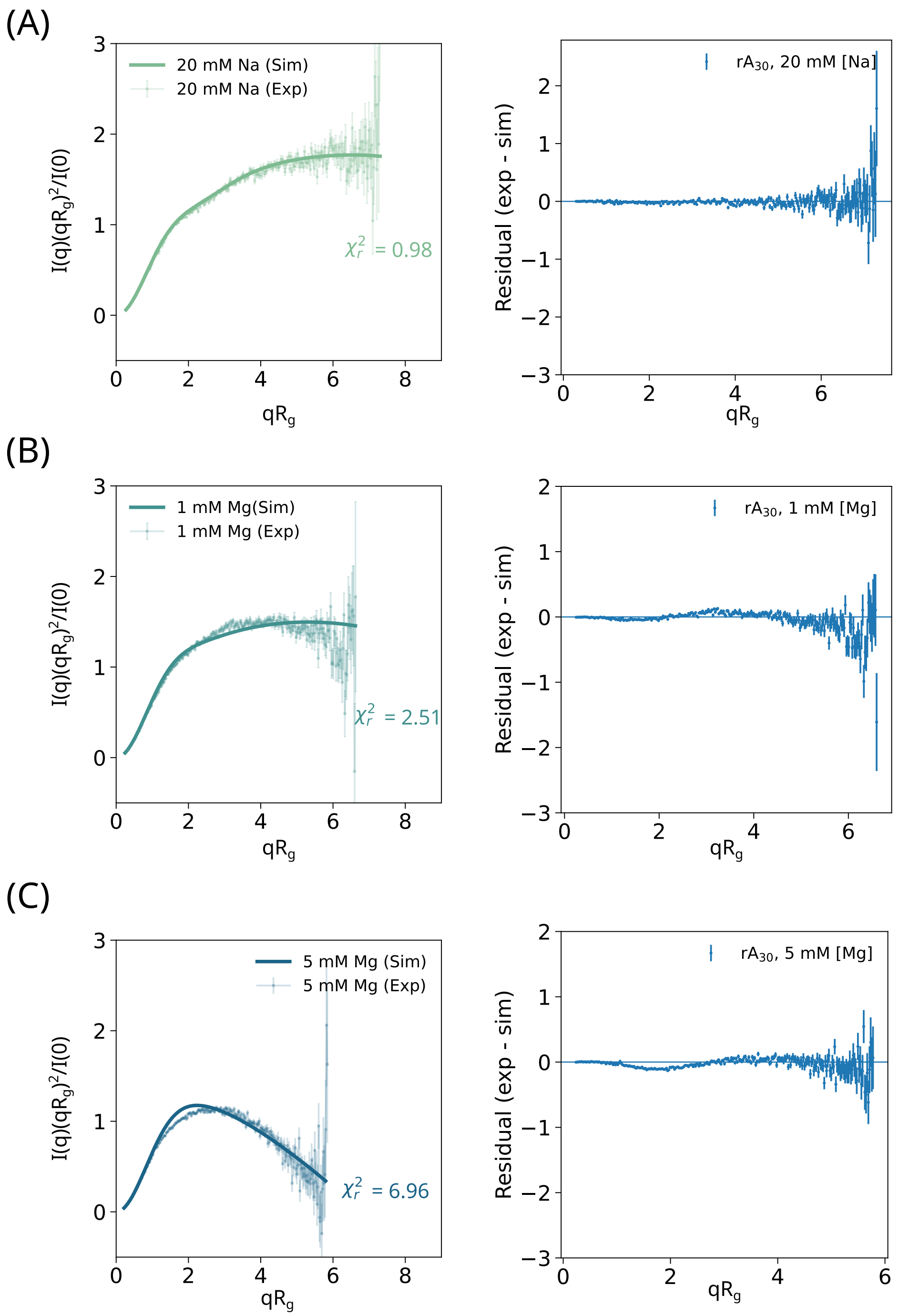

Figure S5: **(A)** Dimensionless Kratky plot of $rA_{30}$ calculated from simulations (solid lines) and compared with experimental data (points) in 20 mM $Na^+$ and 0 mM $Mg^{2+}$. The corresponding residuals (exp - sim) are shown on the right. **(B)** Same as (A), but in 1 mM $Mg^{2+}$ and 20 mM $Na^+$. **(C)** Same as (A), but in 5 mM $Mg^{2+}$ and 20 mM $Na^+$.

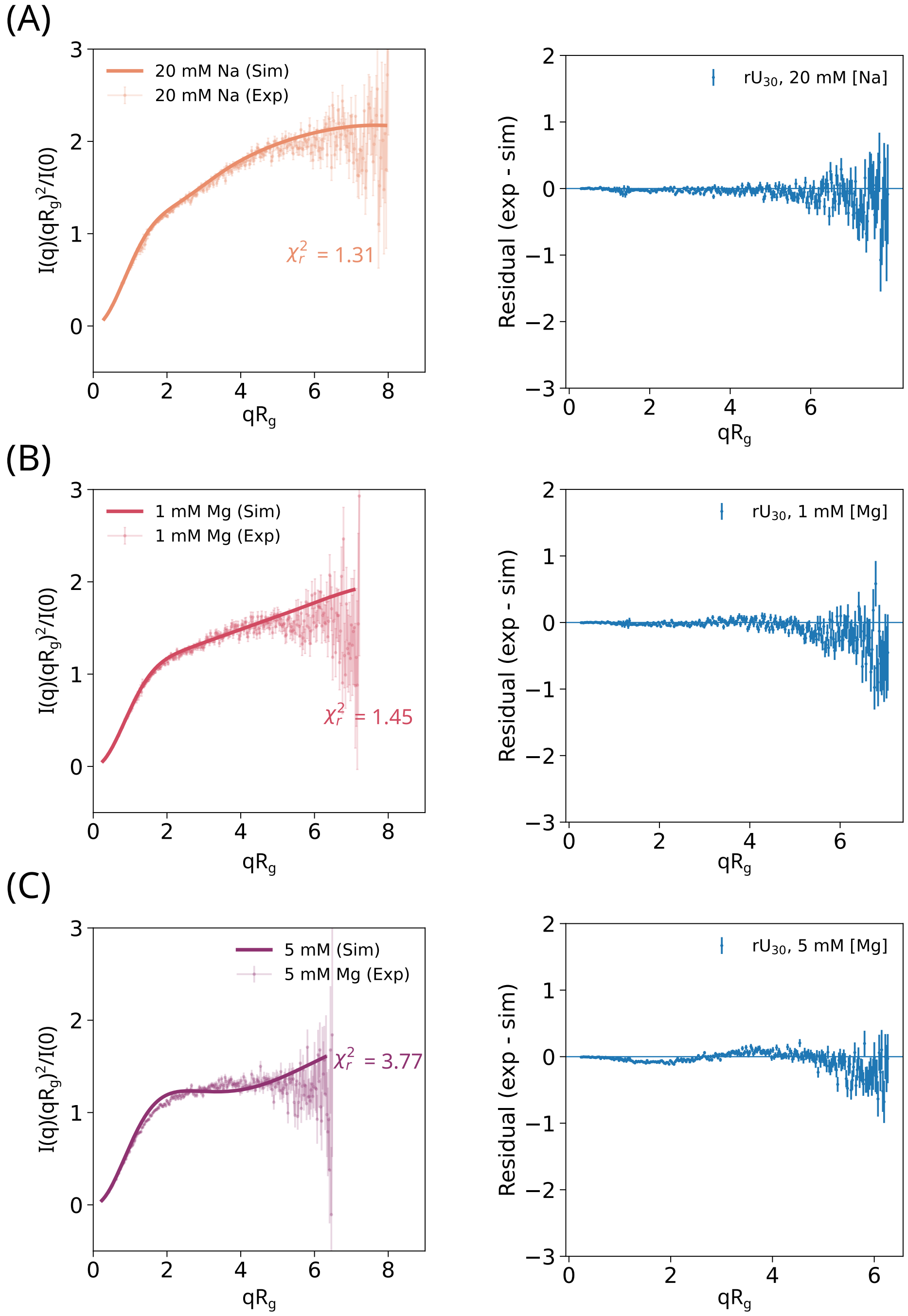

Figure S6: **(A)** Dimensionless Kratky plot of $rU_{30}$ calculated from simulations (solid line) and compared with experimental data (points) in 20 mM $Na^+$ and 0 mM $Mg^{2+}$. The corresponding residuals (exp - sim) are shown on the right. **(B)** Same as (A), but in 1 mM $Mg^{2+}$ and 20 mM $Na^+$. **(C)** Same as (A), but in 5 mM $Mg^{2+}$ and 20 mM $Na^+$.

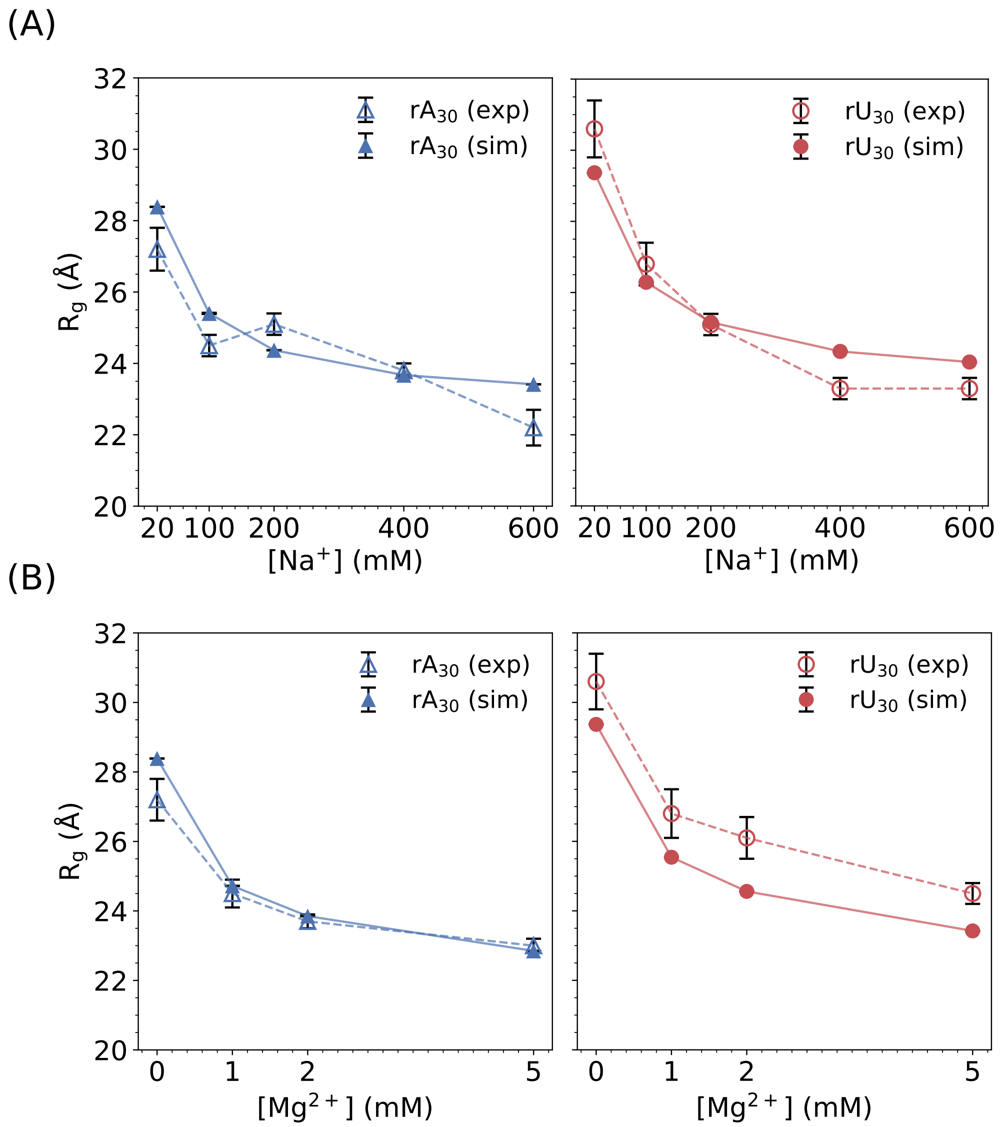

Figure S7: **(A)** Comparison between calculated radii of gyration ($R_g$) for $rA_{30}$ (left) $rU_{30}$ (right) with experimentally determined $R_g$ in 20 mM, 100 mM, 200 mM, 400 mM, and 600 mM $Na^+$. **(B)** Comparison between calculated radii of gyration ($R_g$) for $rA_{30}$ (left) $rU_{30}$ (right) with experimentally determined $R_g$ in 0 mM, 1 mM, 2 mM, and 5 mM $Mg^{2+}$ with 20 mM $Na^+$.

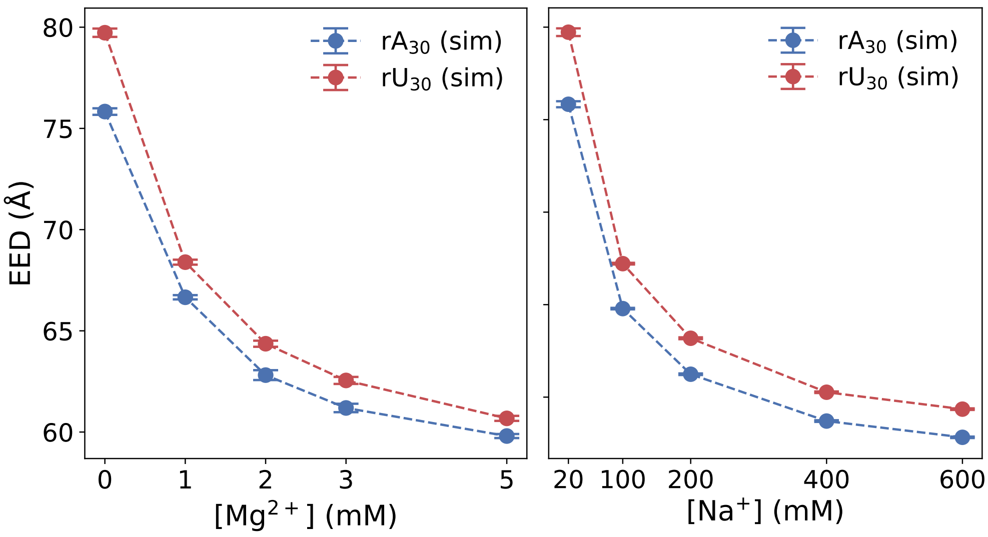

Figure S8: End-to-end distances comparison between rA$_{30}$ and rU$_{30}$ in various [Mg$^{2+}$] (left) and [Na$^+$] (right). rA$_{30}$ consistently exhibits shorter end-to-end distances than rU$_{30}$ across ionic conditions, reflecting stronger stacking propensity.

Table S5: **Tabulated values of $R_g$ from experiment and simulation**

| Ion Conc. | $rU_{30}$ (exp) (Å) | $rU_{30}$ (sim) (Å) | $rA_{30}$(exp) (Å) | $rA_{30}$ (sim) (Å) |
|---|---|---|---|---|
| 20 mM NaCl | $30.6 \pm 0.8$ | $29.37 \pm 0.01$ | $27.2 \pm 0.6$ | $28.39 \pm 0.01$ |
| 100 mM NaCl | $26.8 \pm 0.6$ | $26.29 \pm 0.02$ | $24.5 \pm 0.3$ | $25.40 \pm 0.01$ |
| 200 mM NaCl | $25.1 \pm 0.3$ | $25.16 \pm 0.01$ | $25.1 \pm 0.3$ | $24.37 \pm 0.01$ |
| 400 mM NaCl | $23.3 \pm 0.3$ | $24.34 \pm 0.01$ | $23.8 \pm 0.2$ | $23.67 \pm 0.01$ |
| 600 mM NaCl | $23.3 \pm 0.3$ | $24.05 \pm 0.01$ | $22.2 \pm 0.5$ | $23.42 \pm 0.01$ |
| 1 mM $MgCl_2$ | $26.8 \pm 0.7$ | $25.54 \pm 0.01$ | $24.5 \pm 0.4$ | $24.72 \pm 0.02$ |
| 2 mM $MgCl_2$ | $26.1 \pm 0.6$ | $24.56 \pm 0.03$ | $23.7 \pm 0.2$ | $23.85 \pm 0.02$ |
| 5 mM $MgCl_2$ | $24.5 \pm 0.3$ | $23.42 \pm 0.02$ | $23.0 \pm 0.2$ | $22.85 \pm 0.01$ |

Table S6: **Tabulated values of $\Gamma_{Mg}$ from experiment and simulation**

| Ion Conc. | $rU_{30}$ (exp) | $rU_{30}$ (sim) | $rA_{30}$ (exp) | $rA_{30}$ (sim) |
|---|---|---|---|---|
| 1 mM $MgCl_2$ | $6.16 \pm 0.08$ | $6.71 \pm 0.02$ | $7.80 \pm 0.07$ | $6.72 \pm 0.03$ |
| 2 mM $MgCl_2$ | $7.97 \pm 0.08$ | $8.63 \pm 0.01$ | $9.29 \pm 0.13$ | $8.63 \pm 0.01$ |
| 3 mM $MgCl_2$ | $8.97 \pm 0.13$ | $9.68 \pm 0.03$ | $10.46 \pm 0.16$ | $9.60 \pm 0.03$ |
| 5 mM $MgCl_2$ | N/A | $10.77 \pm 0.05$ | N/A | $10.82 \pm 0.01$ |

Table S7: **Tabulated values of end-to-end distances from simulation**

| Ion Conc. | $rA_{30}$ (sim) | $rU_{30}$ (sim) |
|---|---|---|
| 0 mM $MgCl_2$ | $75.83 \pm 0.16$ | $79.93 \pm 0.21$ |
| 1 mM $MgCl_2$ | $66.66 \pm 0.11$ | $68.39 \pm 0.12$ |
| 2 mM $MgCl_2$ | $62.81 \pm 0.24$ | $64.36 \pm 0.15$ |
| 3 mM $MgCl_2$ | $61.19 \pm 0.21$ | $62.55 \pm 0.17$ |
| 5 mM $MgCl_2$ | $59.80 \pm 0.10$ | $60.68 \pm 0.12$ |
| 20 mM NaCl | $75.83 \pm 0.16$ | $79.73 \pm 0.21$ |
| 100 mM NaCl | $64.79 \pm 0.04$ | $67.22 \pm 0.05$ |
| 200 mM NaCl | $61.24 \pm 0.04$ | $63.18 \pm 0.05$ |
| 400 mM NaCl | $58.71 \pm 0.04$ | $60.26 \pm 0.04$ |
| 600 mM NaCl | $57.83 \pm 0.04$ | $59.35 \pm 0.04$ |

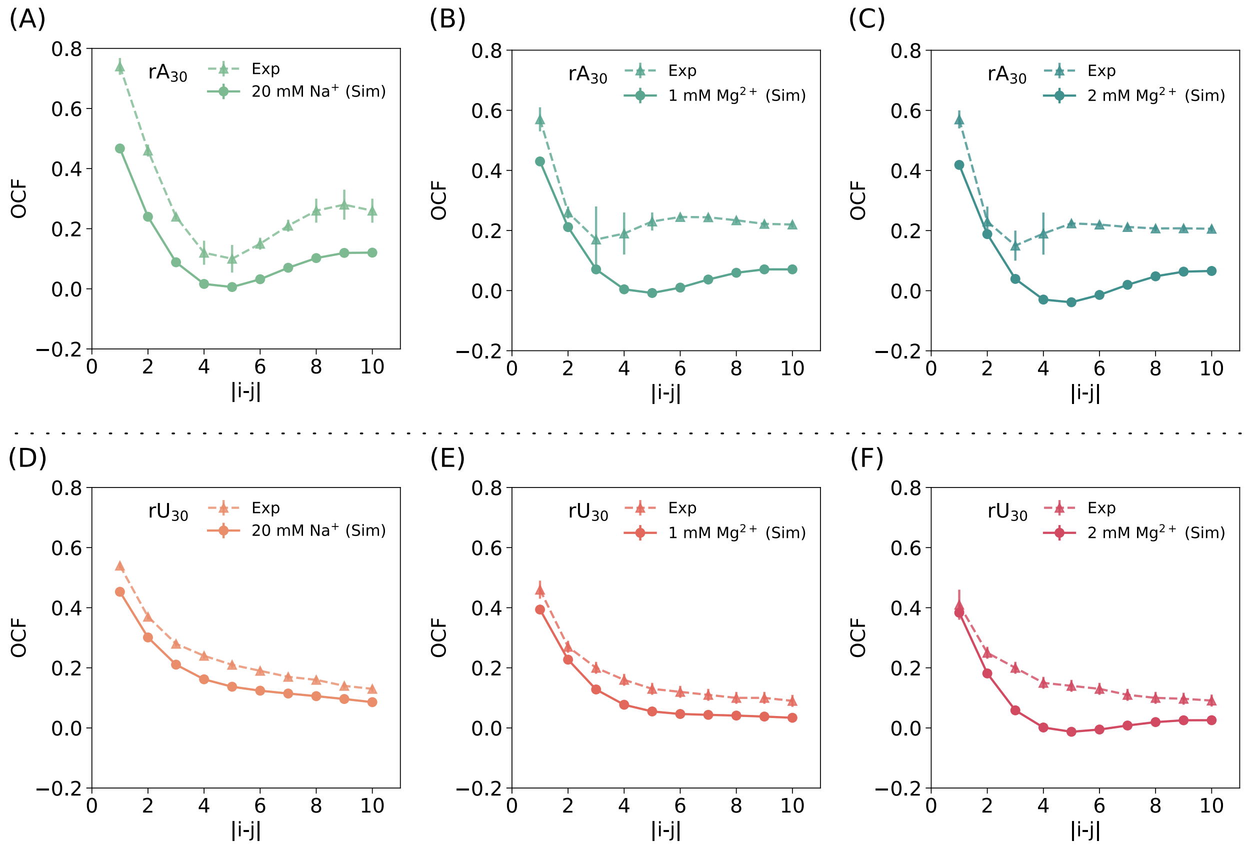

Figure S9: Individual comparison of simulated OCFs from this work and previously reported OCFs derived from experiment. **(A - C)** $rA_{30}$ in 20 mM $Na^+$, 1 mM, and 2 mM $Mg^{2+}$. **(D - F)** $rU_{30}$ under the same ionic conditions.

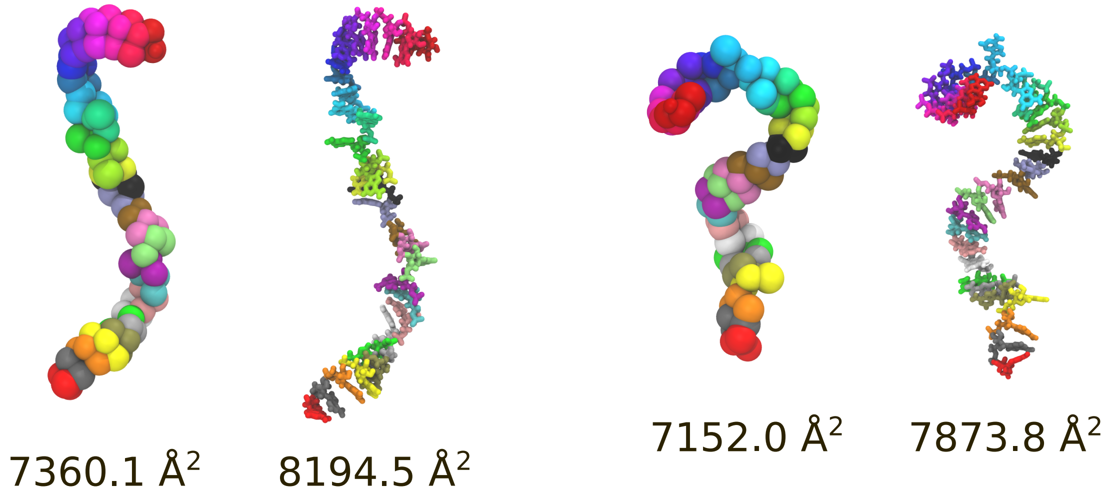

Figure S10: Representative comparison of solvent-accessible area (SASA) between coarse-grained (CG) and atomistic structures. SASA values derived from CG models are approximately 10% lower than those from atomistic simulations, likely due to surface smoothing inherent to the CG representation.

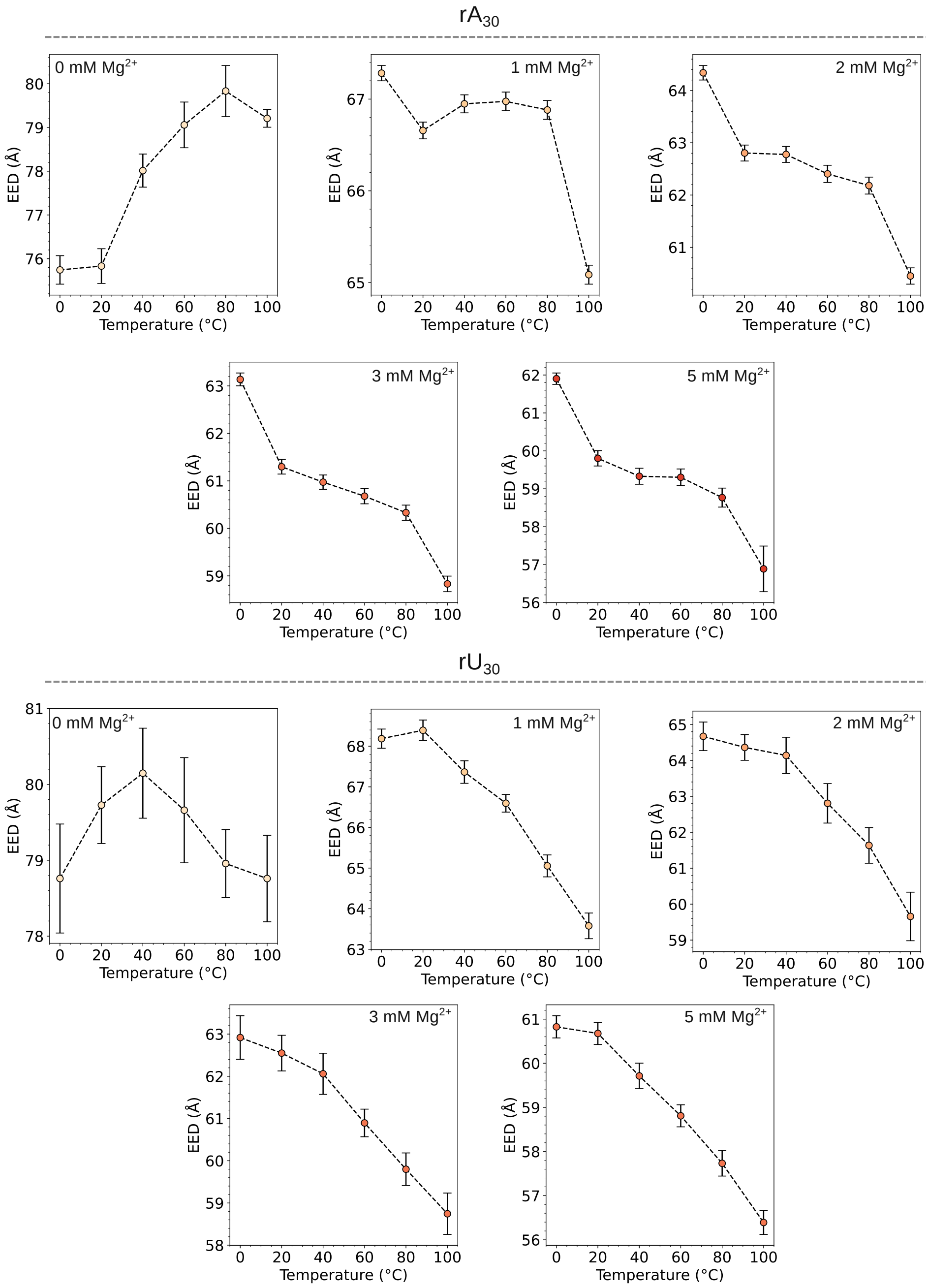

Figure S11: End-to-end distance of $rA_{30}$ (top) and $rU_{30}$ (bottom) as a function of temperature across varying [$Mg^{2+}$] concentrations. This analysis highlights the sequence- and ion-dependent thermal response of ssRNA chain extension.

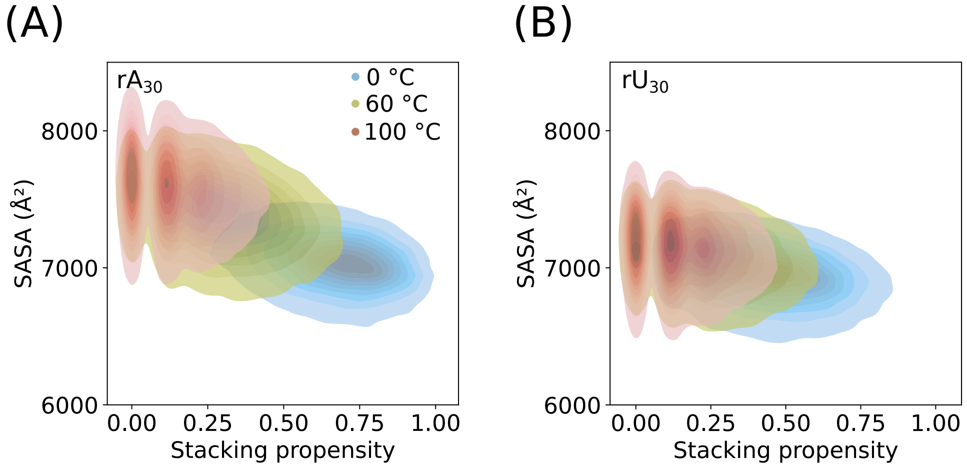

Figure S12: **(A)** Distribution of SASA *vs.* stacking propensity for $rA_{30}$ at 0 °C, 60 °C, and 100 °C. **(B)** Same as (A), but for $rU_{30}$. For both ssRNAs, the distributions shift toward lower stacking propensity and higher SASA with increasing temperature, with $rA_{30}$ exhibiting a more pronounced shift than $rU_{30}$.

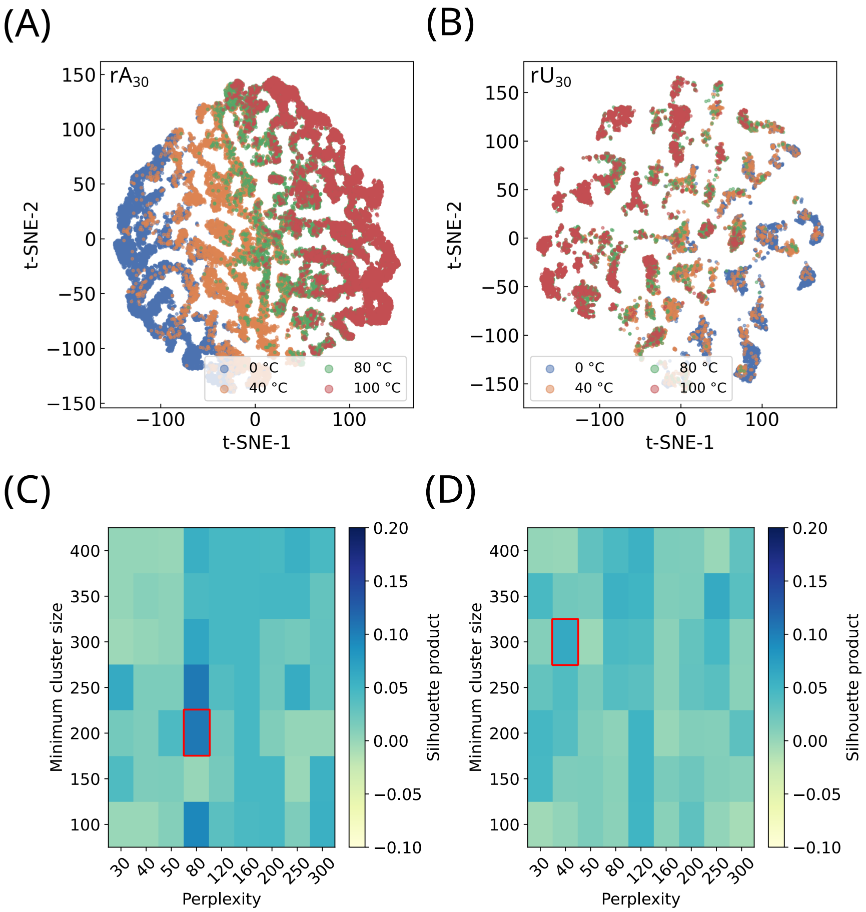

Figure S13: **(A)** t-SNE structural clustering of feature space consisting of $R_g$, end-to-end distance, SASA, potential energy of stacking, potential energy of Mg-P interactions for $rA_{30}$. **(B)** Same as (A) but for $rU_{30}$. The embeddings are used to qualitatively visualize structural similarity across ensembles at different temperatures. We emphasize that t-SNE primarily preserves local relationships and that the global arrangement of points should not be over-interpreted. In this context, $rA_{30}$ exhibits more pronounced separation of locally similar configurations across temperatures, whereas $rU_{30}$ shows greater overlap, suggesting comparatively weaker temperature-dependent changes in local structural organization. **(C, D)** Heatmaps of the Silhouette-product score used to evaluate hyperparameter performance across t-SNE perplexity and HDBSCAN minimum cluster size sweeps for $rA_{30}$ and $rU_{30}$, respectively. Red boxes mark the optimal parameter combination used to generate the embeddings in (A) and (B).

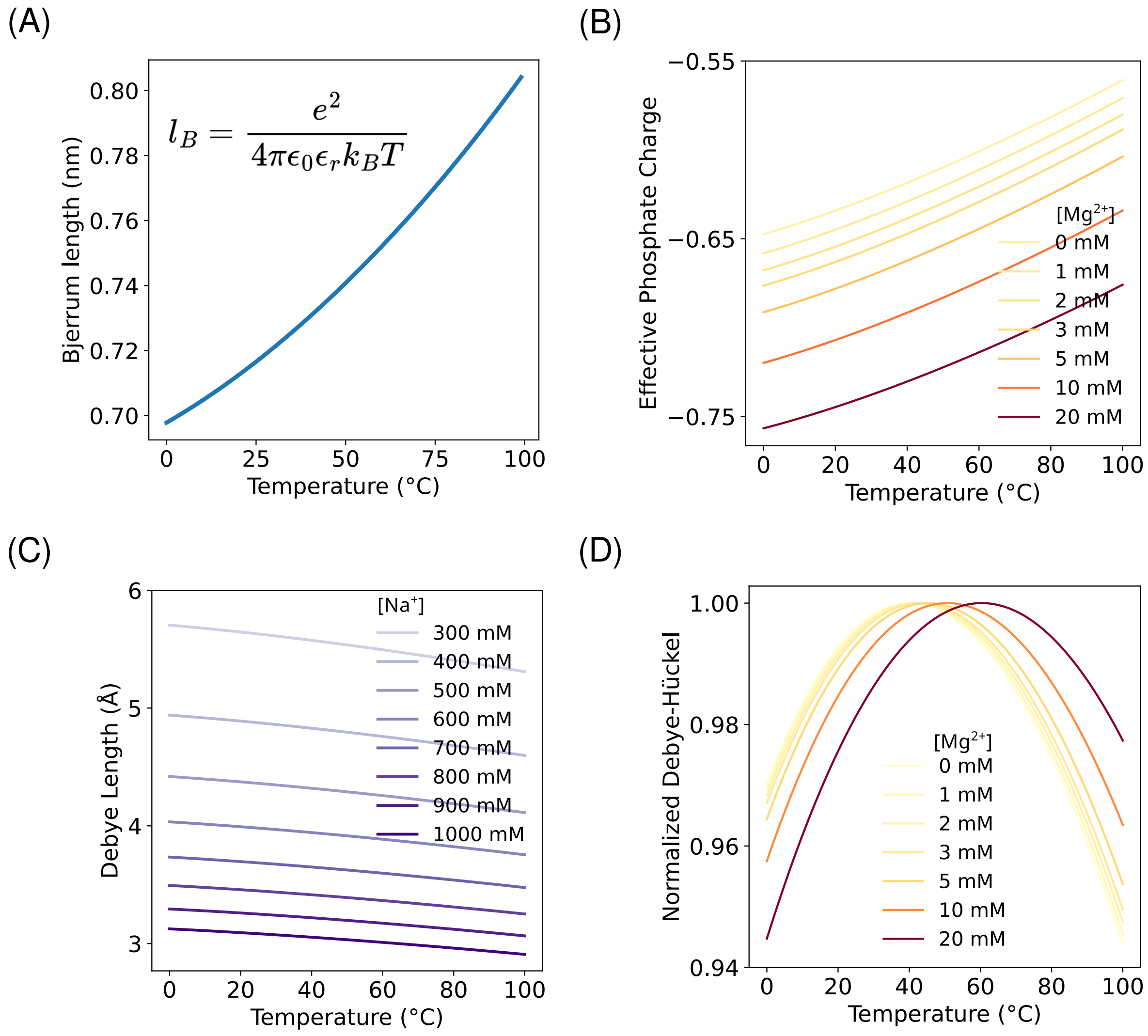

Figure S14: **(A)** Bjerrum length ($l_B$) in water as a function of temperature. **(B)** Phosphate charge becomes smaller at high temperature due to stronger monovalent ion adsorption. Increasing $Mg^{2+}$ concentration releases monovalent ions to the buffer, increasing the effective P charge, and thus enhancing Mg-P interactions. **(C)** Debye length slightly decreases with temperature, reflecting reduced electrostatic screening length at high temperature. **(D)** Normalized Debye–Hückel interaction potential as a function of temperature, rising at low temperature due to enhanced electrostatic attractions, then decreasing at high temperature as screening strengths and backbone charges are progressively neutralized.

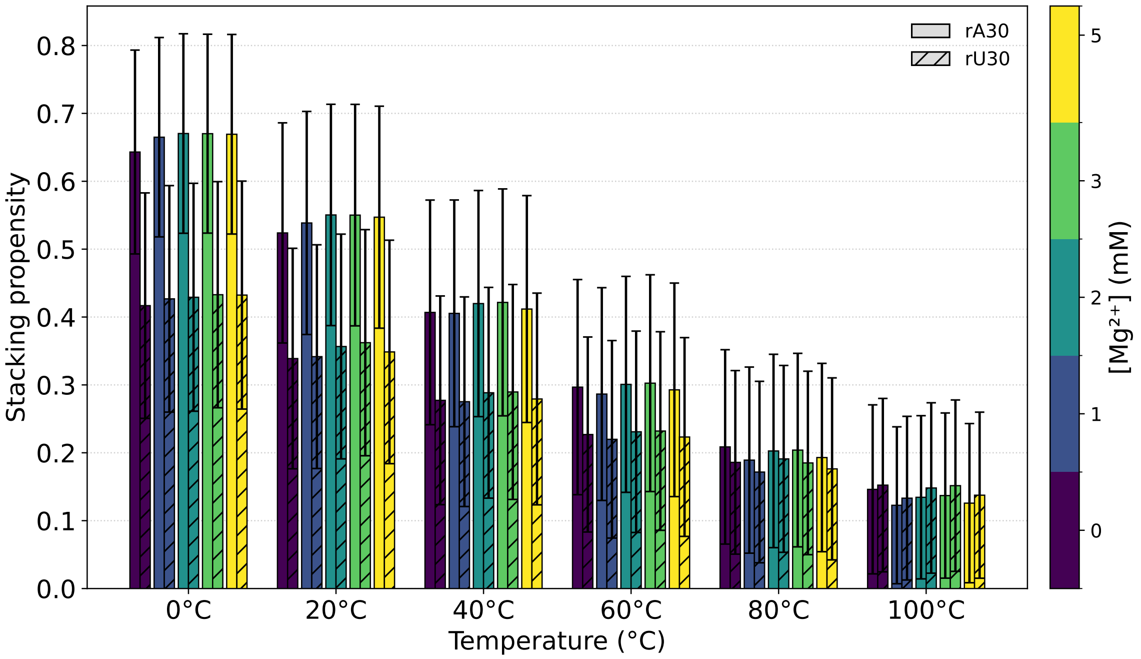

Figure S15: Stacking propensity as a function of temperature for $rA_{30}$ and $rU_{30}$ in various $Mg^{2+}$ conditions. Error bars represent the standard deviation.

Table S8: Temperature dependence of $R_g$ of $rA_{30}$ at varying $Mg^{2+}$ concentrations. Uncertainties are reported as the standard error of the mean (SEM) from block averaging.

| **$rA_{30}$** | | |
|---|---|---|
| $[Mg^{2+}]$ (mM) | Temperature (°C) | Average $R_g \pm$ SEM (Å) |
| 0 | 0 | $27.19 \pm 0.03$ |
| 0 | 20 | $28.17 \pm 0.04$ |
| 0 | 40 | $28.94 \pm 0.04$ |
| 0 | 60 | $29.27 \pm 0.04$ |
| 0 | 80 | $29.39 \pm 0.04$ |
| 0 | 100 | $29.19 \pm 0.04$ |
| 1 | 0 | $24.62 \pm 0.02$ |
| 1 | 20 | $25.31 \pm 0.02$ |
| 1 | 40 | $25.80 \pm 0.02$ |
| 1 | 60 | $25.88 \pm 0.02$ |
| 1 | 80 | $25.80 \pm 0.02$ |
| 1 | 100 | $25.23 \pm 0.02$ |
| 2 | 0 | $23.76 \pm 0.03$ |
| 2 | 20 | $24.27 \pm 0.04$ |
| 2 | 40 | $24.65 \pm 0.04$ |
| 2 | 60 | $24.70 \pm 0.03$ |
| 2 | 80 | $24.54 \pm 0.04$ |
| 2 | 100 | $24.00 \pm 0.04$ |
| 3 | 0 | $23.43 \pm 0.03$ |
| 3 | 20 | $23.82 \pm 0.04$ |
| 3 | 40 | $24.18 \pm 0.03$ |
| 3 | 60 | $24.24 \pm 0.03$ |
| 3 | 80 | $24.07 \pm 0.04$ |
| 3 | 100 | $23.67 \pm 0.03$ |
| 5 | 0 | $23.10 \pm 0.01$ |
| 5 | 20 | $23.44 \pm 0.02$ |
| 5 | 40 | $23.78 \pm 0.02$ |
| 5 | 60 | $23.88 \pm 0.02$ |
| 5 | 80 | $23.70 \pm 0.02$ |
| 5 | 100 | $23.07 \pm 0.04$ |

Table S9: Temperature dependence of $R_g$ of $rU_{30}$ at varying $Mg^{2+}$ concentrations. Uncertainties are reported as the standard error of the mean (SEM) from block averaging.

| **$rU_{30}$** | | |
|---|---|---|
| $[Mg^{2+}]$ (mM) | Temperature (°C) | Average $R_g \pm$ SEM (Å) |
| 0 | 0 | $29.04 \pm 0.04$ |
| 0 | 20 | $29.38 \pm 0.04$ |
| 0 | 40 | $29.45 \pm 0.04$ |
| 0 | 60 | $29.39 \pm 0.04$ |
| 0 | 80 | $29.22 \pm 0.04$ |
| 0 | 100 | $29.08 \pm 0.04$ |
| 1 | 0 | $26.06 \pm 0.02$ |
| 1 | 20 | $26.19 \pm 0.02$ |
| 1 | 40 | $26.01 \pm 0.03$ |
| 1 | 60 | $25.75 \pm 0.03$ |
| 1 | 80 | $25.27 \pm 0.03$ |
| 1 | 100 | $24.80 \pm 0.03$ |
| 2 | 0 | $25.13 \pm 0.05$ |
| 2 | 20 | $25.21 \pm 0.04$ |
| 2 | 40 | $25.13 \pm 0.04$ |
| 2 | 60 | $24.74 \pm 0.05$ |
| 2 | 80 | $24.40 \pm 0.05$ |
| 2 | 100 | $23.83 \pm 0.05$ |
| 3 | 0 | $24.65 \pm 0.04$ |
| 3 | 20 | $24.70 \pm 0.05$ |
| 3 | 40 | $24.59 \pm 0.03$ |
| 3 | 60 | $24.26 \pm 0.04$ |
| 3 | 80 | $23.89 \pm 0.04$ |
| 3 | 100 | $23.46 \pm 0.04$ |
| 5 | 0 | $24.06 \pm 0.02$ |
| 5 | 20 | $24.17 \pm 0.02$ |
| 5 | 40 | $23.99 \pm 0.02$ |
| 5 | 60 | $23.73 \pm 0.02$ |
| 5 | 80 | $23.39 \pm 0.02$ |
| 5 | 100 | $22.96 \pm 0.02$ |

Table S10: Temperature dependence of $\Gamma_{Mg}$ of $rA_{30}$ at varying $Mg^{2+}$ concentrations. Uncertainties are reported as the standard deviation (SD) from block averaging.

| **$rA_{30}$** | | |
|---|---|---|
| $[Mg^{2+}]$ (mM) | Temperature (°C) | Average $\Gamma_{Mg}$ ± SD |
| 1 | 0 | $6.66 \pm 0.04$ |
| 1 | 20 | $6.70 \pm 0.03$ |
| 1 | 40 | $6.84 \pm 0.04$ |
| 1 | 60 | $7.05 \pm 0.02$ |
| 1 | 80 | $7.17 \pm 0.03$ |
| 1 | 100 | $7.34 \pm 0.09$ |
| 2 | 0 | $8.51 \pm 0.06$ |
| 2 | 20 | $8.62 \pm 0.07$ |
| 2 | 40 | $8.79 \pm 0.04$ |
| 2 | 60 | $8.89 \pm 0.03$ |
| 2 | 80 | $8.95 \pm 0.05$ |
| 2 | 100 | $9.08 \pm 0.05$ |
| 3 | 0 | $9.48 \pm 0.02$ |
| 3 | 20 | $9.63 \pm 0.01$ |
| 3 | 40 | $9.80 \pm 0.02$ |
| 3 | 60 | $9.83 \pm 0.01$ |
| 3 | 80 | $9.95 \pm 0.03$ |
| 3 | 100 | $10.05 \pm 0.02$ |
| 5 | 0 | $10.65 \pm 0.13$ |
| 5 | 20 | $10.74 \pm 0.10$ |
| 5 | 40 | $10.85 \pm 0.08$ |
| 5 | 60 | $10.96 \pm 0.04$ |
| 5 | 80 | $11.02 \pm 0.08$ |
| 5 | 100 | $11.13 \pm 0.04$ |

Table S11: Temperature Dependence of $\Gamma_{Mg}$ of $rU_{30}$ at varying $Mg^{2+}$ concentrations. Uncertainties are reported as the standard deviation (SD) from block averaging.

| **$rU_{30}$** | | |
|---|---|---|
| $[Mg^{2+}]$ (mM) | Temperature (°C) | Average $\Gamma_{Mg}$ ± SD |
| 1 | 0 | $6.59 \pm 0.06$ |
| 1 | 20 | $6.71 \pm 0.06$ |
| 1 | 40 | $6.95 \pm 0.04$ |
| 1 | 60 | $7.17 \pm 0.02$ |
| 1 | 80 | $7.28 \pm 0.06$ |
| 1 | 100 | $7.33 \pm 0.05$ |
| 2 | 0 | $8.55 \pm 0.05$ |
| 2 | 20 | $8.64 \pm 0.04$ |
| 2 | 40 | $8.79 \pm 0.03$ |
| 2 | 60 | $8.92 \pm 0.03$ |
| 2 | 80 | $8.99 \pm 0.03$ |
| 2 | 100 | $9.16 \pm 0.02$ |
| 3 | 0 | $9.54 \pm 0.01$ |
| 3 | 20 | $9.67 \pm 0.01$ |
| 3 | 40 | $9.72 \pm 0.03$ |
| 3 | 60 | $9.97 \pm 0.02$ |
| 3 | 80 | $9.98 \pm 0.01$ |
| 3 | 100 | $10.10 \pm 0.03$ |
| 5 | 0 | $10.72 \pm 0.09$ |
| 5 | 20 | $10.77 \pm 0.07$ |
| 5 | 40 | $10.89 \pm 0.07$ |
| 5 | 60 | $10.96 \pm 0.09$ |
| 5 | 80 | $11.13 \pm 0.05$ |
| 5 | 100 | $11.19 \pm 0.09$ |

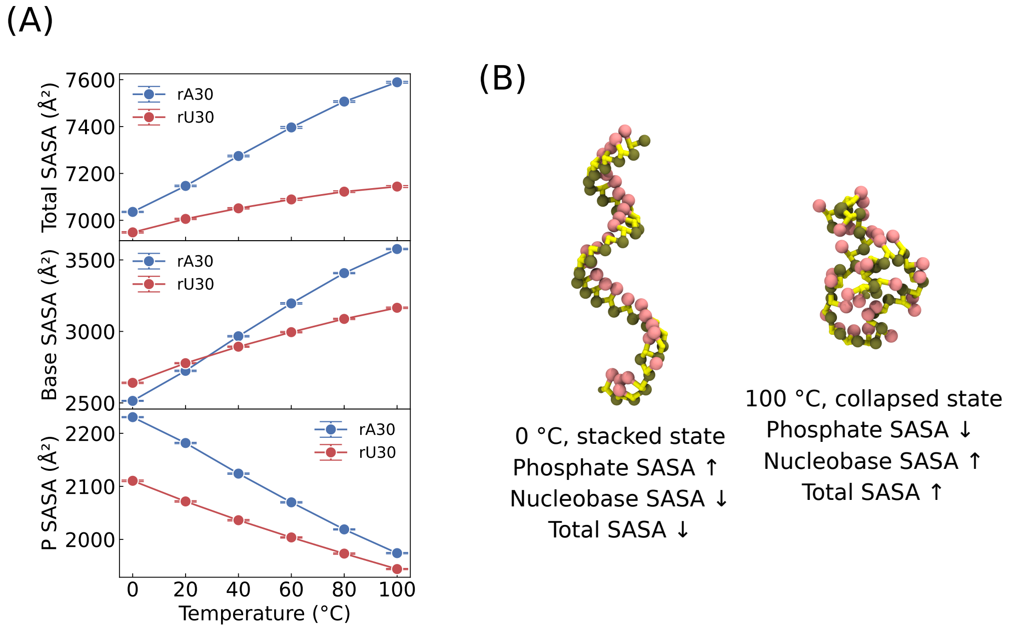

Figure S16: **(A)** SASA of $rA_{30}$ and $rU_{30}$ in 5 mM $Mg^{2+}$ as a function of temperature. **Top:** Total SASA increases with temperature, and the difference between $rA_{30}$ and $rU_{30}$ increases as a function of increasing temperature. **Middle:** Nucleobase SASA increases with temperature; $rA_{30}$ increases at a faster rate and surpasses $rU_{30}$ at higher temperatures. **Bottom:** Phosphate SASA decreases as temperature increases, and $rA_{30}$ decreases at a faster rate than $rU_{30}$. **(B)** Illustration of $rA_{30}$ at 0 °C with nucleobases in stacked conformation resulting in lower nucleobase and total SASA. On the other hand, $rA_{30}$ at 100 °C collapses with broken stacking resulting in higher nucleobase and total SASA.

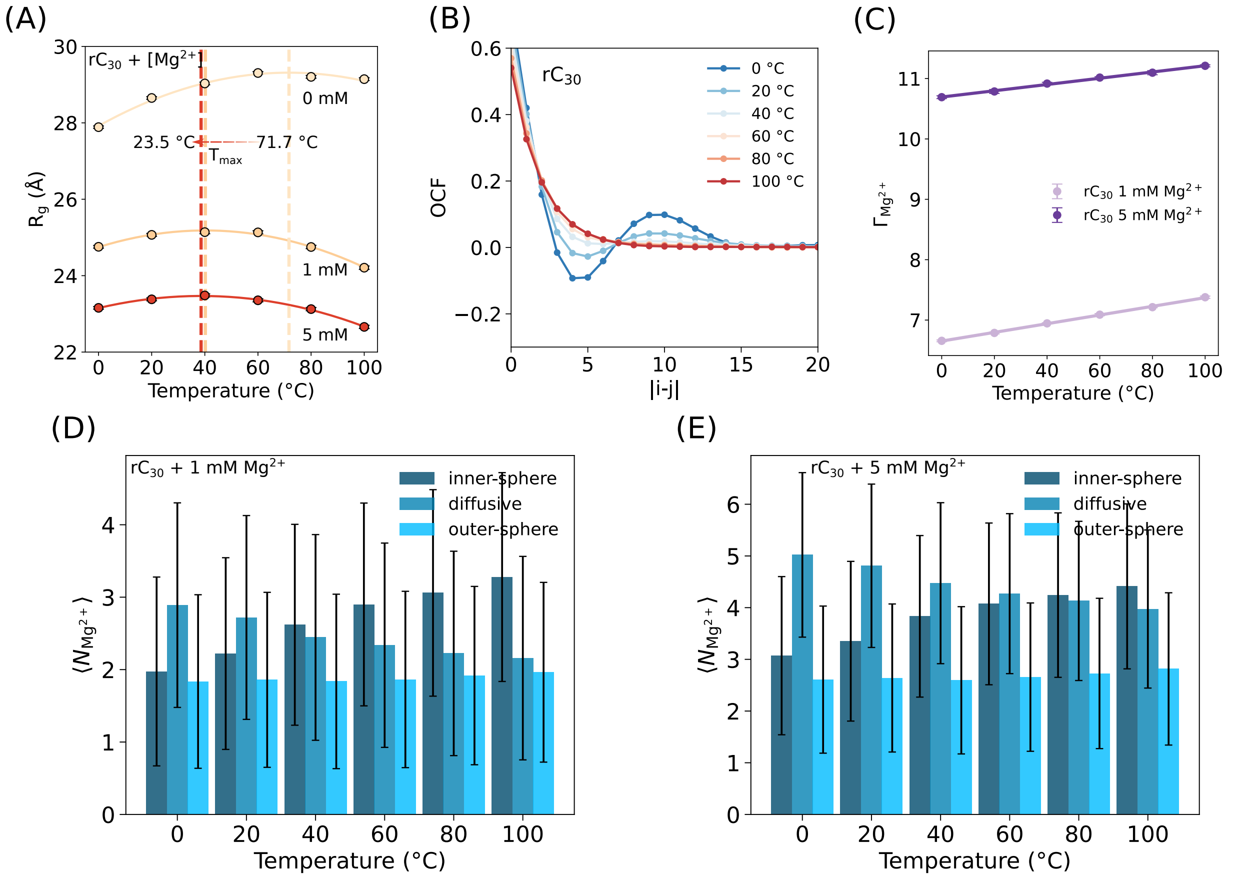

Figure S17: **(A)** $R_g$ of $rC_{30}$ as a function of temperature in 0, 1, and 5 mM $Mg^{2+}$. Vertical dashed lines denote $T_{max}$, the temperature of maximum expansion. **(B)** Orientational correlation functions (OCFs) from 0 °C to 100 °C for $rC_{30}$. **(C)** Preferential interaction coefficient for $Mg^{2+}$ ($\Gamma_{Mg}$) increases linearly with temperature for $rC_{30}$ at 1 mM (light purple) and 5 mM (dark purple) $Mg^{2+}$. **(D)** Ion atmosphere composition from 0 to 100 °C of $rC_{30}$ in 1 mM $Mg^{2+}$. **(E)** Same as (D), but for $rC_{30}$ in 5 mM $Mg^{2+}$.

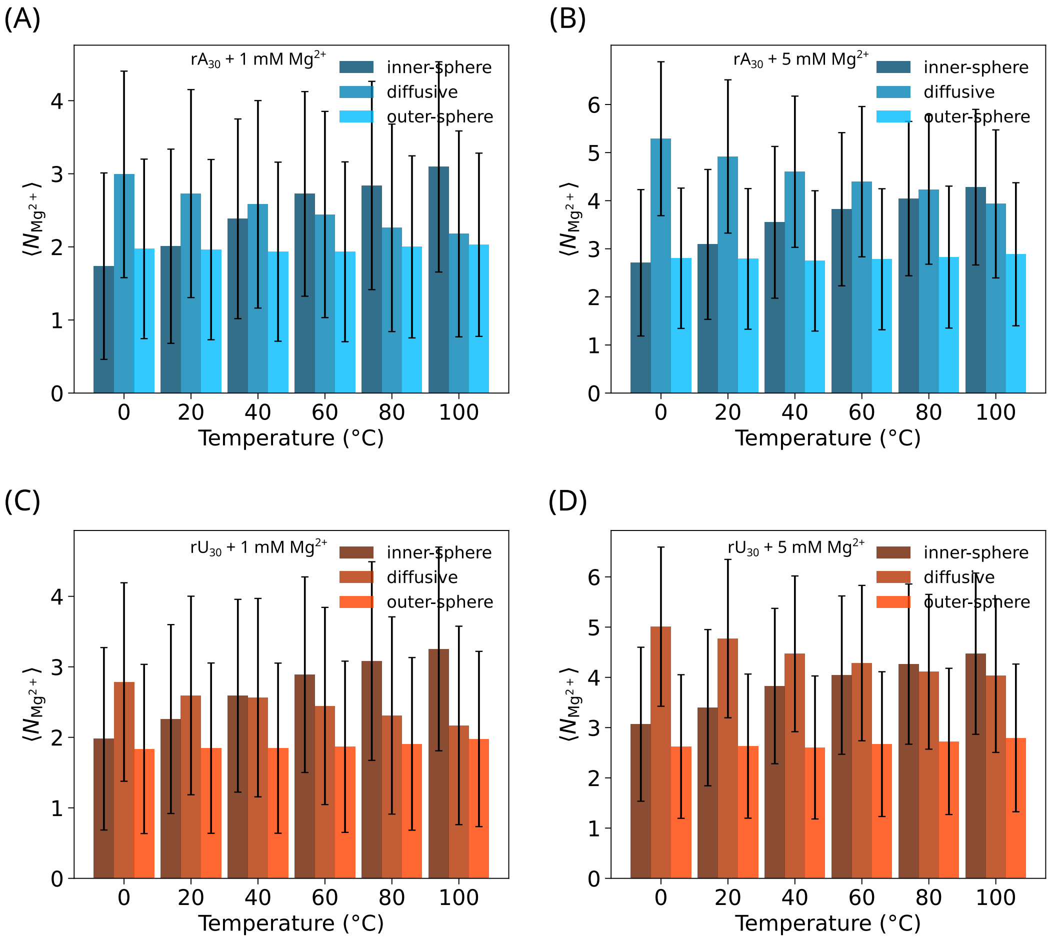

Figure S18: **(A)** Ion atmosphere composition from 0 to 100 °C of $rA_{30}$ in 1 mM $Mg^{2+}$ and 20 mM $Na^{+}$. **(B)** Same as (A), but for $rA_{30}$ in 5 mM $Mg^{2+}$ and 20 mM $Na^{+}$. **(C)** Ion atmosphere composition from 0 to 100 °C of $rU_{30}$ in 1 mM $Mg^{2+}$ and 20 mM $Na^{+}$. **(D)** Same as (C), but for $rU_{30}$ in 5 mM $Mg^{2+}$ and 20 mM $Na^{+}$ .

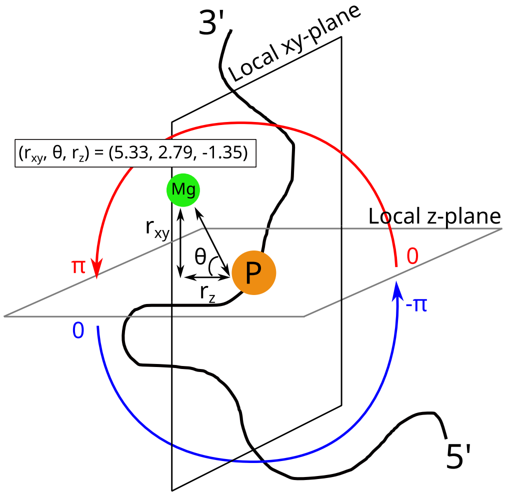

Figure S19: **Schematic of the local coordinate system used for ion atmosphere projection.** For each phosphate (P), a local frame is defined with the origin at the phosphate and the z-axis aligned with the RNA principal axis (5' to 3' direction). The position of a $Mg^{2+}$ ion is described by the in-plane distance $r_{xy}$, azimuthal angle $\theta$, and out-of-plane displacement $r_z$. This local representation ensures that ion positions are characterized relative to the phosphate, independent of global RNA conformation.

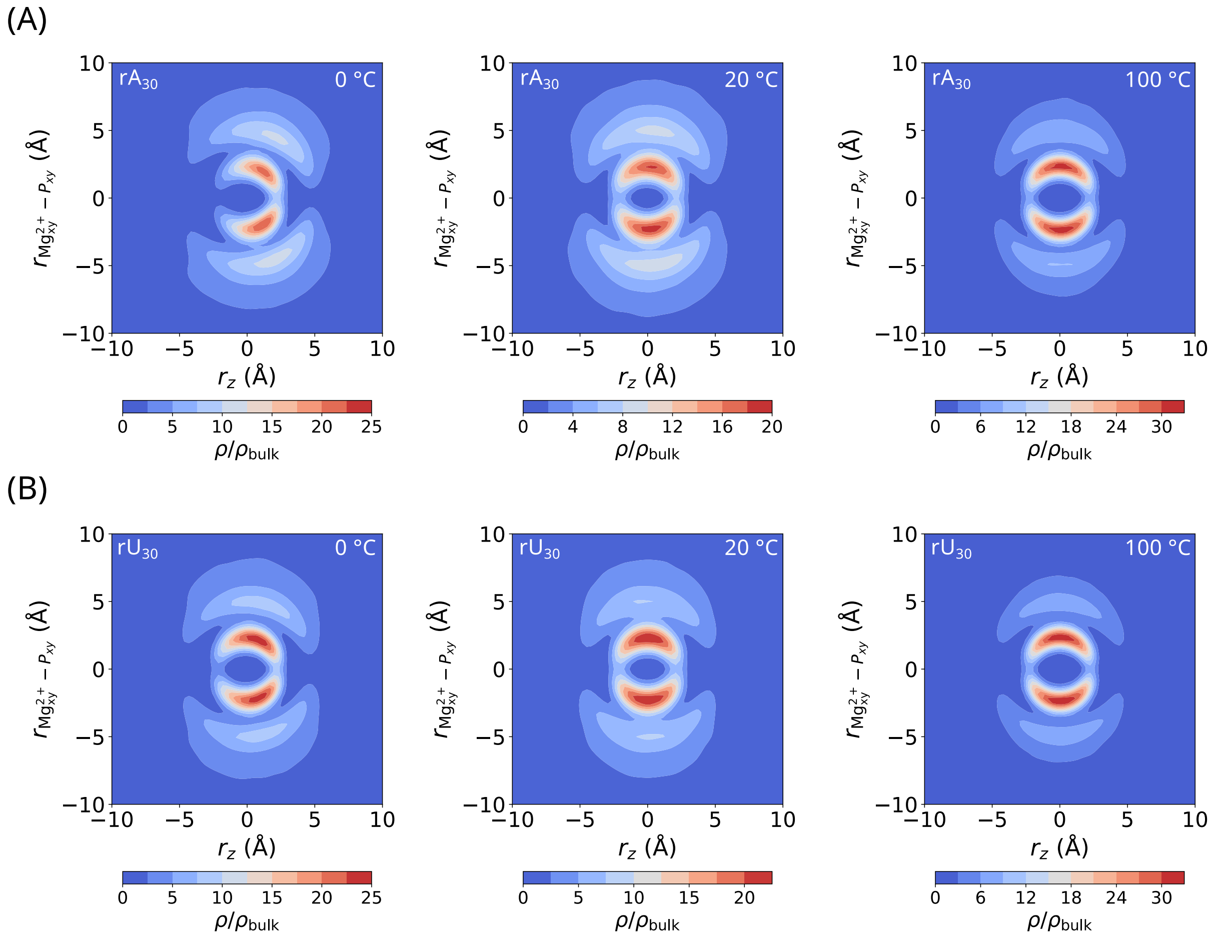

Figure S20: **(A)** Ion atmosphere projections of $rA_{30}$ at 0 °C, 20 °C, and 100 °C (left to right). **(B)** Same as (A), but for $rU_{30}$. As temperature increases, the right-skewed distribution arising from helical structures disappears, and the ion atmosphere collapses as diffusive ions migrate into the inner-sphere region.

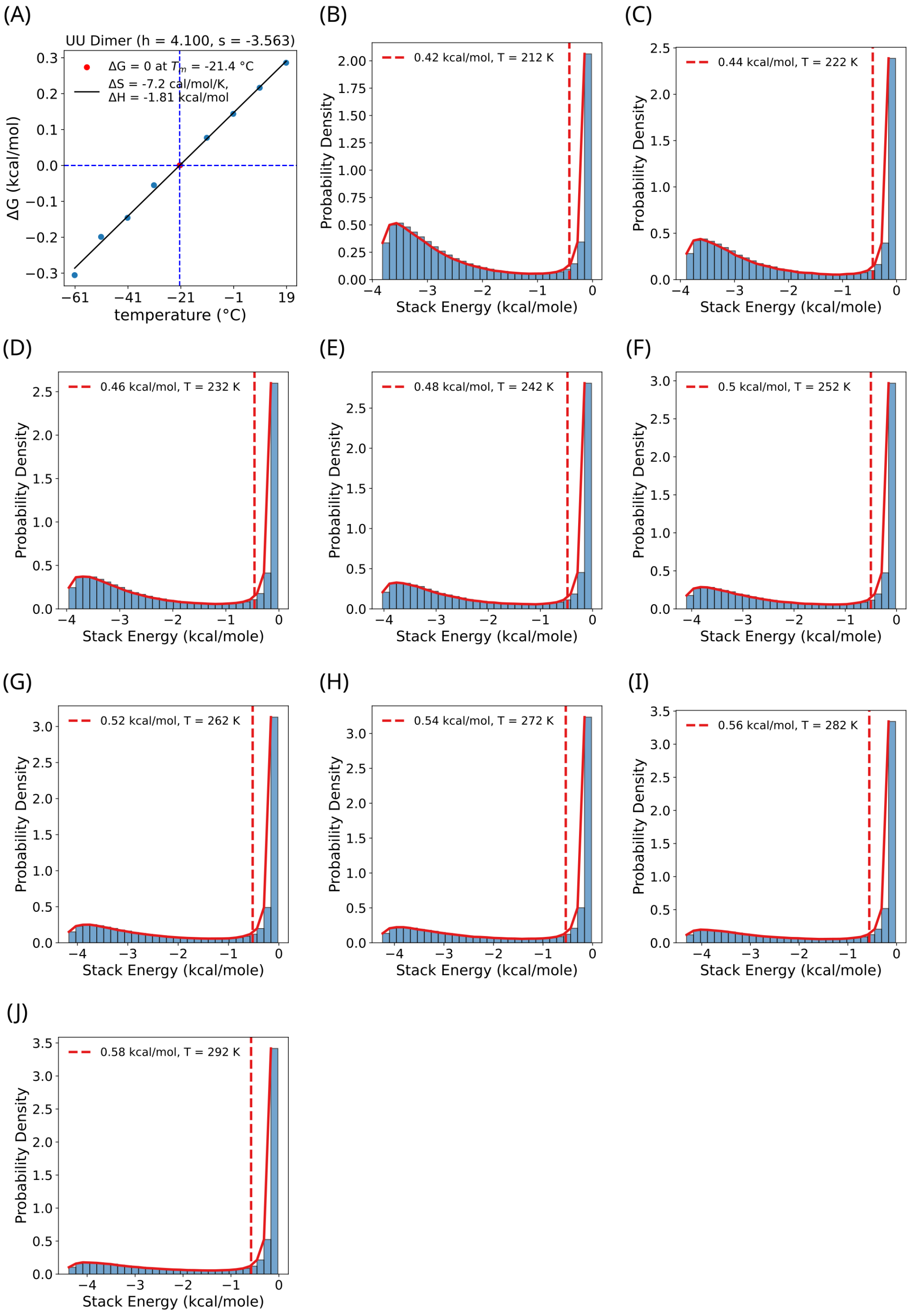

Figure S21: **(A)** Free energy of stacking, $\Delta G$, for the dimer 5'-UU-3'. **(B-J)** Probability distribution of stacking interaction energies, $U_{ST}$, obtained from coarse-grained simulations of UU dinucleotide over the temperature range 212 to 292 K. Configurations with $U_{ST} < -k_B T$ are counted as stacked.

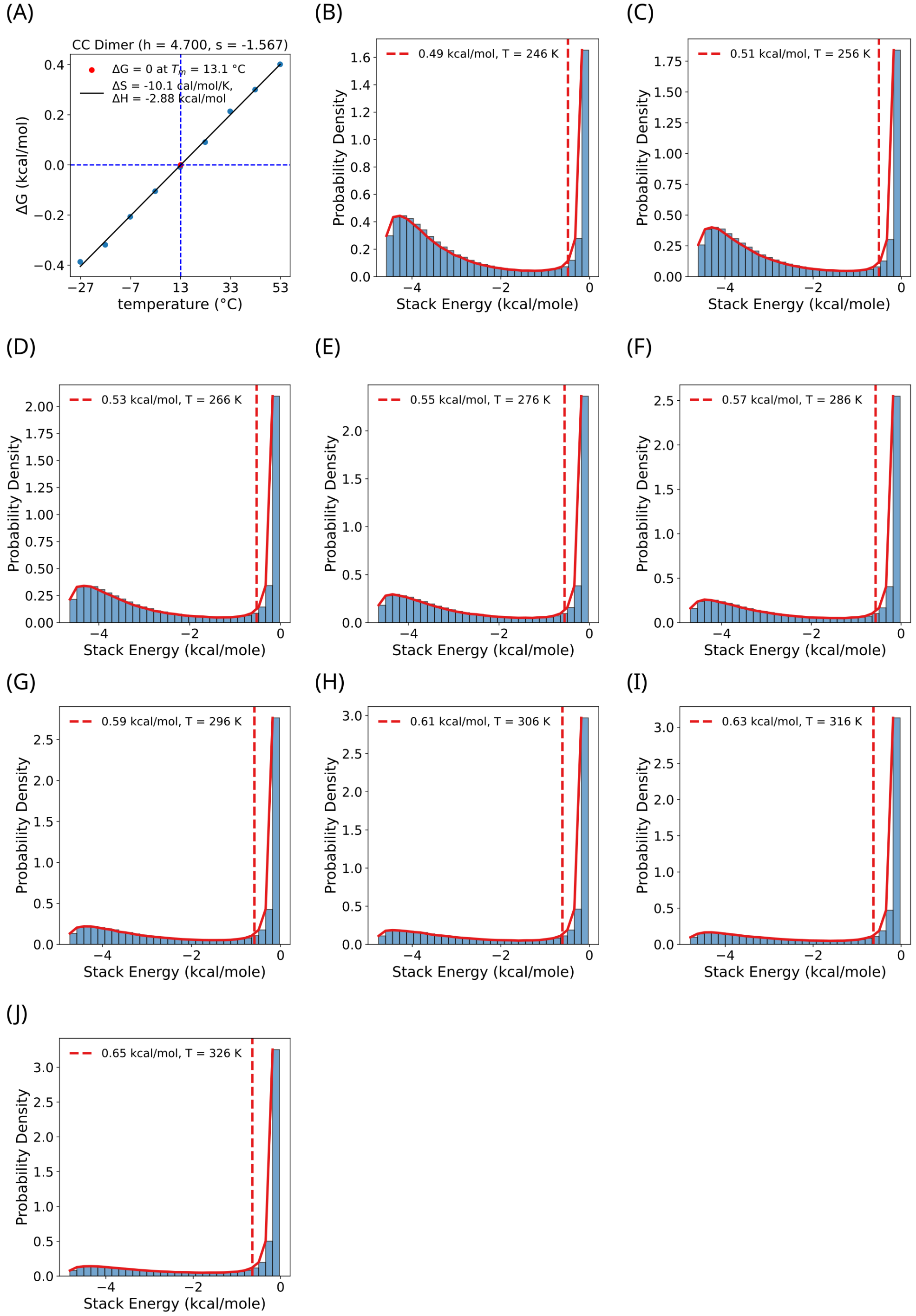

Figure S22: Same as Fig. S21, but for CC.

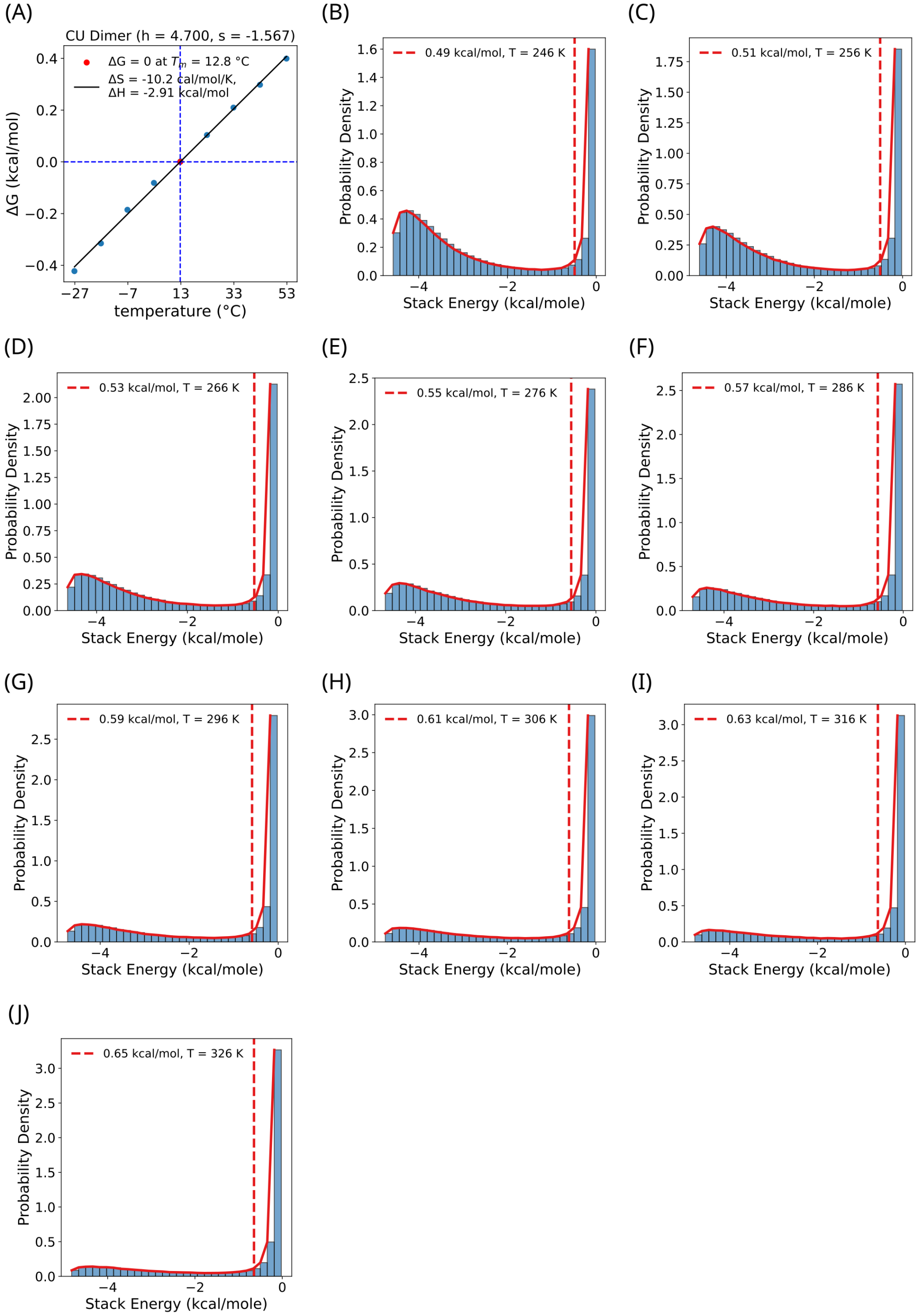

Figure S23: Same as Fig. S21, but for CU.

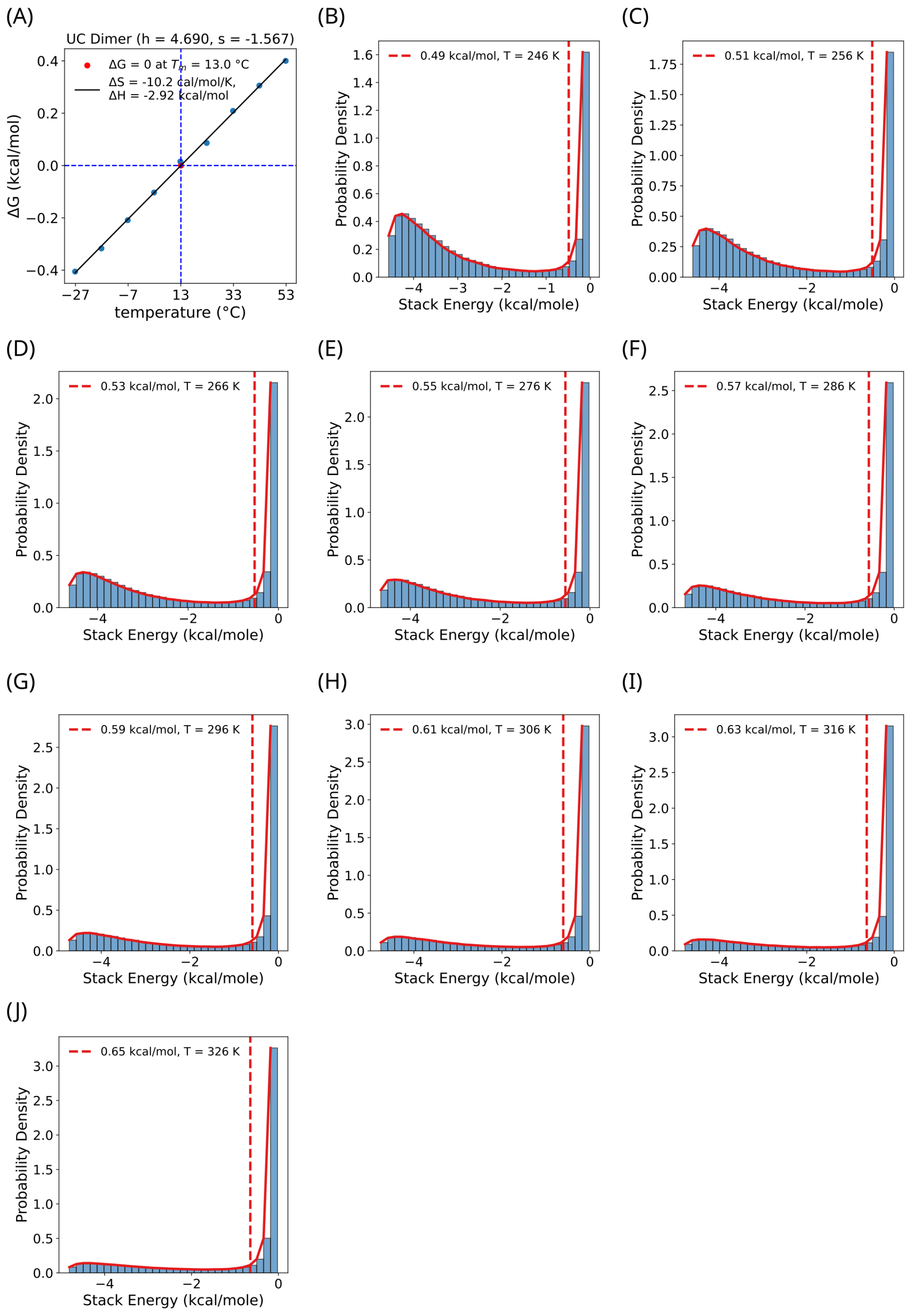

Figure S24: Same as Fig. S21, but for UC.

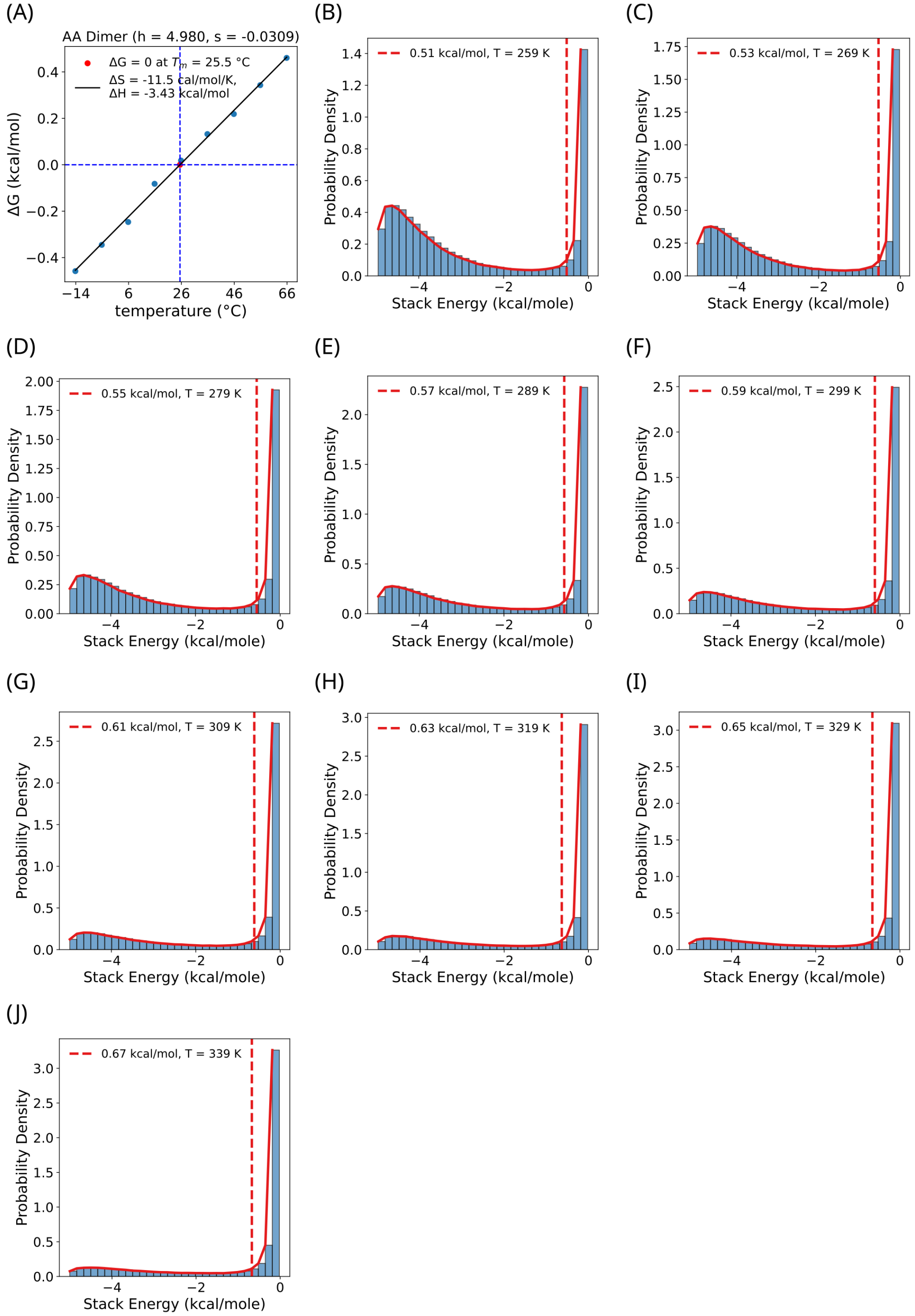

Figure S25: Same as Fig. S21, but for AA.

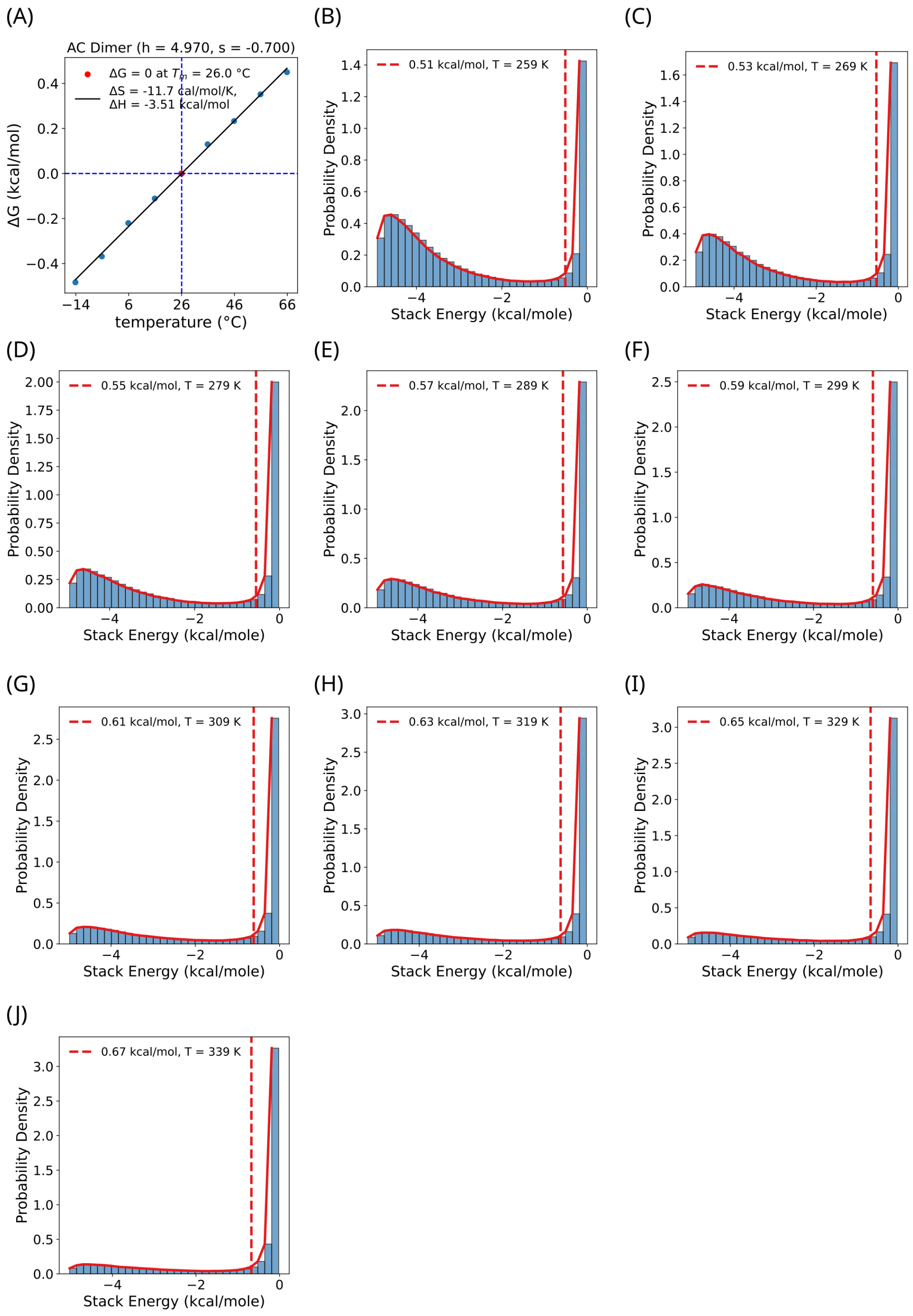

Figure S26: Same as Fig. S21, but for AC.

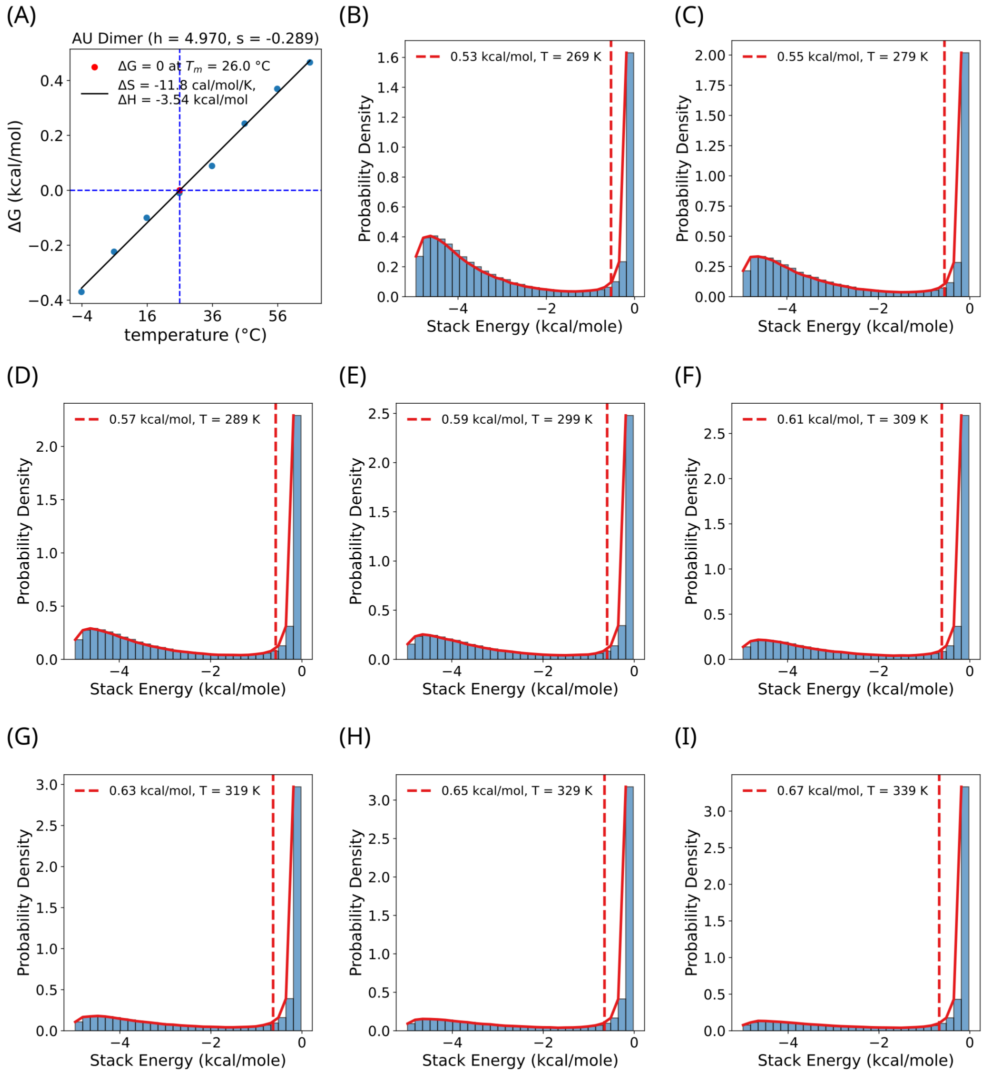

Figure S27: Same as Fig. S21, but for AU.

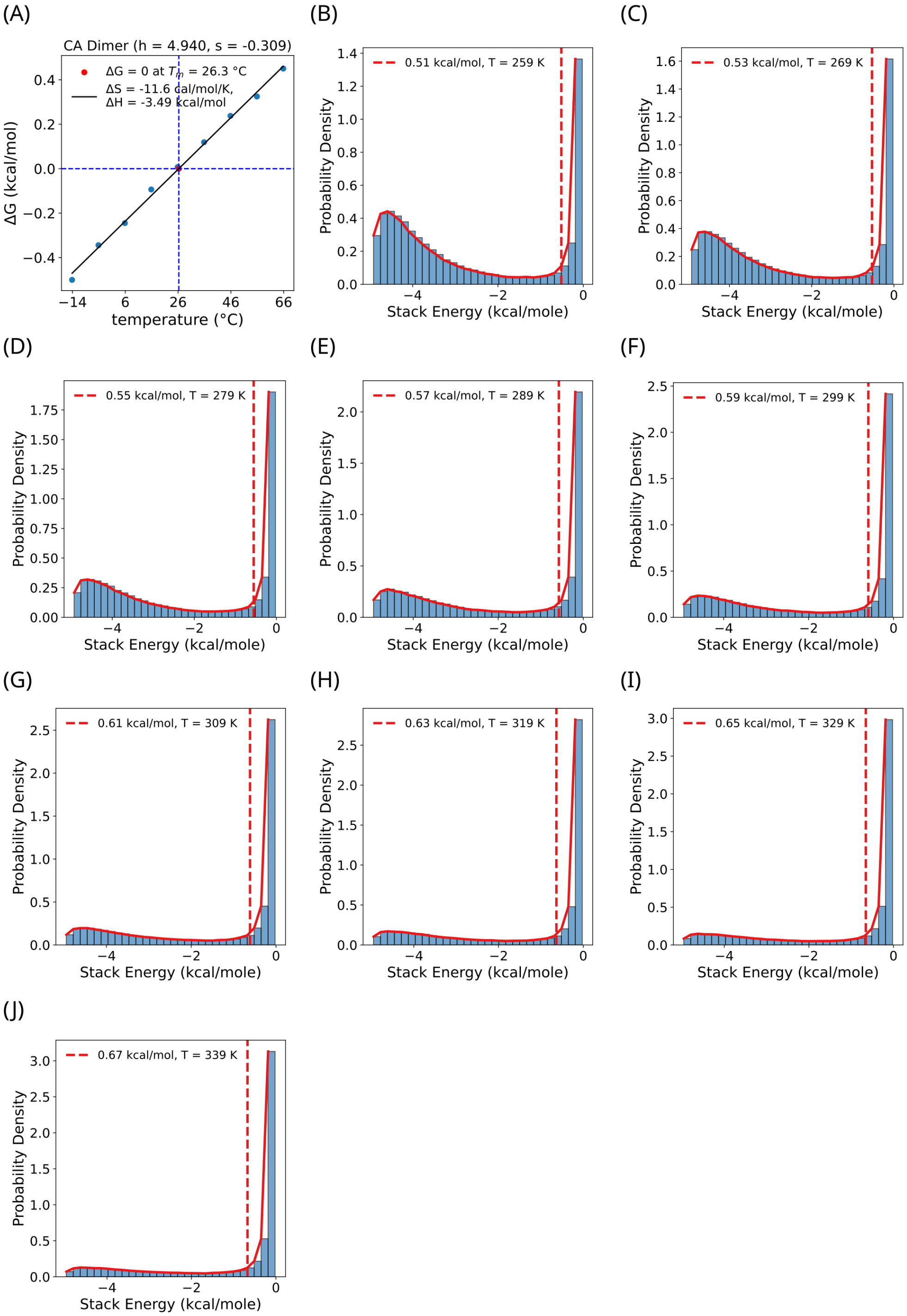

Figure S28: Same as Fig. S21, but for CA.

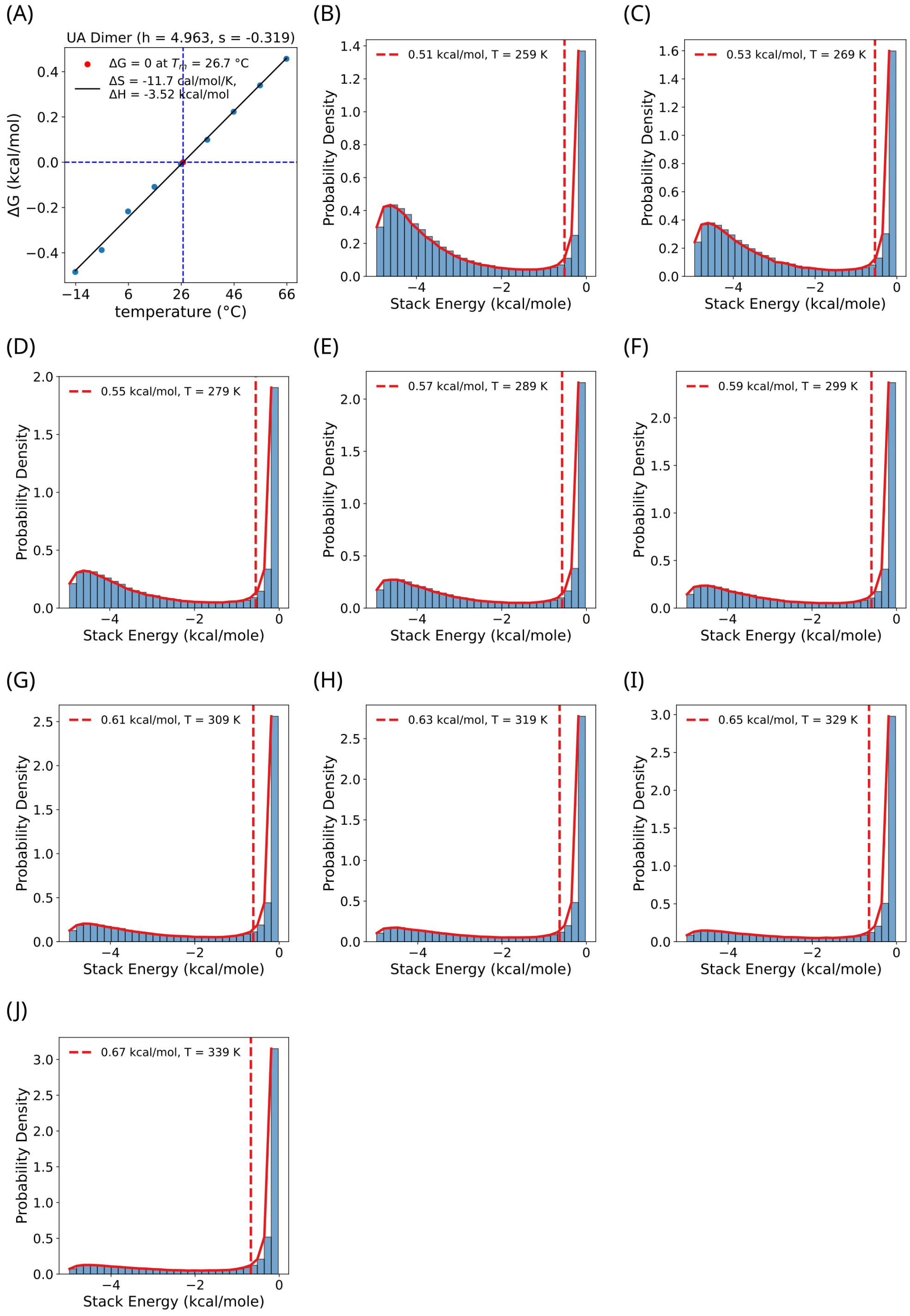

Figure S29: Same as Fig. S21, but for UA.

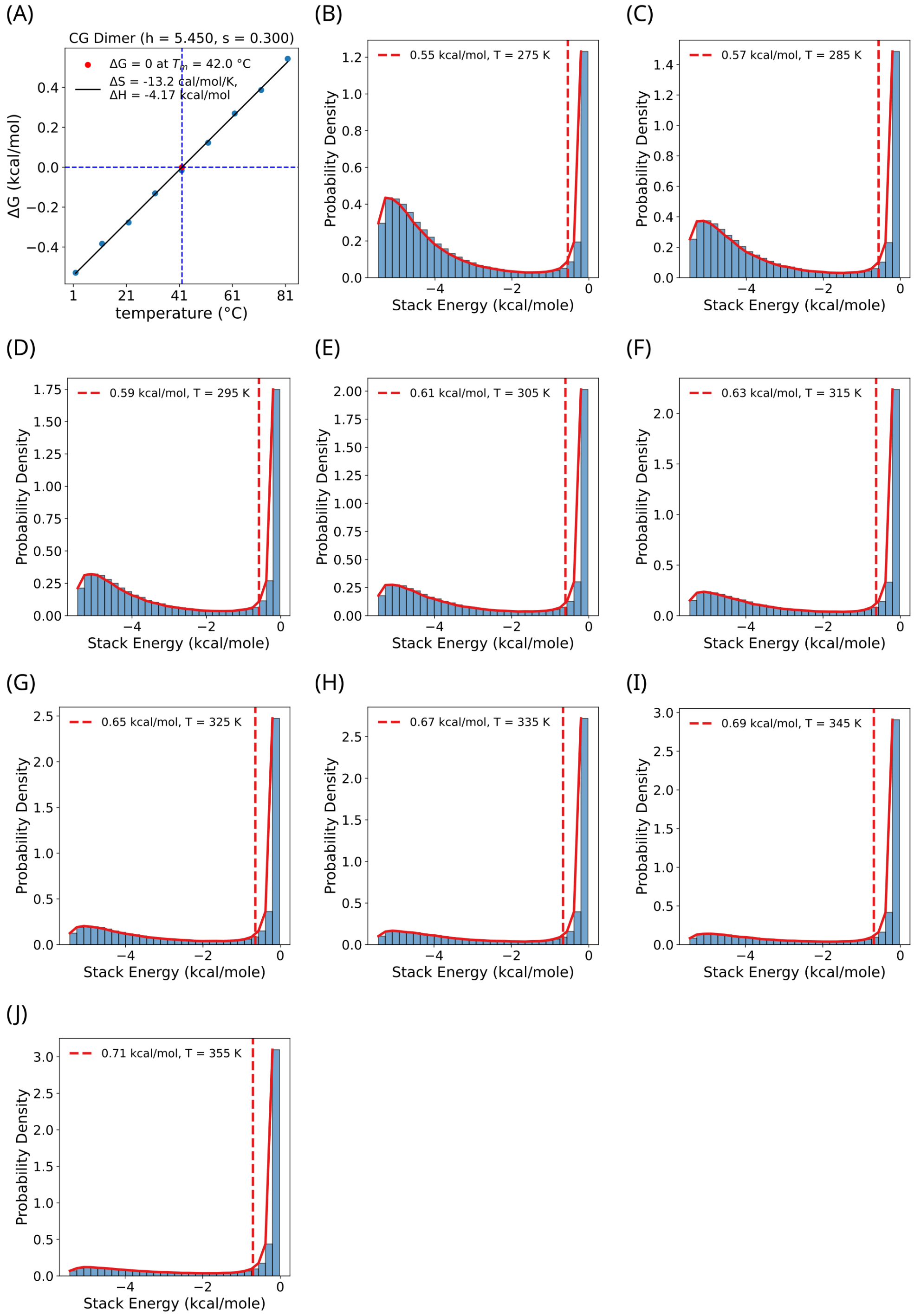

Figure S30: Same as Fig. S21, but for CG.

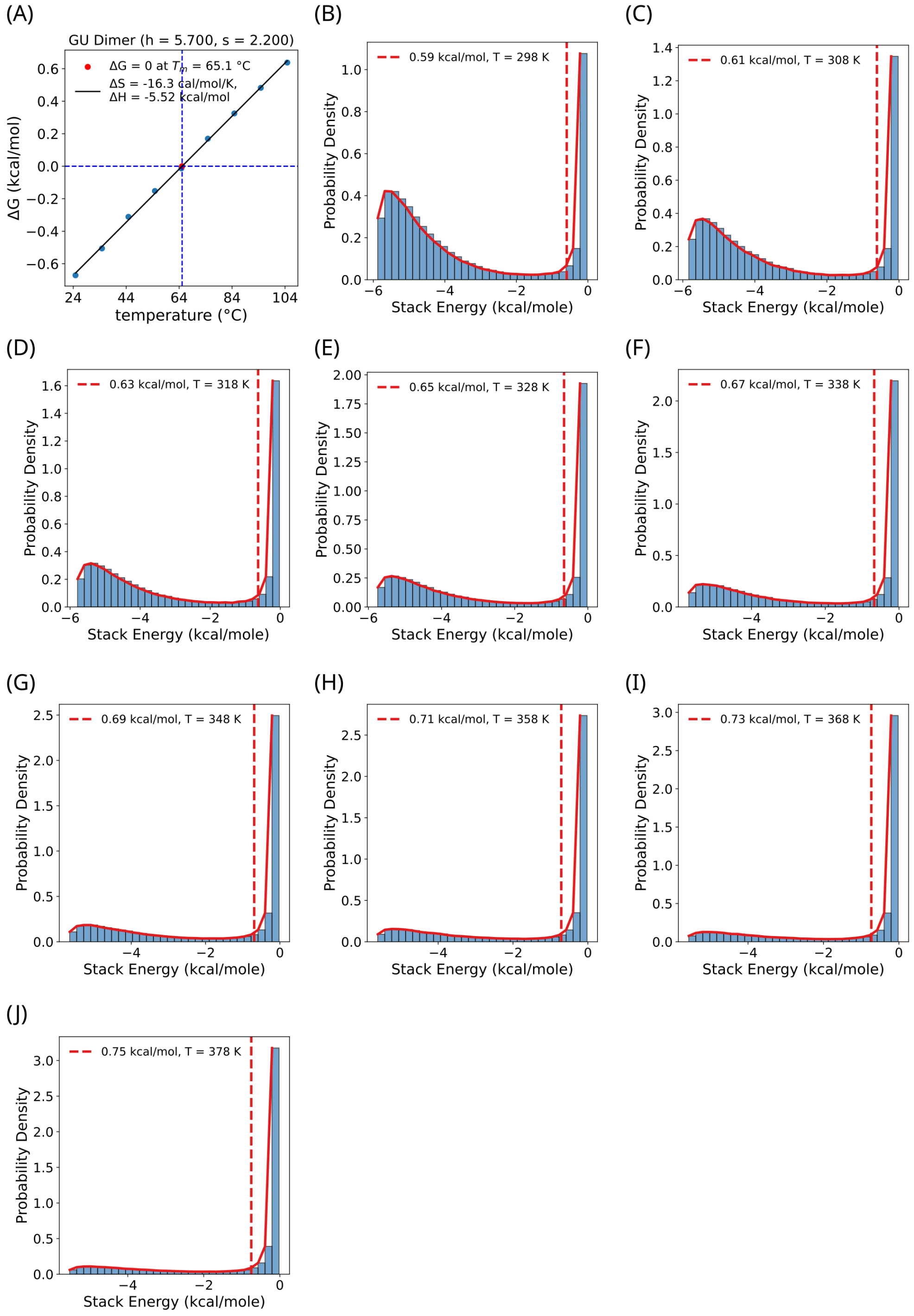

Figure S31: Same as Fig. S21, but for GU.

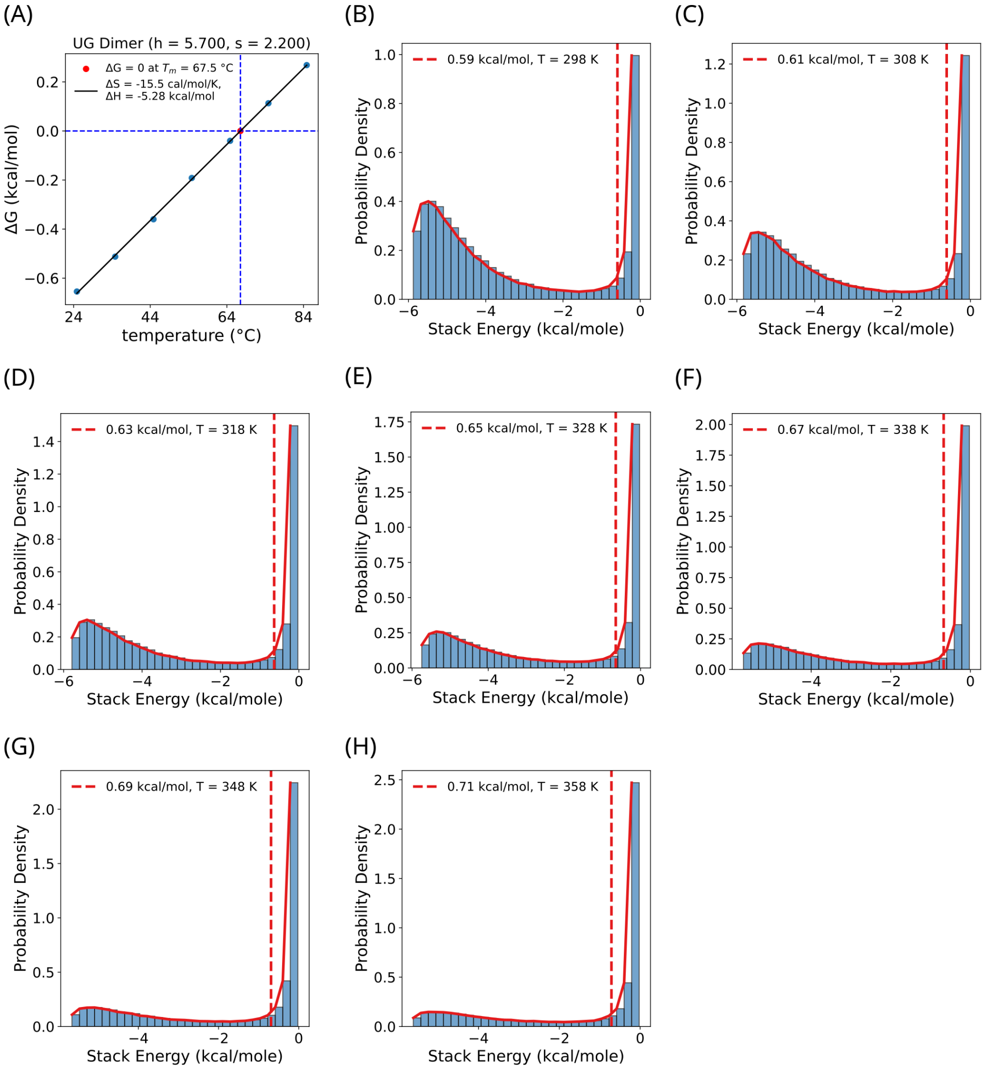

Figure S32: Same as Fig. S21, but for UG.

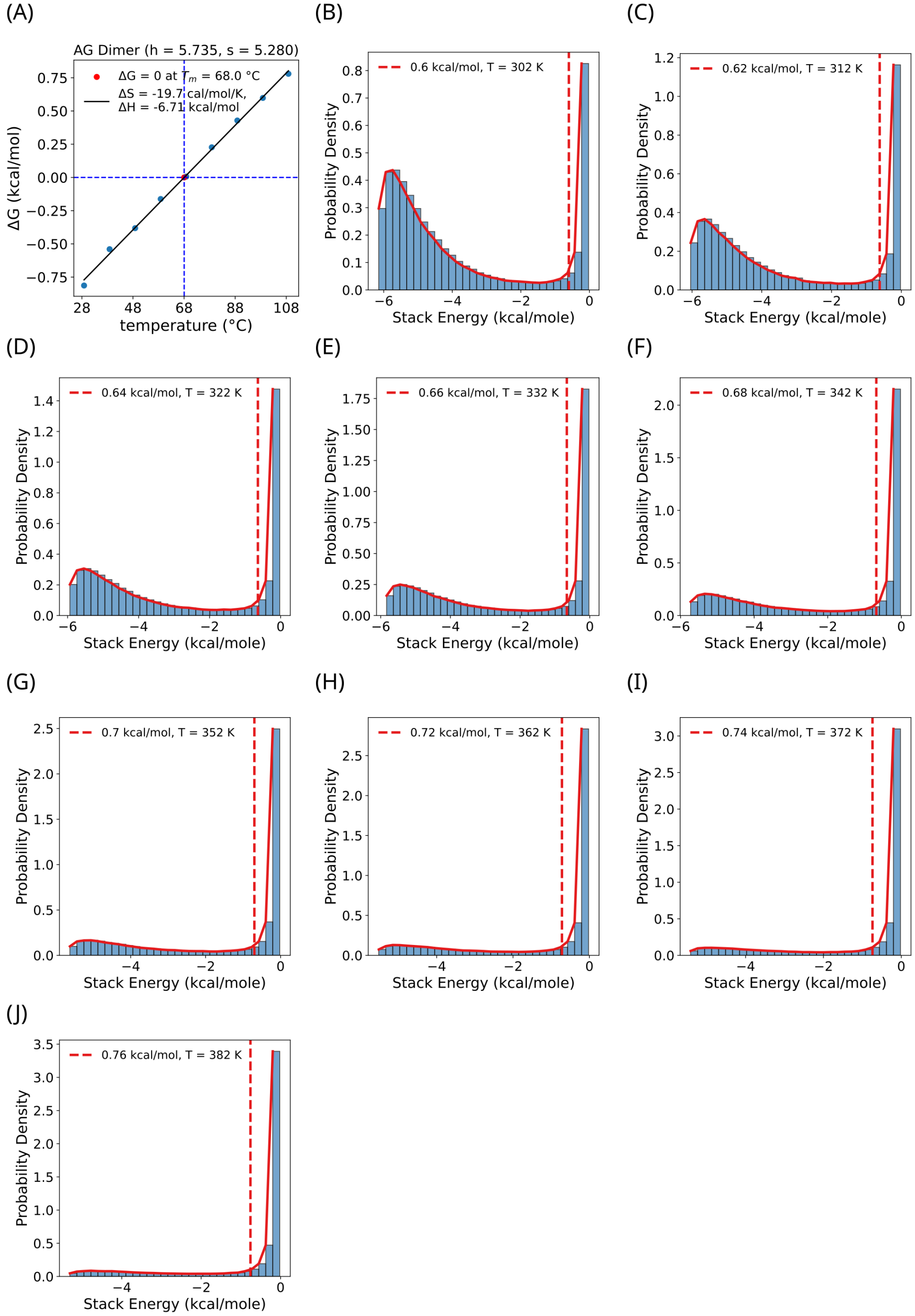

Figure S33: Same as Fig. S21, but for AG.

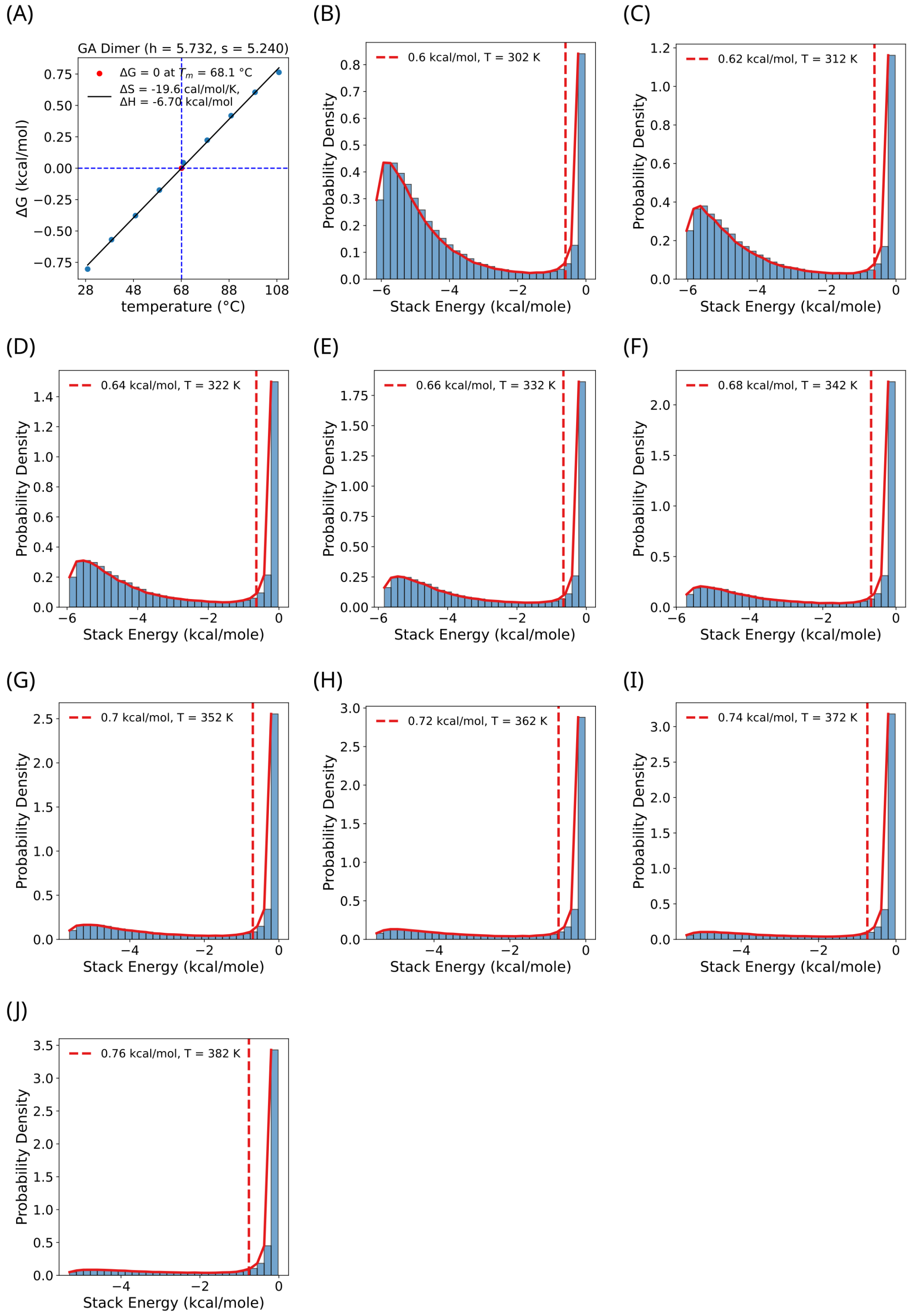

Figure S34: Same as Fig. S21, but for GA.

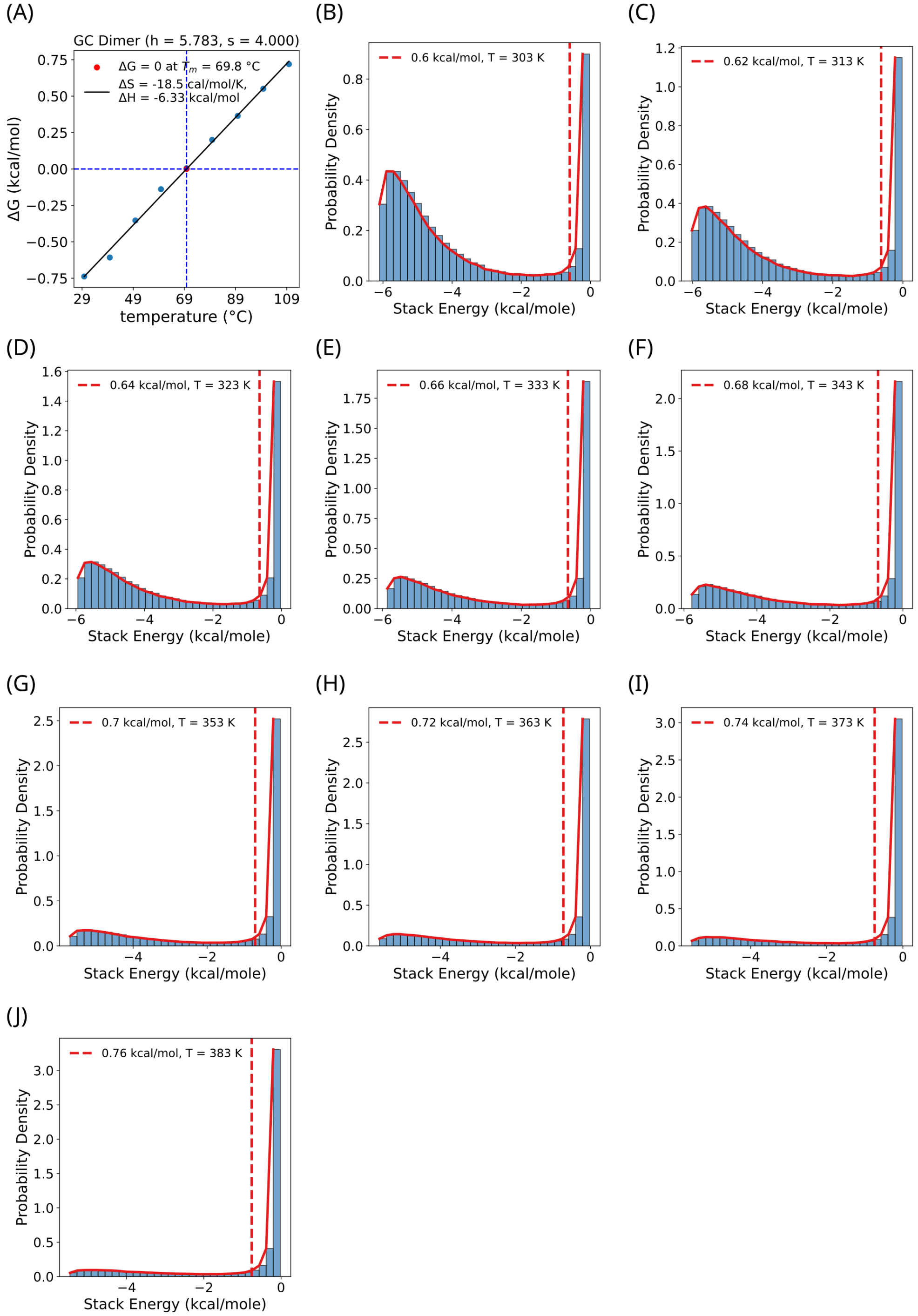

Figure S35: Same as Fig. S21, but for GC.

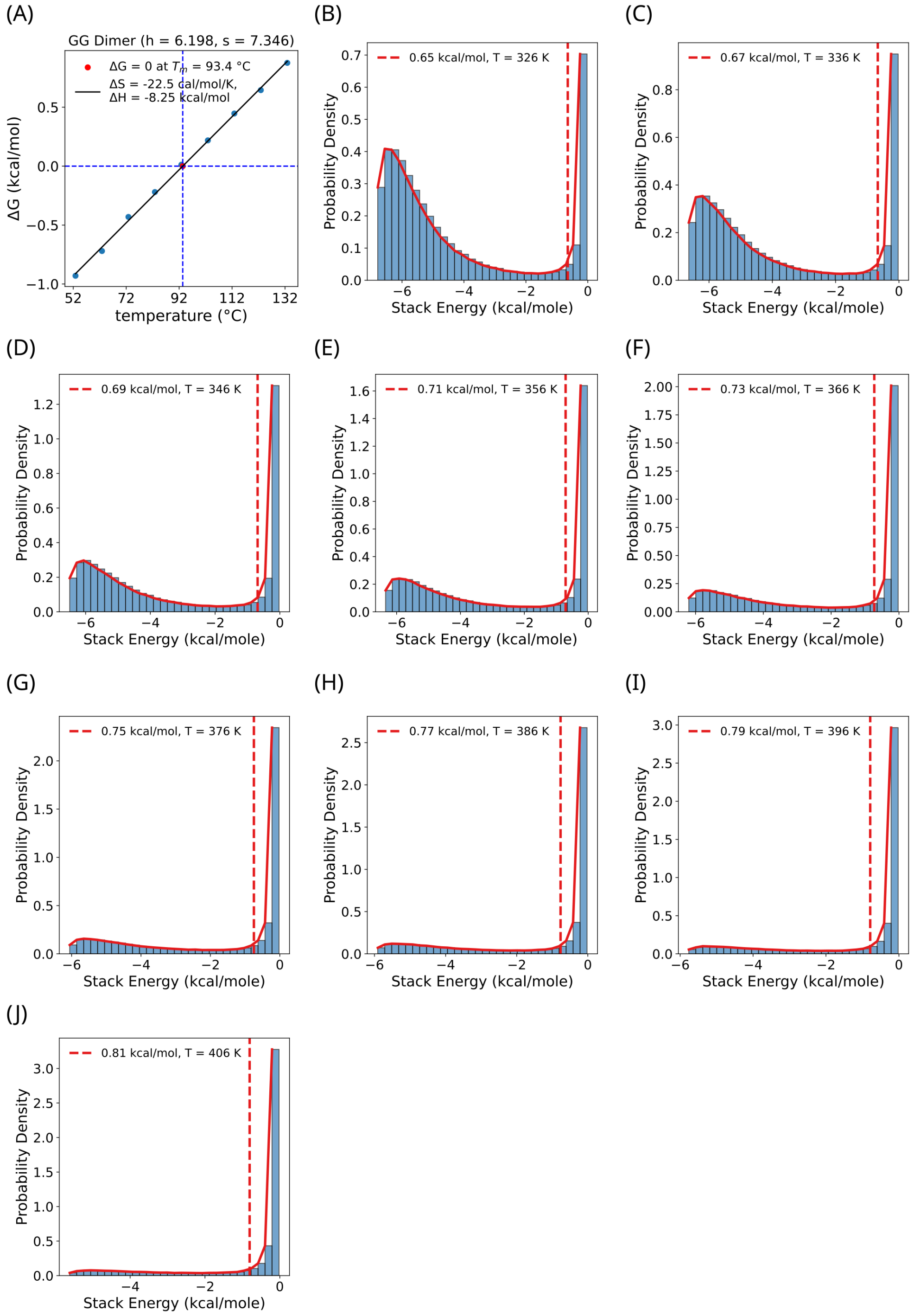

Figure S36: Same as Fig. S21, but for GG.

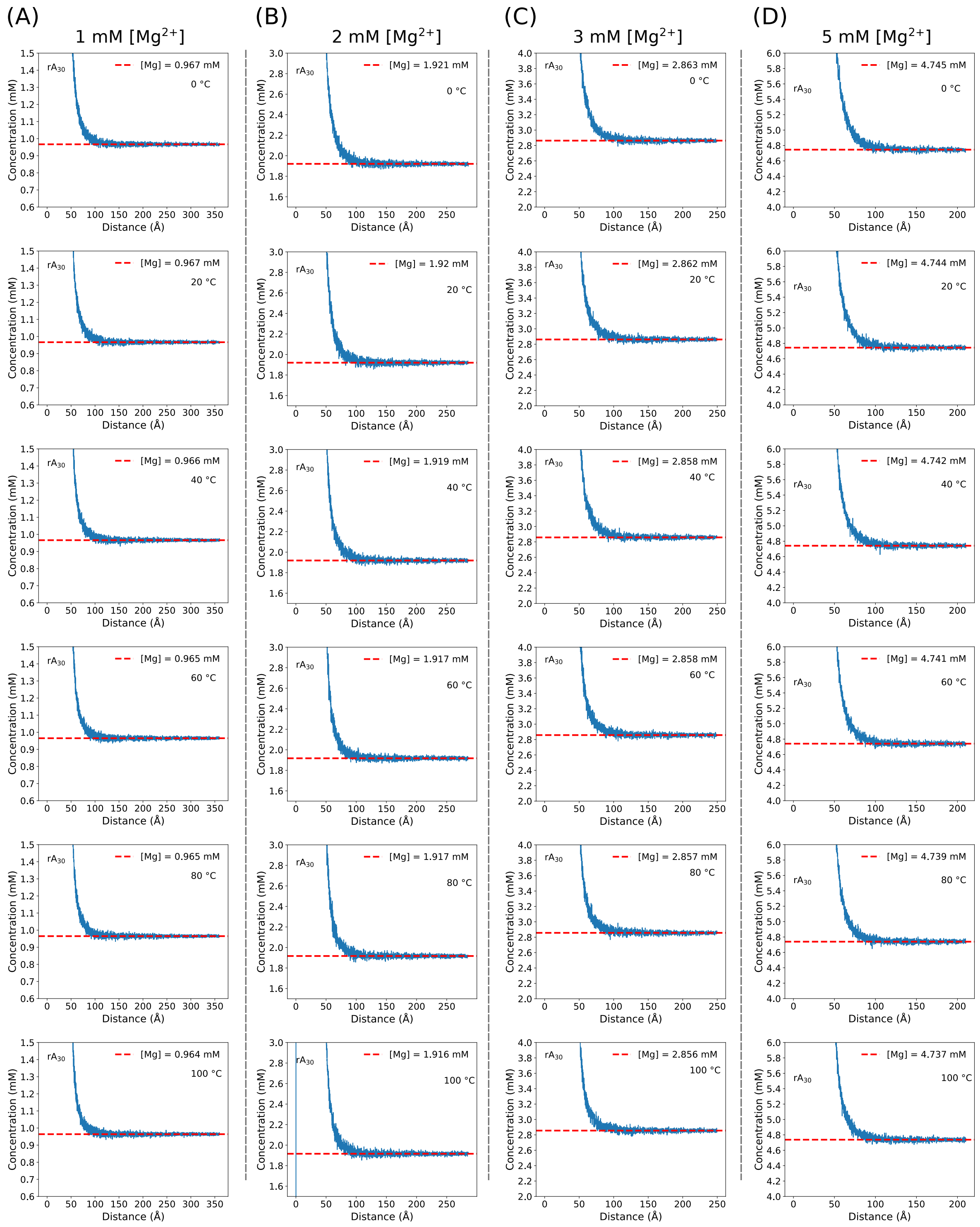

Figure S37: **Effective $Mg^{2+}$ concentration around $rA_{30}$.** Effective local [$Mg^{2+}$] around $rA_{30}$ from the ssRNA COM at temperatures ranging from 0 °C to 100 °C. Each panel corresponds to a different bulk input [$Mg^{2+}$]: (A) 1 mM, (B) 2 mM, (C) 3 mM, (D) 5 mM.

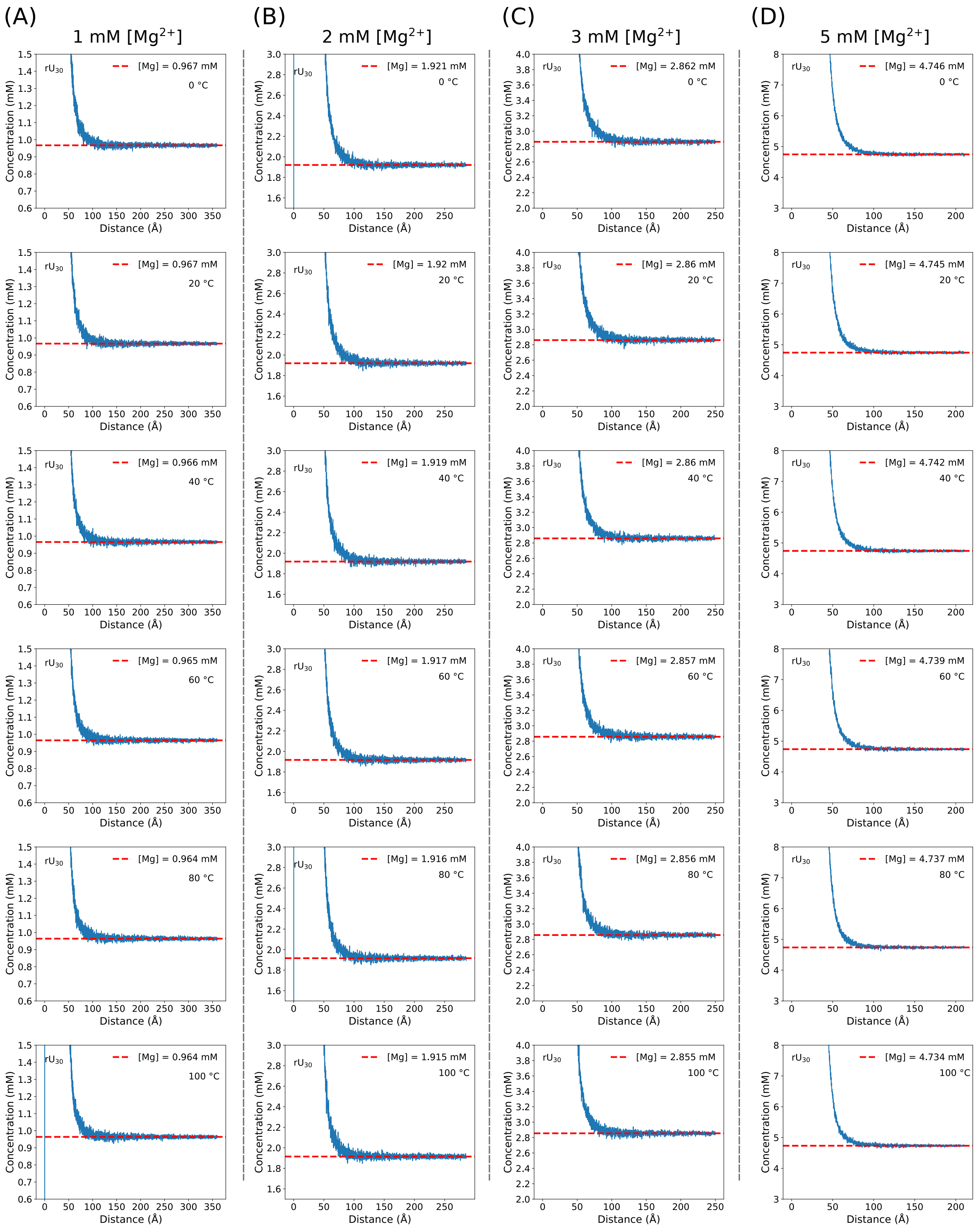

Figure S38: Same as Fig. S37, but for $rU_{30}$.

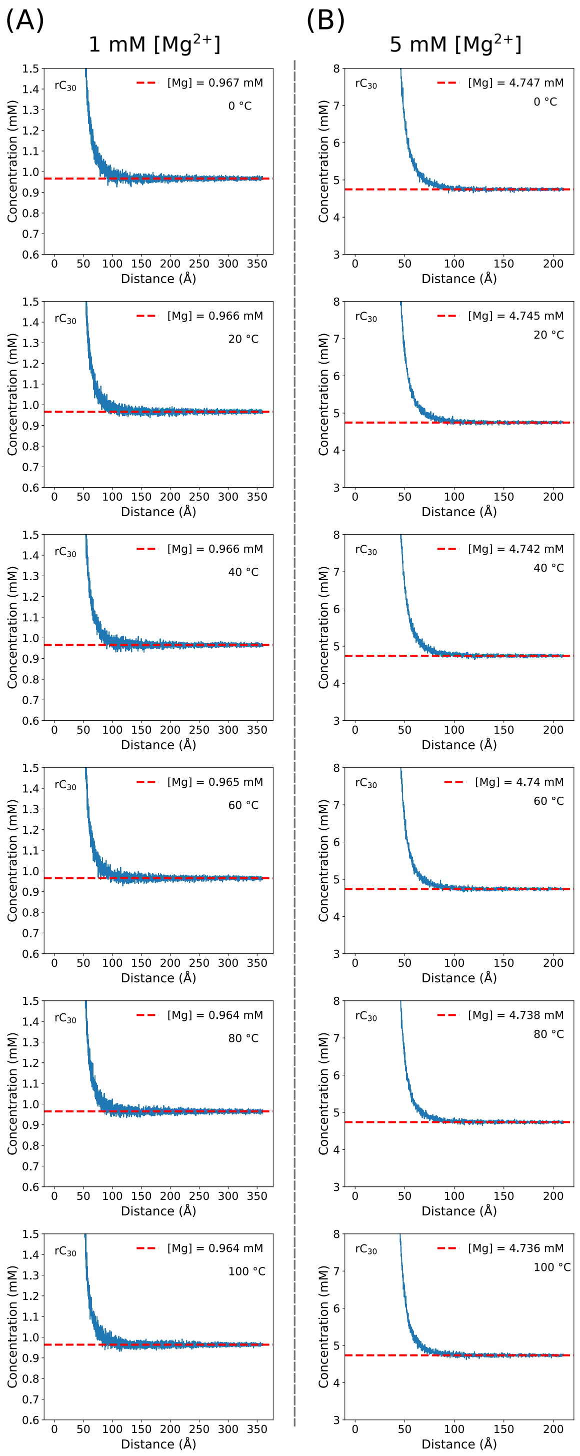

Figure S39: Same as Fig. S37, but for $rC_{30}$.

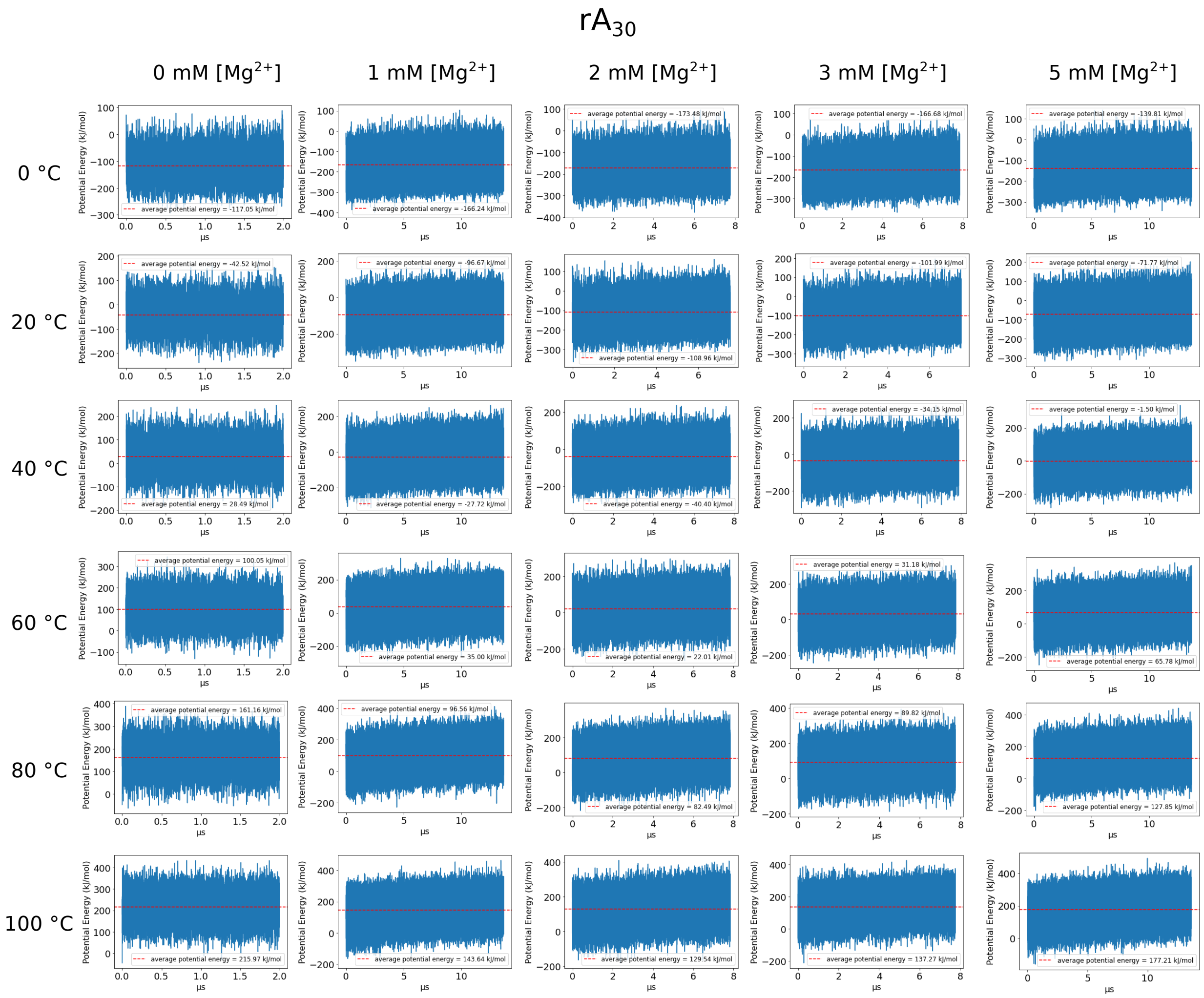

Figure S40: Potential energy profiles of $rA_{30}$ trajectories.

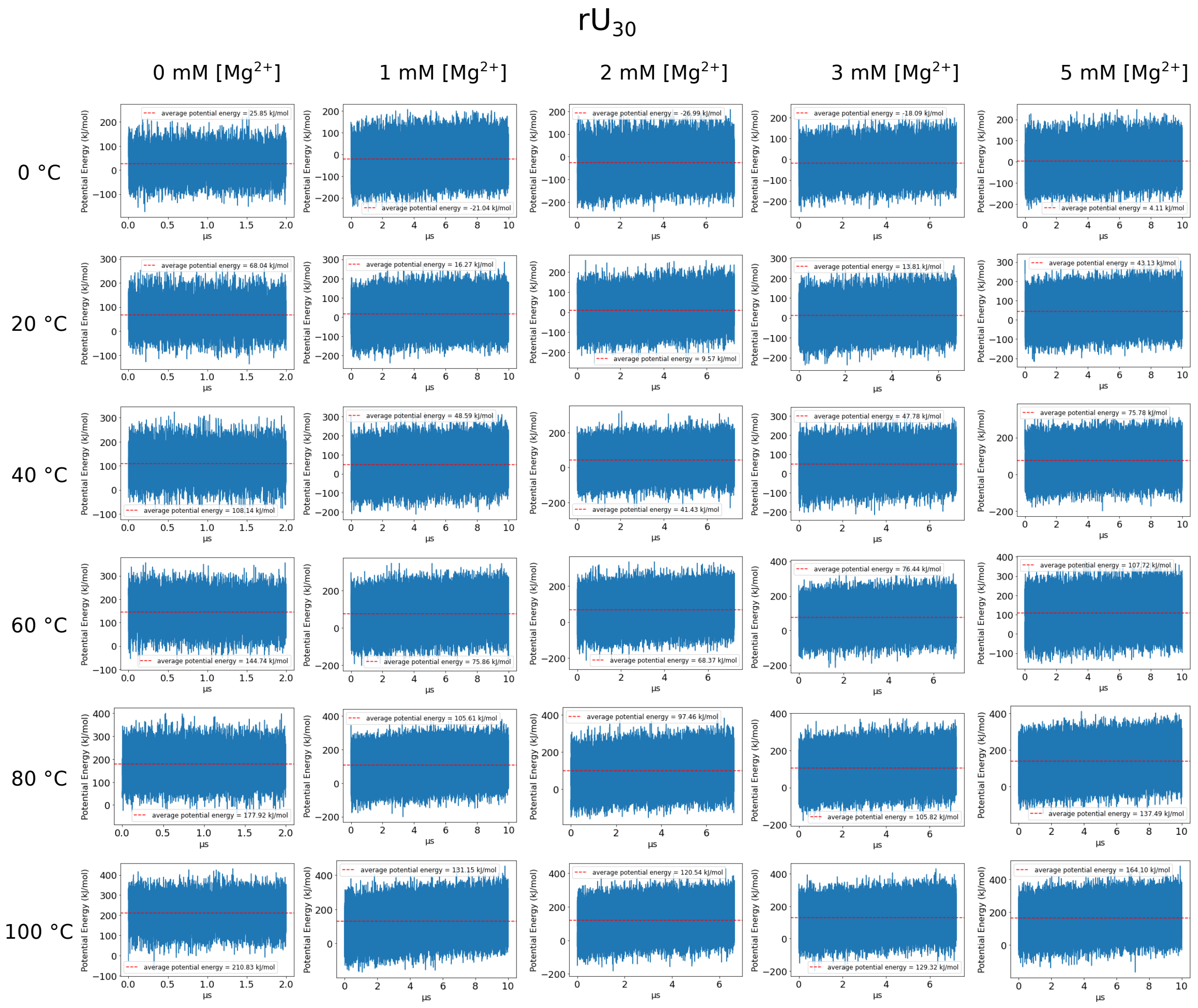

Figure S41: Potential energy profiles of $rU_{30}$ trajectories.

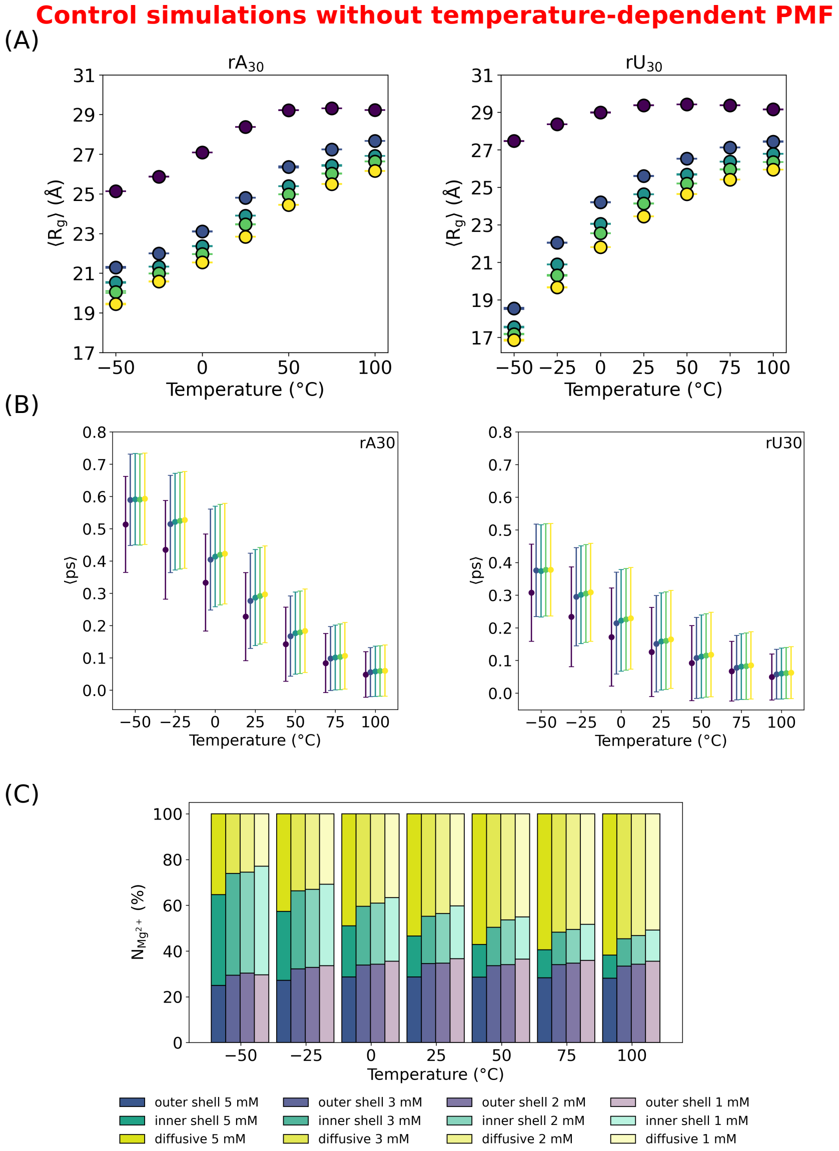

Figure S42: **(A)** Radius of gyration $R_g$ as a function of temperature for $rA_{30}$ and $rU_{30}$ with the $Mg^{2+}$-phosphate PMF held fixed (temperature-independent). In contrast to the full model, no collapse transition is observed upon heating in the presence of $Mg^{2+}$. **(B)** Stacking propensity as a function of temperature shows expected monotonic decrease, indicating that stacking alone does not drive collapse under these conditions. **(C)** Distribution of $Mg^{2+}$ binding mode (inner-sphere, outer-sphere, diffusive populations) as a function of temperature. In the absence of temperature-dependent ion interactions, the characteristic shift from diffusive to inner-sphere coordination is not observed; instead, the diffusive ion population increases with temperature. These results demonstrate that the temperature dependence of the $Mg^{2+}$-phosphate interactions is essential for driving RNA collapse and associated ion distribution.

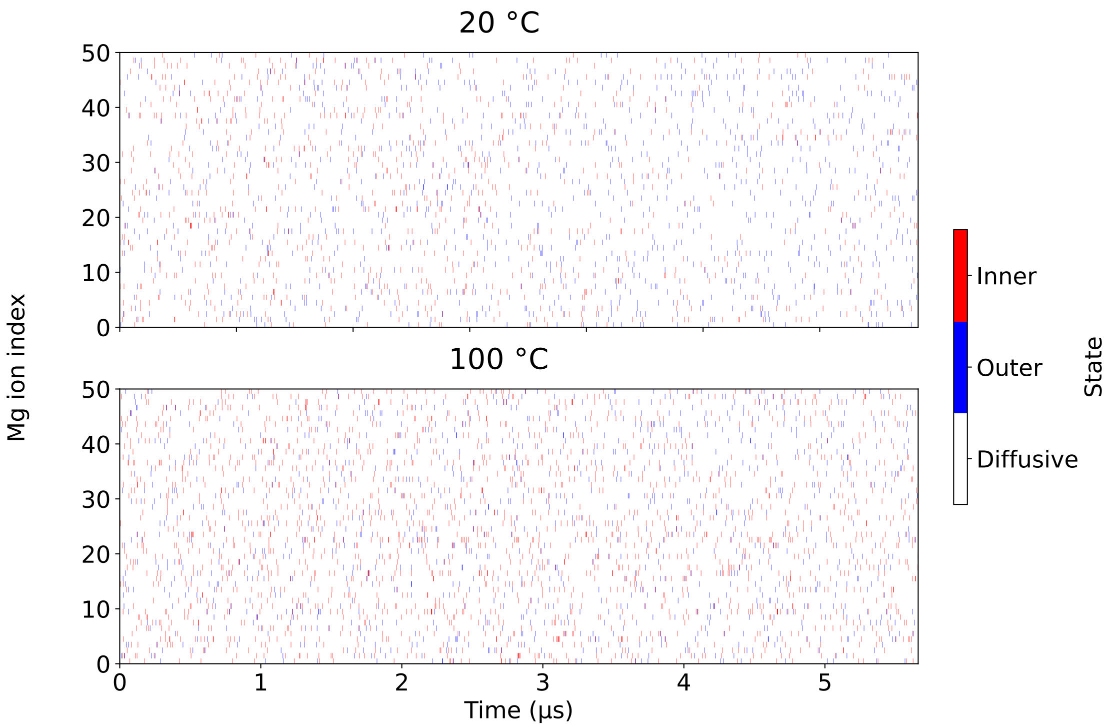

Figure S43: **Time evolution of $Mg^{2+}$ binding states.** Binding states of a representative subset of 50 $Mg^{2+}$ ions shown as a function of time at 20 °C (top) and 100 °C (bottom). Inner-sphere, outer-sphere, and diffusive states are colored red, blue, and white, respectively. Frequent transitions between states are observed throughout the trajectories, indicating that $Mg^{2+}$ ions dynamically exchange between coordination modes and are not kinetically trapped in any binding configuration.

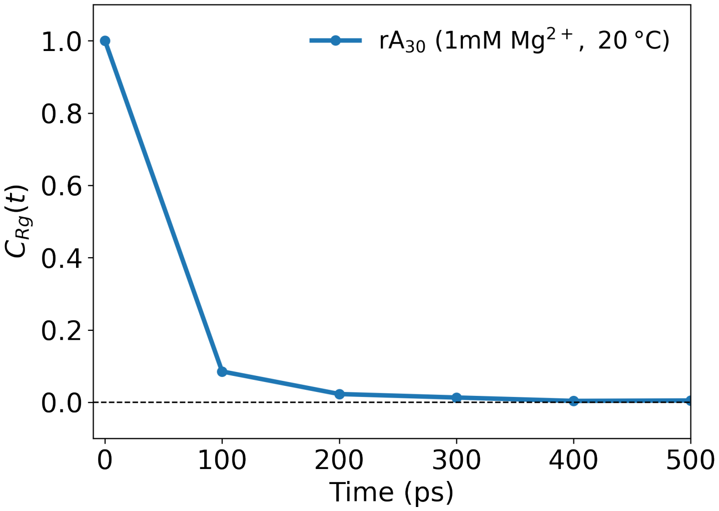

Figure S44: **Autocorrelation analysis of simulation observables.** Normalized autocorrelation function (Eq. 1) $C_{Rg}(t)$ of $R_g$ for a representative system ($rA_{30}$ in 1 mM $Mg^{2+}$ at 20 °C). Configurations were saved every 20 ps, and analyses were performed every 5 frames. The autocorrelation decays rapidly and approaches zero within $\approx$ 400 ps, indicating loss of memory and statistical independence beyond this timescale. The estimated correlation time is therefore much shorter than the block size used for statistical analysis (100 ns), justifying the use of 5,000-frame blocks as effectively independent samples.
